# Extensive nuclear datasets resolve the phylogeny of siphonous green algae and identify genome duplications as a contributing factor to evolutionary adaptations

**DOI:** 10.64898/2025.12.12.693860

**Authors:** Riyad Hossen, Saelin Bjornson, Trevor Bringloe, Heroen Verbruggen

## Abstract

The Bryopsidales, a group of siphonous green algae, are of particular evolutionary interest due to their unusual cellular organization and striking morphological and ecological diversity. There are indications for genome duplication, but low taxon sampling has limited our insights of these processes and how they may contribute to these traits. The relationships among certain bryopsidalean lineages remain unresolved, even with plastid genome-scale datasets. Nuclear genomes offer promise for resolving phylogenetic uncertainties and pinpointing duplication events, but progress has been hindered by the limited availability of such datasets. Here, we present new nuclear genome data for 44 taxa sampled across the phylogenetic breadth of Bryopsidales, and conduct phylogenomic analyses of 708 nuclear genes with coalescent and concatenation approaches. Our results significantly advance the resolution of Bryopsidales relationships, including confident placement of previously hard-to-resolve lineages like *Pseudobryopsis*, Ostreobineae and the Halimedineae tribes. We identified many ancient gene duplications across the Bryopsidales tree, including potential whole genome duplications in the Ostreobiaceae and Caulerpaceae. Our work presents the most highly-resolved phylogeny of Bryopsidales to date and offers an extensive framework for the exploration of the potential roles of genome duplications that may have facilitated niche adaptations and invasive trait development.

## Introduction

Nuclear gene data has become a cornerstone of phylogenetic inference in land plants, and angiosperms in particular, with genomes already available for many species across the angiosperm tree of life and biodiversity-focused projects using target capture filling in gaps rapidly (e.g. Johnson *et al*., 2019). The Chlorophyta, the algal sister lineage of the Streptophyta that gave rise to the land plants, have seen much less work of this nature and, even though deeper-level phylogenetics to infer relationships among phyla have been addressed (Jackson *et al*., 2018; Del Cortona *et al*., 2020; Li *et al*., 2021), there has been less work to fill in the biodiversity within the major groups.

The siphonous green algae comprise ca. 564 species of green seaweeds in the order Bryopsidales, predominantly occurring in tropical and temperate marine environments (Kerswell, 2006; Guiry & Guiry, 2024). Their thallus (algal body) shows a siphonous structure, meaning that it is composed of a single large tubular cell, wherein numerous nuclei and chloroplasts move around via cytoplasmic streaming (Vroom & Smith, 2003). They exhibit a wide range of morphologies, from simple microscopic forms just a millimeter tall to highly complex macroscopic thallus structures that can become several meters tall and can form reefs and other structures (Huisman, 2015; Huisman & Verbruggen, 2015).

The order includes some iconic seaweed genera (e.g. *Caulerpa*, *Codium*, *Halimeda*) that play significant ecological roles as primary producers, habitat constructors, and calcium carbonate producers in coral reefs, rocky shores, lagoons, and seagrass beds (Bulleri *et al*., 2006; Bulleri *et al*., 2010; Castro-Sanguino *et al*., 2020). Some genera, like *Caulerpa*, are infamously invasive (Williams & Grosholz, 2002) and some are used as food and medicine (de; Meinita *et al*., 2022; Pan-Utai *et al*., 2023). Moreover, the group serves as a model in research on algal evolution (Cremen *et al*., 2018; Ochiai *et al*., 2024), cell biology (Ranjan *et al*., 2015; Arimoto *et al*., 2019), ecophysiology (Kim *et al*., 2001; Ukabi *et al*., 2013; Yildiz & Dere, 2015), biogeography (Verbruggen *et al*., 2005; Varela-Álvarez *et al*., 2015), climatic niche evolution (Verbruggen *et al*., 2009b), and coral reef biology (Iha *et al*., 2021; Galindo-Martínez *et al*., 2022).

Due to their ecological importance and diverse applications, the phylogenetic relationships of siphonous algae have been studied in some detail. Early molecular phylogenetic studies that primarily relied on single markers (e.g. chloroplast *rbc*L or nuclear ribosomal 18S rDNA) have provided a basic blueprint of the higher-level structure of the group, including the recognition and refinement of the two main suborders Bryopsidineae and Halimedineae (Woolcott *et al*., 2000; Lam & Zechman, 2006; Curtis *et al*., 2008), their families (Kooistra, 2002; Lagourgue & Payri, 2022), and genera (Verbruggen *et al*., 2007; Kazi *et al*., 2013; Draisma *et al*., 2014). These marker gene-based studies were limited in resolution, with several taxa not placed confidently. Later work based on multiple markers and whole plastomes further refined phylogenetic relationships (Cremen *et al*., 2019), provided a time-scale of the group’s diversification (Verbruggen *et al*., 2009a) and defined the suborder Ostreobineae for the genus *Ostreobium* (Verbruggen *et al*., 2017), an under-studied group of limestone-boring algae (Tandon *et al*., 2023).

While this work provides a sound basis, several phylogenetic relationships have remained unresolved, including the relationships among the ecologically dominant families and tribes in the Halimedineae, as well as the placement of the genera *Pseudobryopsis* and *Ostreobium*. In the Halimedineae suborder, the relationships of Udoteaceae, Rhipiliaceae, and Halimedaceae have shown inconsistencies across studies and, generally speaking, low statistical support (Lam & Zechman, 2006; Curtis *et al*., 2008; Verbruggen *et al*., 2009a; Cremen *et al*., 2019). Similarly, *Pseudobryopsis* in Bryopsidineae has traditionally been considered related to Bryopsidaceae and Derbesiaceae based on life history and morphologies (Hillis-Colinvaux, 1984) but molecular studies have recovered it (or the related genus *Trichosolen*) in a range of different positions, leaving its position uncertain (Lam & Zechman, 2006; Curtis *et al*., 2008; Cremen *et al*., 2019). The genus *Ostreobium* has similarly proven difficult to place, with some studies assigning it to Bryopsidineae or yielding inconclusive results (Vroom *et al*., 1998; Woolcott *et al*., 2000; Curtis *et al*., 2008) and others suggesting it is an early-branching lineage (Verbruggen *et al*., 2017; Cremen *et al*., 2019). To address these unresolved questions in Bryopsidales phylogenetics, several of which could not be resolved even with whole plastomes, it will be crucial to access new data sources, including additional taxon sampling in the difficult-to-place taxa, and large numbers of genes to obtain sound resolution and understand sources of conflict. Nuclear-encoded protein-coding sequences can greatly facilitate investigations into phylogenetics and evolutionary processes and have helped to resolve many questions (Del Cortona *et al*., 2020; Hou *et al*., 2022).

Duplications of genes and whole genomes (WGD) are considered major drivers of plant evolution that contribute genomic innovation and functional novelty (Ohno, 1970; van de Peer *et al*., 2009; Schranz *et al*., 2012; Rensing, 2014). Gene duplication can facilitate adaptation and support diverse lifestyles, particularly in new or extreme environments, by enhancing fitness through increased gene dosage, dividing ancestral functions to optimize specific tasks (subfunctionalization), or acquiring new functions (neofunctionalization) (Ohno, 1970; Kondrashov, 2012; Magadum *et al*., 2013; Zhang *et al*., 2023; Cho *et al*., 2023). Duplication events, including WGDs and polyploidization by hybridization, can enable similar innovations to occur at large-scale, although it can have a range of outcomes (Charron *et al*., 2019; Carretero-Paulet & van de Peer, 2020; Bowers & Paterson, 2021). Many genes resulting from duplications are subsequently lost (van Hoek & Hogeweg, 2009; Panchy *et al*., 2016; Levin, 2019), and the adaptive potential of duplications may be most beneficial under periods of environmental change (Cuypers and Hogeweg 2014; Carretero-Paulet and van de Peer 2020).

WGD are common in land plants, particularly in angiosperms where they have been linked to evolutionary innovations in complexity, novel morphologies, metabolic diversification, and adaptation to new habitats (Wu *et al*., 2020; Leslie & Mander, 2025). However, very little information is available about WGD in the Chlorophyta (One Thousand Plant Transcriptomes Initiative, 2019). A recent genome study of the important CaCO3 reef-forming species *Halimeda opuntia* (L.) Lamouroux suggested that three intraspecific polyploidisation events had occurred, contributing to the diversification of calcification-related genes (Zhang et al, 2024). Given the peculiar adaptations and extreme vigor seen in other Bryopsidales, such as for example endolithic lifestyles in Ostreobiaceae and highly invasive characteristics in *Caulerpa*, it is possible that gene or genome duplications may have contributed to the evolutionary and ecological success of more taxa in this order. However, genome duplication and its wider evolutionary implications across Bryopsidales remains largely unexplored.

To date, large datasets of nuclear-encoded sequences are only available for eight bryopsidalean species and these datasets have not been used to evaluate within-order phylogenetic relationships or gene duplication events. Here, we generated and analyzed a large dataset including 44 new draft nuclear genomes and transcriptomes with the objective to reassess lineage relationships of Bryopsidales, and use this new species tree to identify ancient nodes with considerable gene duplication events suggestive of WGD, and investigate the functions of these duplicated genes.

## Materials and methods

### Taxon sampling

We sampled 71 taxa for our study, including 44 newly sequenced and 27 from publicly available databases (Table S1). This included 3 taxa from Ostreobineae, 11 from Bryopsidineae, and 37 from the Halimedineae suborder, with a focus on improving representation in lineages with previously uncertain placements, particularly *Pseudobryopsis* and various Halimedineae tribes (Udoteae, Rhipiliopsideae, Rhipileae, and Pseudocodieae). For outgroups, we selected 6 Dasycladales, 3 Cladophorales, 5 Chlorophyceae, and 6 other Ulvophyceae (1 Oltmansiellopsidales, 1 Ignatiales, and 4 Ulvales), from which various combinations were used in phylogenetic inference. Fresh samples were primarily collected from Heron Island, Great Barrier Reef, along with additional specimens from John A. West’s algal culture collection, and some previously made collections. All raw reads from newly sequenced samples are available in the NCBI SRA database under BioProject PRJNA1216519.

### Nucleic acid extractions and sequencing

For RNA isolation, fresh samples were preserved in RNA*later* Stabilization Solution (Invitrogen) immediately after collection, except for *Halimeda opuntia* and *Derbesia* sp., for which we used live cultures (30 μmol photons/m²/s; 12:12 dark/light cycle; Provasoli enriched seawater media) flash-frozen in liquid nitrogen. Total RNA was extracted using TRIzol (Invitrogen) or Qiagen RNeasy Plant Mini Kit, following the manufacturer’s instructions.

For species we could not collect fresh, we opted to sequence total DNA from ethanol-preserved, herbarium-pressed or silica-gel-preserved materials. DNA was extracted from these preserved samples following an optimised CTAB method (Hossen *et al*., 2025), and Illumina libraries were prepared (VAHTS Universal DNA kit) and sequenced on the Illumina NovaSeq platform, generating 150 bp paired-end reads and aiming for 30x or higher coverage of the nuclear genome.

### Transcriptome and genome assembly and annotation

Sequence reads were quality-checked with FastQC v0.12.1 and cleaned by removing reads with a Phred Score of 20 or below, and reads containing adapters or poly-N sequences using Trimmomatic v0.39. Cleaned RNA reads were de novo assembled with Trinity v.2.8.3 with default parameters (Grabherr *et al*., 2011). Transcripts of each taxon were clustered using CD-HIT with 98% sequence similarity, and the longest transcript in each cluster was selected as a unigene for further analyses (Li & Godzik, 2006). Protein coding sequences (CDS) were predicted with TransDecoder v5.5.0 (Haas *et al*., 2013). DNA reads were assembled using SPAdes v3.15.5 (Bankevich *et al*., 2012), and assemblies were further scaffolded and redundancy-filtered with Redundans (Pryszcz & Gabaldon, 2016).

Our samples were not axenic, and since Bryopsidales often host diverse bacterial communities within their thalli contamination removal was essential (Pushpakumara *et al*., 2023; Hollants *et al*., 2013). The BlobToolKit v2.6.5 (Challis *et al*., 2020) was used to remove contaminating contigs by visualizing coverage determined by samtools (Li *et al*., 2009) and hits to NCBI nt and Uniprot Reference Proteomes databases.

Contigs were softmasked using RepeatMasker v4.1.5 (Smit *et al*., 2013), using de novo repeats found from RepeatModeler v2.0.4 (Flynn *et al*., 2020), as well as Tandem Repeats Finder (Benson, 1999) and WindowMasker (Morgulis *et al*., 2006). RNA reads were mapped to genomes using Hisat v2.2.1 (Kim *et al*., 2019). When RNA of the same species was not available, RNA from the closest available species was used, and mapped with increased sensitivity by --score-min L,0,-1. For CDS prediction, BRAKER1 (Hoff *et al*., 2016), and BRAKER2 (Brůna *et al*., 2021) were both run and transcripts selected with TSEBRA using an intron support of 0.5 (Gabriel *et al*., 2021). In addition, BRAKER3 (Gabriel *et al*., 2024) was run. From these two pipelines, we selected the one producing the most complete BUSCOs (Benchmarking Universal Single-Copy Orthologs) for each genome, as determined with BUSCO v5.4.5 (Seppey *et al*., 2019). For protein alignments in BRAKER, the Viridiplantae_orthoDB v11 database (Kuznetsov *et al*., 2023) was used, and iteratively updated for each genome by adding proteins from each subsequent CDS prediction to increase representation in Bryopsidales.

### Ortholog selection, alignment, and trimming

To obtain nuclear genes for phylogenetic inference, we ran BUSCO v5.4.5, which enables the rapid and reliable selection of lineage-specific nuclear-encoded conserved genes (Zhang *et al*., 2019; Timilsena *et al*., 2022). Based on its homology search algorithm and the Chlorophyta database, BUSCO returned a dataset of 1080 orthologous genes (OGs), which were filtered to have minimum taxon occupancy of 80% (retaining 813 OGs). The CDS of each OG were aligned using MAFFT v7.310 with both nucleotide and translated peptide sequence specific alignment options “--nuc” and “--amino”. Ambiguously aligned regions of each OG were trimmed using TrimAl with a 70% gap penalty, and short OGs (<100 bp) were removed, resulting in a final dataset of 708 OGs for downstream analyses.

### Paralog filtering

To address cases where certain taxa had multiple sequences within BUSCO OGs, we applied a systematic paralog filtering approach to retain a single representative sequence per taxon for each OG for downstream phylogenetic inference. We first constructed gene trees of the unfiltered OGs using maximum likelihood (ML) inference in IQ-TREE v2.2.2.6 with 100 standard bootstrap replicates using Model Finder Plus (MFP) to select a suitable model. From these trees, we manually identified paralogous sequences, analogous to the workflow used in other recent phylogenomic studies (Yang *et al*., 2015; Yang *et al*., 2018). Note that we use the term “paralog”, although the trees may not always allow distinguishing ancient duplication (true paralogs) from ancient allopolyploidisation events (homeologs). Detected paralogs were classified into three categories: (1) false positives, which were distantly related to the primary cluster of orthologous sequences and more likely to be erroneously clustered unrelated sequences; (2) outparalogs, representing copies that arose from a duplication in a lineage containing two or more taxa; and (3) inparalogs, which resulted from recent duplication events within the taxon in which duplication was observed. For gene trees containing multiple copies for particular taxa, we used a combination of branch support (higher; >80% bootstrap) and branch lengths (shorter) to manually identify the most likely ortholog candidates among the copies, and paralogs were excluded for phylogenetic analyses. The resulting datasets, both for nucleotide and amino acids sequences, are given in the Zenodo repository associated with this study (https://doi.org/10.5281/zenodo.17059203).

### Phylogenetic inferences

The final dataset of 708 paralog-filtered OGs was then analyzed using both coalescent and concatenation approaches. For the coalescent approach, ML gene trees were built using IQ-TREE with automatic selection of the best-fit substitution model (e.g., MFP option). Branch support was evaluated with 100 standard bootstrap replicates and 1000 ultrafast bootstrap replicates (Minh *et al*., 2013). The species tree was inferred using ASTRAL v5.7.3, estimating local posterior probabilities for branch support (Sayyari & Mirarab, 2016). For the concatenation analyses, the gene alignments were concatenated into a supermatrix using PhyKIT (Steenwyk *et al*., 2021), and tree inference performed in IQ-TREE, utilizing a range of protein and nucleotide substitution models selected based on model fit (Kalyaanamoorthy *et al*., 2017). Branch support in the concatenation analyses was estimated with 1000 ultrafast bootstrap replicates and 500 standard bootstrap replicates. Trees were visualized and annotated with the FigTree software (http://tree.bio.ed.ac.uk/software/figtree/) and the R package ggtree (Wickham, 2016).

### Gene duplication counting and WGD inference

To investigate gene duplications (GD) as potential indicators of whole-genome duplication (WGD) events, we employed a phylogenomic approach using protein-coding nucleotide sequences, building on the paralog filtering step outlined earlier. To ensure robust analyses, we excluded four taxa due to either low BUSCO completeness scores (< 35%) or low sequence representation (<60% of relevant OGs detected) (Table S2). Outparalogs, when identified, were considered evidence of gene duplications that occurred at an ancestral level, following established protocols in phylogenomics (Ren *et al*., 2018; Guo *et al*., 2020). A small number of OGs had topologies too complex to confidently identify outparalogs manually and were excluded from the analysis, leaving 675 BUSCO OGs.

Since Bryopsidales are an ancient group (ca. 500-600 Ma, Verbruggen *et al*., 2009; Hou *et al*., 2022), some other commonly used methods to estimate WGD events are unlikely to be successful with our data. Ks distributions saturate over long time scales and are unreliable when gene retention rates are < 10% (Tiley *et al*., 2018). Synteny analysis requires near-complete genome assemblies, which are unavailable for the majority of taxa. As a consequence, we rely mostly on GD counting methods. First, we carried out manual counting based on interpretations of gene trees. Second, we performed gene tree reconciliation using Notung, applying a 100-bootstrap support threshold (Vernot *et al*., 2008). The program GuestTreeGen from GenPhyloData was used to simulate gene trees (Sjöstrand *et al*., 2013), using birth and death rates calculated from Notung. For each birth and death rate, we simulated 20,000 trees, and empirical p-values of our observed duplication counts were found by sampling the 675 OG trees 10,000 times. Lastly, WGDgc was used on gene counts to test individual branches for WGD under a maximum-likelihood model (geomMean=1.1, conditioning=”oneOrMore”, with birth, death and retention rates found by maximum-likelihood estimation) (Rabier et al., 2013). The likelihood ratio test was used to test each WGD hypothesis, using previously published significance levels (Rabier et al., 2013).

Since *Caulerpa lentillifera* was previously found to have undergone WGD (Xu *et al*., 2025), that a high-quality genome assembly is available for this species, and that we considered the *Caulerpa* lineage recent enough to possibly retain evidence of synteny, we carried out synteny analysis between *C. lentillifera* chromosomes and our draft genome assemblies of *C. brownii*, *C. verticillata* and *C. webbiana.* Contigs over 10 kb in length from these genomes that aligned to *C. lentillifera* chromosomes with blastn were selected, and dot plots were created with wgdi (Sun *et al*., 2022), based on diamond blast hits of longest protein isoforms found with the wgdi script rundiamond.py.

### Functional annotation of duplicated genes

Functional annotations of the duplicated genes were performed using a combination of BlastKOALA, KofamKOALA, and BLASTp searches. BlastKOALA (Kanehisa *et al*., 2016) and KofamKOALA (Aramaki *et al*., 2020) web servers were used to annotate sequences against the KEGG Orthology (KO) database (Table S3). BLASTp searches were conducted using the BLAST+ suite (v2.13.0) against the manually curated UniProtKB/Swiss-Prot protein database (Boutet *et al*., 2016) with an E-value cutoff of 1e-05, retrieving the top 10 hits for each query sequence. Outputs from all three approaches were integrated, prioritizing KEGG-based annotations, while UniProtKB annotations were utilized to resolve ambiguous or missing functions for duplicated genes not assigned by other approaches. Finally, the functions of duplicated genes were visualized in R v4.4.0 using the ggplot2 package (Wickham, 2016).

## Results

### Highly resolved relationships of Bryopsidineae and Halimedineae

The 708 nuclear genes that we selected, with a minimum of 80% taxon coverage for each, resulted in very well-resolved phylogenies (Fig. 1A) using both the coalescent and concatenation approaches. We obtained largely consistent results using nucleotide or amino acid alignments, various substitution models, and different outgroup configurations. While we inferred phylogenies using multiple outgroup combinations, we present here the tree constructed with *Chlorophyceae* as the outgroup. Phylogenies based on alternative outgroup configurations are provided in the supplementary figures (Fig. S1-30) and all gene trees and concatenated trees are available in the Zenodo repository.

**Fig. 1.**
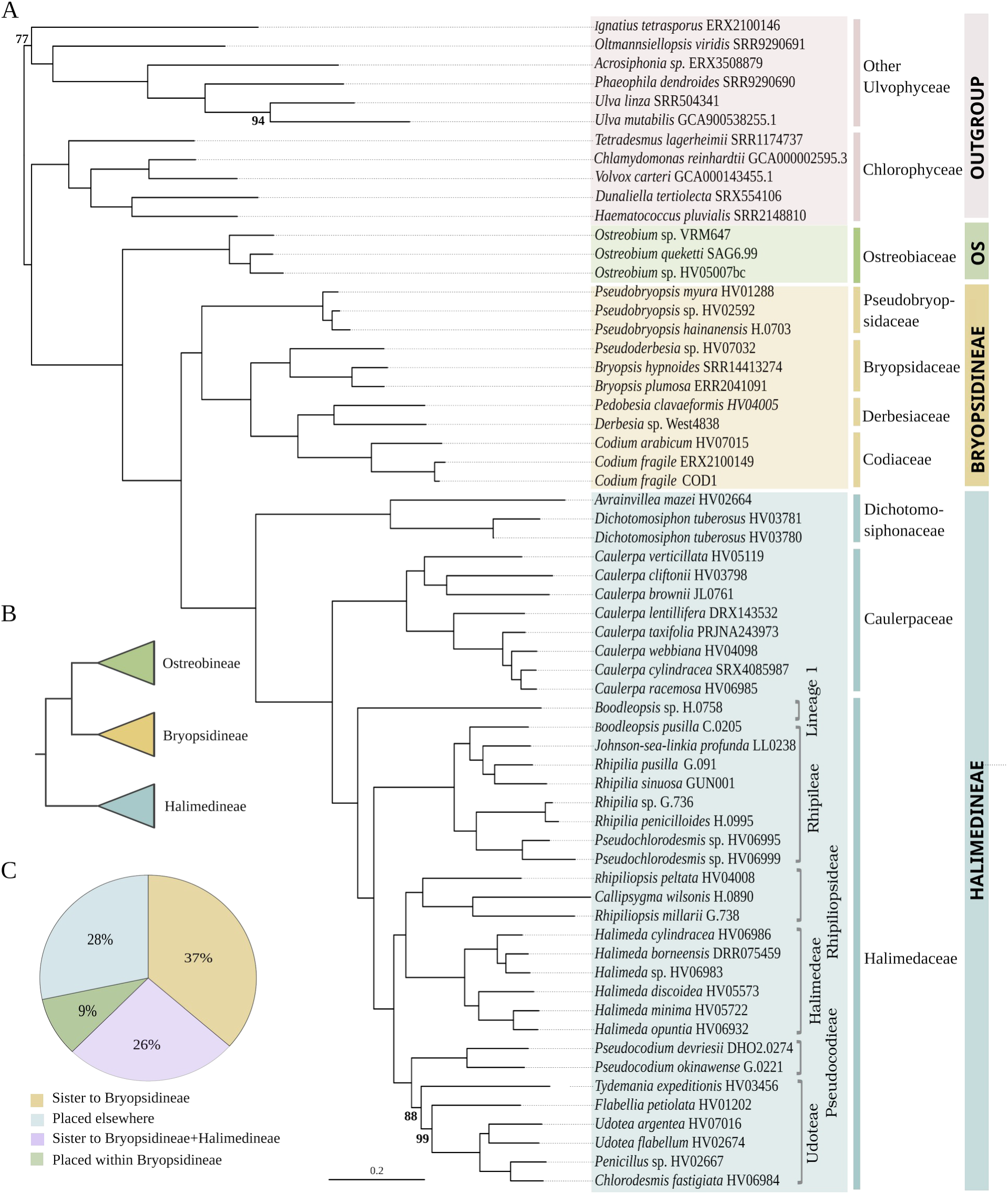
Phylogenetic relationships of Bryopsidales inferred from 708 nuclear genes. (A) Maximum likelihood phylogeny based on the concatenated nucleotide sequence alignment of protein-coding genes, using the GTR+F+I+R7 model. Branches without values received full support. (B) Simplified diagram of the topology inferred from a multispecies coalescent model, showing Ostreobineae as sister to Bryopsidineae. (C) Pie chart representing the percentage of gene trees supporting alternative Ostreobineae placements.

In the suborder Bryopsidineae, we recovered monophyletic families and fully resolved relationships within and among them (Fig. 1A). The previously difficult-to-place genus *Pseudobryopsis* was consistently placed as an early-branching lineage of this suborder, sister to all remaining families. Unlike previous studies that used a single member of *Pseudobryopsis*, our analysis included three species, which were robustly placed together. Among the remaining families, Bryopsidaceae was recovered as the sister group to a clade comprising Derbesiaceae and Codiaceae, with perfect support in all analyses.

In the suborder Halimedineae, the previously uncertain relationships between the tribe-level lineages (Udoteae, Pseudocodieae, Halimedeae, Rhipiliopsideae, Rhipileae and the unnamed “Lineage 1”, following the numbering in Cremen *et al.,* 2019) were recovered with very high support. Only the position of Rhipileae, recovered as sister to the clade composed of Rhipiliopsideae, Halimedeae, Pseudocodieae and Udoteae, exhibited slightly lower bootstrap support (90-98%) in some amino acid-based phylogenies. The *Caulerpa* topology was in line with its accepted infra-generic classification (Draisma *et al.,* 2014), yet offered substantially stronger support than previous analyses. It is worth noting that *Caulerpa verticillata* (*Charoideae* subgenus), a lineage with highly labile placement in previous trees, was confidently recovered as sister to *C. cliftonii* and *C. brownii* in DNA analyses even though three of our amino acid-based phylogenies showed uncertainty about its placement (Fig. S23, S28-29). The genus *Boodleopsis* was seen to occur in two lineages, with an unidentified *Boodleopsis* species (sample H.0758, culture strain West4288) placed in Lineage 1 and *Boodleopsis pusilla* in the Rhipileae tribe. In the Udoteae, *Tydemania expeditionis* was recovered as sister to the rest of the tribe with 69-100% bootstrap support, and *Udotea argentea* HV07016 and *U. flabellum* recovered as sister taxa (94-100% support) in most trees. However, the placement of *T. expeditionis* and *U. flabellum* is uncertain in some trees (Fig. S21-29).

### Phylogenetic position of Ostreobineae

All analyses fully supported relationships among the three *Ostreobium* strains (Fig. 1 and Fig. S1-23), and the suborder Ostreobineae as a whole was recovered as sister to all remaining Bryopsidales (suborders Bryopsidineae and Halimedineae) with maximum support (100%) in almost all analyses. The only exception is the ASTRAL coalescent tree using Dasycladales and Cladophorales as outgroups, which recovered Ostreobineae as sister to Bryopsidineae with full local posterior probability (Fig. 1B; Fig. S1,S6). Our analysis of the position of Ostreobineae across the 708 gene trees showed that 37% of gene trees positioned Ostreobineae as sister to Bryopsidineae, 26% to the clade consisting of Bryopsidineae+Halimedineae, 9% within Bryopsidineae, and 28% elsewhere in the tree (Fig. 1C). The average support values for the relationship of Ostreobineae as sister to Bryopsidineae were comparatively lower than for it being sister to the clade consisting of Bryopsidineae and Halimedineae (Fig. S31).A vast majority of the analyses, including all those with other outgroup combinations, placed Ostreobineae as sister to Bryopsidineae+Halimedineae.

### Gene and genome duplication events in major clades of siphonous green algae

With our dataset of 675 nuclear genes, and following taxon filtering based on BUSCO scores (Fig. S32) we applied various methods to quantity gene duplications (GD) and identify branches with putative WGD events. Our manual gene tree reconciliation detection of outparalogs identified 14 branches with GDs (Fig. 2A), with two having particularly large numbers of GDs: OST (comprising the *Ostreobium* lineage) with 8.44% of genes showing duplications and CAUL2 (including *Caulerpa lentillifera*, *C. taxifolia*, *C. webbiana*, *C. racemosa*, and *C. cylindracea*) with 7.56% of genes being duplicated. An example of the DBP5 gene, an RNA helicase involved in mRNA export from the nucleus, shows clear evidence for duplication in the CAUL2 and OST branches (Fig. 2B). Automated gene tree reconciliation inferred more GDs overall across the tree, with OST and CAUL2 nodes having 15.7% and 13.4% of genes duplicated, respectively (Fig. 2C). The Caulerpaceae exhibited frequent GDs, with four nodes within the family showing evidence of GD, including the CAUL2 clade mentioned above and the branch subtending *C. taxifolia*, *C. webbiana*, *C. racemosa*, and *C. cylindracea* (CAUL3; 1.78% with manual detection and 7.1% with Notung; Figs 2A,C). Other lineages had sporadic manually counted GDs at the family or tribe level, typically with one or two genes except for Bryopsidaceae (BRY; 4 GDs) and the Halimedeae tribe (HALTR; 3 GDs), which also had elevated GD counts inferred by Notung (Fig. 2C). The NO87 node consisting of *Halimeda cylindraceae, H. borneensis and Halimeda* sp. HV06983 also had higher GD counts found by Notung, but not in our manual counts (Fig. 2C).

**Fig. 2.**
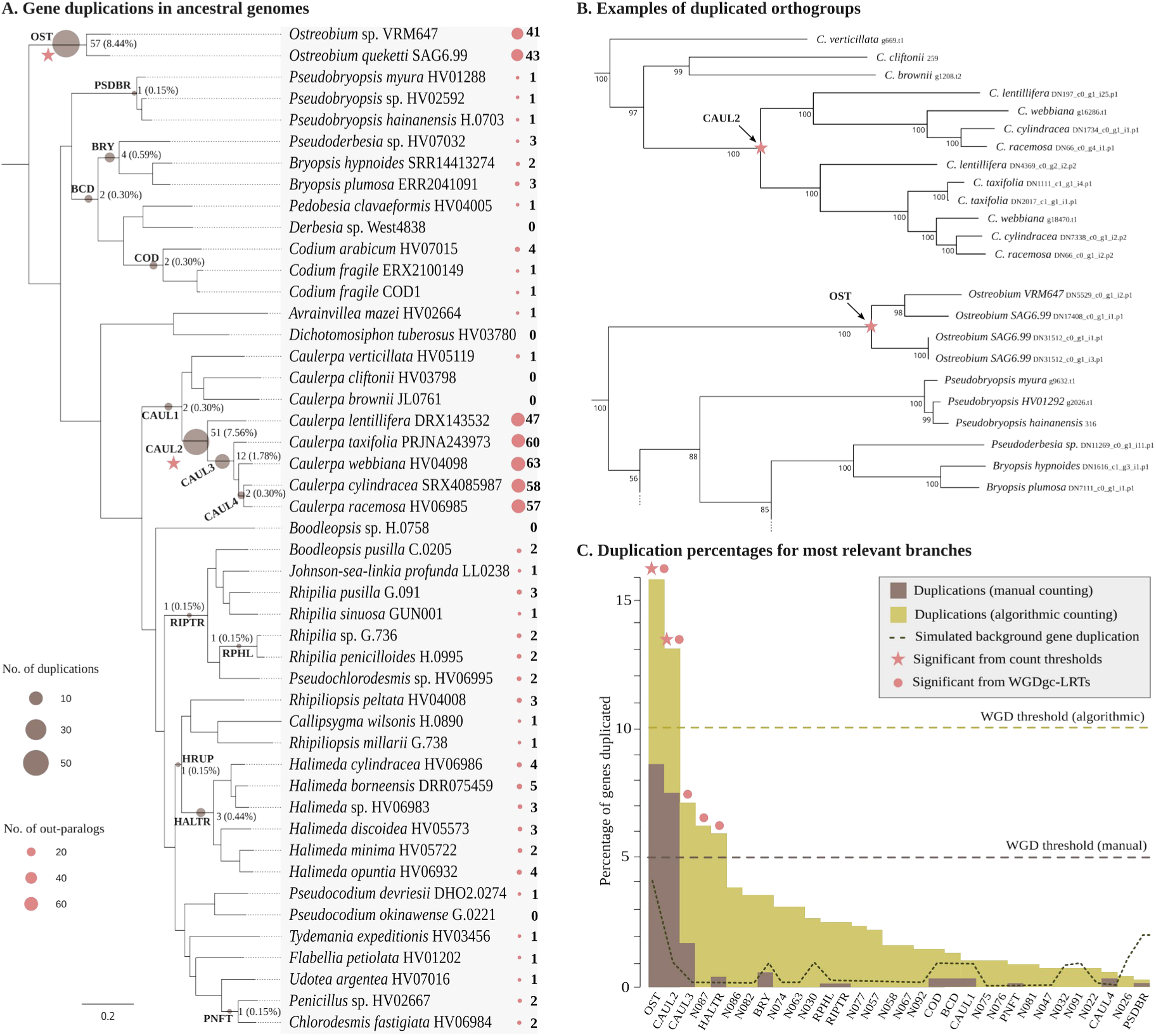
Gene and genome duplications in Bryopsidales (A) Outparalog based duplications mapped onto the phylogeny, with circles indicating the ancestral branches where duplications were inferred, alongside the percentage of gene duplications observed at each branch. Colored circles and numbers following taxon names represent the number of outparalogs identified within each genome. Stars denote proposed WGD branches. (B) Two fragments of the DBP5 gene (DDX19, KEGG K18655, BUSCO 1477at3041) outlining our manual detection of outparalogs, focused on gene duplications occurring in the CAUL2 and OST lineages. (C) Comparative summary of gene duplication rates from manual outparalog counting, gene tree reconciliation with Notung, gene-tree simulations, and ML statistical tests for WGD significance across key branches.

Two nodes, OST and CAUL2, exceeded all thresholds for WGD suggested in the literature (Fig. 2C), being 5% when using manual counting methods (Guo et al., 2020) and 10% for Notung (Cai et al., 2019). Gene tree simulations further confirmed that only OST, CAUL2, and CAUL3 had significantly more GDs than expected under a null model without WGD (Fig. 2C). Duplication counts from algorithmic methods were higher than those counted manually, and for several branches exceeded the simulated values under the null model, yet most of these still fell below the 5% threshold we applied to indicate WGD. The maximum likelihood-based WGDgc approach, based only on gene counts across species to test for the presence of WGDs, again identified OST and CAUL2, as well as CAUL3, HALTR, and N087 as significant (Fig. 2C). The detailed results for simulations and WGDgc are given in the Zenodo repository.

Analysis of synteny between the high-quality *C. lentillifera* genome and our draft assemblies of three other *Caulerpa* species revealed that *C. webbiana* contains similar duplicated chromosomes (Figs 3A, S34). Conversely, the earlier-branching *C. verticillata* and *C. brownii* contain only single chromosomes, and the best blast hits of their protein-coding sequences are equally dispersed across both chromosomes of a duplicated pair in *C. lentillifera*. This confirms that a WGD event occurred along the CAUL2 branch subtending the MRCA of *C. webbiana* and *C. lentillifera*. Synteny analysis was attempted for the *Ostreobium* lineage, which also has a published genome (Iha *et al*., 2021), but was inconclusive, likely due to a combination of ancient divergence times (Verbruggen *et al*., 2009a), low gene densities and fragmented assemblies (Iha *et al*., 2021).

**Fig. 3.**
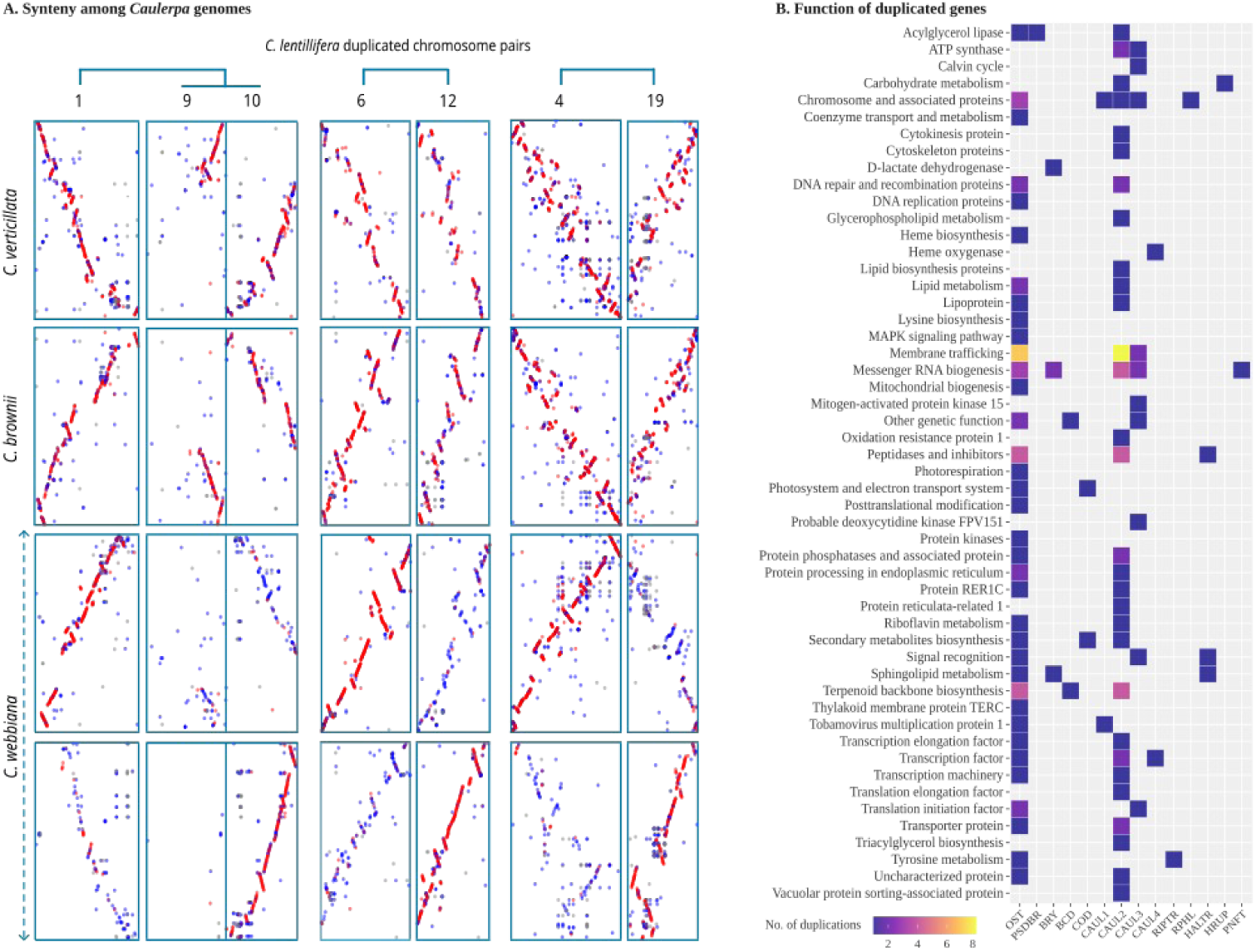
Synteny and functions of duplicated genes. (A) Synteny analysis highlighting duplicated chromosomal blocks from available genomes of the core Caulerpaceae lineage. Red dots indicate best blast hits of proteins, and blue are other significant hits. Note that entire chromosomes from *C. lentillifera* are shown, while for other species we used concatenations of smaller contigs. (B) Functional annotations of duplicated genes observed at each ancestral node.

Functional annotation of outparalog-detected duplications revealed that duplicated genes in the OST and CAUL2 lineages belonged to several functional categories (Fig. 3B, Fig. S33). Both nodes had duplicated genes with functions involved in membrane trafficking, terpenoid backbone biosynthesis, peptidases and inhibitors, DNA repair and recombination, and chromosome-associated proteins. However, the OST duplications had functions related to protein kinases, photosystems, transporters, signaling, DNA replication, mitochondrial biogenesis, lysine biosynthesis, and posttranslational modification. In contrast, CAUL2 duplications were rich in protein phosphatases, vacuolar protein sorting, cytoskeletal proteins, and genes involved in lipid biosynthesis. A full list of the node names is given in Fig. S34.

## Discussion

### Nuclear genome data resolves the Bryopsidales tree of life

Based on extensive sampling across the genus diversity of Bryopsidales and whole genome and/or transcriptome shotgun sequencing, we present the most highly resolved Bryopsidales phylogeny to date, and the first to sample extensively from nuclear genomes. Our analysis included a BUSCO-based dataset of 708 nuclear genes and achieved 93-96% of the nodes with maximal support across both coalescent and concatenation approaches (Fig. S1-29), emphasizing the robustness of our phylogenetic results, which remained largely consistent across various models of sequence evolution, sequence types, and outgroup selections.

Our data consistently inferred *Pseudobryopsis* as an early-branching lineage of the Bryopsidineae, resolving inconsistencies seen in prior work, where it was placed with varying support (58-96%) in different places, and relationships were based on a single species and solely plastome data (Cremen *et al*., 2019). Our results oppose the classical interpretation of *Pseudobryopsis*’ relationship with *Bryopsis* (i.e., assigned to the Bryopsidaceae) and *Derbesia* (i.e., assigned to the Derbesiaceae) based on their shared uniaxial morphology and sporic meiosis, a process involving meiotic division during spore formation that results in a diplohaplontic life cycle (Hillis-Colinvaux, 1984; Vroom *et al*., 1998; Woolcott *et al*., 2000). Instead, our recovery of *Pseudobryopsis* as a distinct, early-branching lineage within the Bryopsidineae suggests that these shared traits may instead represent ancestral characteristics of the entire Bryopsidineae suborder, which were subsequently lost in favor of a diplontic life cycle in *Codium*. The inclusion of three taxa from *Pseudobryopsis* as opposed to a single species in previous analysis, allowed us to conclusively place it, helping to break up the long branch that it presented in previous analyses (Cremen *et al*., 2018; Marcelino & Verbruggen, 2016; Verbruggen *et al*., 2017). Moreover, our use of a large-scale nuclear gene dataset fully resolved the relationships of families and genera within Bryopsidineae, where previous studies had recovered several branches with low to moderate support (Lam & Zechman, 2006; Curtis *et al*., 2008; Verbruggen *et al*., 2009a; Leliaert *et al*., 2014; Cremen *et al*., 2019).

Within the Halimedineae, our findings provide strong nuclear genome evidence for relationships among families and tribes. These involve areas of substantial uncertainty observed in marker gene trees, several of which remained unresolved even in analyses of whole plastomes (Lam & Zechman, 2006; Verbruggen *et al*., 2009a; Cremen *et al*., 2019; Huisman & Verbruggen, 2023). Additionally, our nuclear genome data strongly supports previous hypotheses about polyphyletic genera within this suborder. For instance, *Boodleopsis* was recovered as polyphyletic, appearing as sister to all other Halimedaceae (Lineage 1) and as a separate lineage within the Rhipileae tribe. A similar scenario played out for *Pseudochlorodesmis*. While the samples we had access to for this work were all in the Rhipilieae lineage, other papers clearly show that this genus is polyphyletic, with representatives appearing in the Rhipilieae, Caulerpaceae and Cremen’s Lineage 2 (Verbruggen *et al*., 2009c; Draisma *et al*., 2014; Marcelino & Verbruggen, 2016; Cremen *et al*., 2019).

*Boodleopsis* and *Pseudochlorodesmis* are structurally similar, with uniaxial siphons that have both creeping and upright axes. The creeping axes spread out forming mats in loose muddy substrates in *Boodleopsis* or growing endolithically inside of reef limestones in *Pseudochlorodesmis*, with the simple, dichotomously branched uprights sticking out of the mud or rock. Morphology-based phylogenies have suggested a sister relationship between *Pseudochlorodesmis* and *Boodleopsis* (Vroom *et al*., 1998) indicating their morphological similarity. The molecular data presented here, along with some previous phylogenies (Cremen *et al.,* 2019; Huisman & Verbruggen, 2023), clearly do not support a sister relationship between these genera. Instead, they suggest that the simple branched uniaxial morphologies consisting of spreading and upright siphons seen in these genera represent ancestral traits of the core Halimedineae and possibly all Bryopsidales, a hypothesis that requires further testing through denser taxon sampling and ancestral state estimation.

Our analyses resolved the placement of *Caulerpa verticillata* in nearly all phylogenies, addressing the discrepancies reported in earlier studies based on marker genes or whole plastomes (Kazi *et al*., 2013; Draisma *et al*., 2014; Cremen *et al*., 2019). This suggests that the partial resolution in previous work likely stemmed from the smaller datasets employed. The differences in the placement of *Tydemania expeditionis* and *Udotea* species appear to be influenced by the type of input sequences used. While our amino acid-based phylogenies align with previous studies that utilized amino acid data (Verbruggen *et al*., 2009a; Cremen *et al*., 2019), the relationships observed in nucleotide-based phylogenies in our analysis provided stronger support (Fig. 1). Despite the overall resolution, some taxa (e.g., *C. verticillata*, *T. expeditionis*, and two *Udotea* species) remained unresolved in a few amino acid-based phylogenies, particularly with specific outgroup combinations. Given that amino acid sequences are one third the length and less variable than nucleotides and may reduce resolution among closely related taxa (Simmons, 2000), we put forward our nucleotide trees as a well-resolved hypothesis to enhance the accuracy and resolution of future phylogenetic studies of this order, while emphasizing the need to examine model adequacy and incorporate dense taxon sampling.

Our work shows that an approach consisting of a combination of transcriptomics and shotgun genomics can lead to well-resolved genome-scale phylogenies within a major group of Chlorophyta. In contrast to land plants and particularly angiosperms, where biodiversity genomics has already led to extensively sampled genome-scale phylogenies (e.g. Grass phylogeny working group, 2025; Colli-Silva *et al*., 2025), similar work had not been done previously for any group of the Chlorophyta. No target capture probes have been developed to work universally across the Chlorophyta, and due to them being more than a billion years old and hence very diverse molecularly, it seems unrealistic that such probes can be developed. Our approach was based on transcriptomics for species where we could obtain fresh material, including some very small filamentous species like *Pseudochlorodesmis*, and draft genomes assembled from short-read shotgun DNA data of silica-gel preserved samples for species that were difficult to source fresh. This approach is economical in the absence of DNA capture approaches and the data we have obtained here may facilitate the development of group-specific capture probes like those developed increasingly frequently for angiosperm groups (e.g. Mandel *et al*., 2017).

### Genome duplication and evolutionary adaptations in key Bryopsidales lineages

Whole genome duplications (WGDs) have long been recognized as a powerful evolutionary mechanism that may facilitate innovation and adaptation in a range of organisms, and land plants in particular. In the Chlorophyta lineage, however, very little evidence currently exists on the topic. We inferred WGD events using both manual and algorithmic gene tree reconciliation approaches. We mainly base our interpretations on the outparalog-based manual counts The algorithmic reconciliation methods show similar trends, but are prone to higher rates of false positives arising from gene tree errors, uneven taxon representation, and sensitivity to parameter settings (Hahn, 2007; Szöllősi et al., 2015). This is also observed in our analysis that reconciliation results across the nodes are generally higher than outparalog detection based GDs counting. Across all techniques that we applied, WGDs were inferred to have occurred at least twice throughout Bryopsidales evolution, in the OST and CAUL2 lineages, using calibrated thresholds for the respective methods (5% for manual and 10% for Nutong). The fact that our dataset is based on BUSCO genes adds to our confidence in this result, as these conservative genes are nearly universally single-copy in Chlorophyta, and therefore these genes may be more duplication-resistant than other gene families (Li *et al*., 2016) and, consequently, GD rates from this dataset are likely to represent a lower boundary for genome-wide GD rates. A recent study also reported evidence of duplication in *Caulerpa lentillifera*, proposing that this duplication occurred after its divergence from the *Bryopsis* lineage (Xu *et al*., 2025). Our densely sampled datasets and analyses substantially refine that result, showing that the duplication in question was relatively recent, taking place in the common ancestor of the core *Caulerpa* lineage (CAUL2).

Molecular clocks and fossils suggest that the Ostreobineae adopted an endolithic lifestyle, living inside of limestone rock, in the early Paleozoic (>500 million years ago (Vogel, 1993; Vogel & Brett, 2009; Verbruggen *et al*., 2009a; Marcelino & Verbruggen, 2016). Our findings indicate that a genome duplication may be connected with that transition and that some key genomic features from that duplication are still preserved in modern *Ostreobium* species. For instance, the expansion of photosystem-related proteins, as inferred for the ancestor OST (Fig. 3B), suggests enhanced light-harvesting capabilities for photosynthesis under low-light conditions. Furthermore, the proliferation of transporter proteins (i.e. Ca^2+^ transporter) likely supported algal growth and development in limestone substrates, aiding in processes such as decalcification (Garcia-Pichel *et al*., 2010). The expansion of coenzyme transporters likely supported algal growth within coral skeletons, particularly for coenzymes like vitamin B12, which algae acquire from surrounding bacteria in the endolithic environment (Croft *et al*., 2005). Despite adaptation to dim light within coral skeletons, some *Ostreobium* strains live in full-spectrum sunlight, such as in intertidal oyster shells (Kornmann & Sahling, 1980), where it may experience high-intensity light during low tide. High-light exposure generates excessive reactive oxygen species (ROS), leading to oxidative stress (Pospíšil, 2016; Foyer, 2018), which can cause DNA damage and, if unmitigated, interfere with growth or even cause death (Dizdaroglu *et al*., 2002; Tuteja *et al*., 2009). Given the prevalence of these conditions in *Ostreobium*’s habitat, it is interesting to think about how this alga maintains ROS homeostasis and preserves DNA integrity. The expansion of DNA repair and chromosome-associated proteins in *Ostreobium*’s ancestors may have evolved as an adaptive response to these challenges. Moreover, duplications in terpenoid biosynthesis pathways, known for their role in chemical defenses against grazers (Paul & Fenical, 1983; Hay, 1996), suggest that these organisms may have inherited defensive strategies from their ancestors to thrive in herbivory-prone tropical reef environments. Our work shows that some of the expansions (e.g., photosystem-related, transporter proteins, and ROS homeostasis) interpreted as adaptations to an endolithic lifestyle in the genome of a modern *Ostreobium* strain (Iha *et al*., 2021), may result from an ancestral whole-genome duplication event.

The Caulerpaceae family, which originated >215 million years ago (Verbruggen *et al*., 2009a; Draisma *et al*., 2014) and is particularly known for its invasive species and larger morphologies within Bryopsidales, appears to be a hotspot for gene and genome duplications. The CAUL2 and CAUL3 lineages align with the species-rich “core *Caulerpa*” sublineage (*Caulerpa* subgenus *Caulerpa*) in contemporary classifications (Kazi *et al*., 2013; Draisma *et al*., 2014), which is inferred to be ca. 209 (stem node) to 86 (crown node) million years old (Draisma *et al*., 2014). Species in this subgenus are often characterized by larger thallus morphologies and widespread distribution (Belton *et al*., 2019; Terada *et al*., 2012) and several among them are highly invasive (Meinesz *et al*., 1995; Verlaque *et al*., 2003; Walters *et al*., 2006; Amat *et al*., 2008; Piazzi *et al*., 2016). Based on our findings, we hypothesize that the duplication involved in terpenoid biosynthesis pathways, which contribute to anti-grazing defenses, may have enhanced the resilience of this lineage, enabling them to thrive across diverse environments. Similarly, duplication of genes involved in triacylglycerol biosynthesis may reflect increased tolerance to abiotic stress (Lu *et al*., 2020). Moreover, ancestral gene duplications, as first proposed by Arimoto et al (2019) in the *Caulerpa lentillifera* genome, may have played a role in shaping the distinct morphologies of core *Caulerpa* within the Bryopsidales. For instance, in *Caulerpa* growth occurs at the frond apex, which contains a unique transcriptomic profile. Highly-expressed apex transcripts include coat proteins and kinases, protein functions that subsequently were found to be expanded in a genome-wide analysis of *C. lentilifera* (Ranjan *et al*. 2015, Arimoto *et al*., 2019). Our findings also indicate ancestral duplication of genes with apex-related functions, such as those related to membrane trafficking, protein sorting, protein dephosphorylation, the cytoskeleton and the endoplasmic reticulum (Ranjan *et al*., 2015).

Our analysis identified functional categories highly duplicated in both the *Caulerpa* and *Ostreobium* WGD events, such as peptidases, and terpenoid backbone biosynthesis, which play roles in development and stress responses (Schaller, 2004; Kohli *et al*., 2012; Roberts *et al*., 2012; El Amerany, 2024), and membrane trafficking proteins, crucial for growth and stress recovery, suggesting adaptive advantages of the duplication of these genes in several parts of the Bryopsidales tree of life.

It is important to note that our analysis of duplication events focuses on out-paralog identification, and therefore only captures ancestral WGD events. Recently, *Halimeda opuntia* was shown to have experienced at least three genome duplication events in the form of consecutive, intraspecific polyploidisation events. These events were estimated to have occurred recently in the *H. opuntia* lineage, and would therefore not be identified as an ancestral WGD with our approach (Zhang *et al*., 2024). In contrast, our analyses detected signals of probable WGD at two nodes within Halimedaceae (HALTR and N087), but only in likelihood-based WGDgc testing on gene counts only. Since these nodes did not surpass the WGD thresholds applied here and no genome assemblies are currently available for synteny-based validation, we interpret them cautiously as tentative WGD signals. Nonetheless, they represent promising candidates for future investigation, with genomic resources expected to be critical for confirming or rejecting their status as WGDs.

While our study was based on 675 BUSCO OGs, representing only part of the genome, it opens the door for future whole-genome phylogenomic studies to more comprehensively explore the genome duplications in Bryopsidales and their potential adaptive nature. Additionally, differential gene expression analyses might help to further test these hypotheses, particularly to assess whether duplicated genes are functionally retained and actively expressed in extant descendant genomes. Although WGD is often portrayed as a key driver of evolutionary innovation, it is not always linked to large-scale evolutionary success, some events may have had minimal or no lasting impact (Mayrose et al. 2011, 2015). In fact, WGD has been described as a high-risk process with the potential for long-term payoffs (Yant & Bomblies, 2015; Clo & Kolář, 2021), reigniting debate over its role in land plant evolution and diversification (Soltis *et al*., 2014; Kellogg, 2014; Clark & Donoghue, 2017).

Our work expands the much more extensive knowledge on WGD in angiosperms to the Chlorophyta lineage, providing evidence for this process and its association with functional changes. Whole genome duplication has been understudied in green algae, with only a very small number of lineages being identified as potential candidates in the 1KP analyses (One Thousand Plant Transcriptomes Initiative, 2019). At face value, this would suggest WGD are rare in Chlorophyta, but we want to be cautious with such statements due to the very different time-frames. The angiosperms, where WGD are well characterised, are much younger than the Chlorophyta, with the whole angiosperm lineage being comparable in age to the families (or even genera) of Chlorophyta (e.g., Verbruggen *et al*., 2009a). With the exception of the *Caulerpa* duplication (between 209-86 Ma), we are looking at substantially older duplications, and we do not doubt that additional events can be found in younger groups if taxon sampling continues to increase. The duplication signal we find on node CAUL3 already suggests that it may be possible that *Caulerpa* has experienced multiple rounds of WGD, and the polyploid *Halimeda* genome along with the duplication signal for two ancestral *Halimeda* lineages (HALTR and N087) further indicates that it occurs at more recent timescales (Zhang *et al*., 2024). Finally, the conservative nature of our BUSCO-based analyses may underestimate true WGD rates.

In plants, ancient WGDs have been repeatedly linked to ecological and functional innovation, often enabling adaptation to new or extreme environments. For example, seagrasses such as *Zostera marina* and *Posidonia oceanica* show ancient WGDs associated with marine adaptation, including improved osmoregulation, light utilization, and stress tolerance (Ma *et al*., 2024). Similarly, the mangrove *Avicennia marina* experienced two recent WGDs that expanded salt-tolerance transcription factors, aiding its success in intertidal habitats (Feng *et al*., 2023; Xu et al., 2023). In arid regions, lineage-specific WGDs in *Kalanchoe* have been linked to enhanced drought tolerance (Yang *et al*., 2017). Beyond habitat-specific adaptations, polyploidy is also thought to contribute to the ecological breadth and invasiveness of many angiosperms, as genomic redundancy can provide stress buffering, trait diversification, and phenotypic plasticity, thereby promoting colonization of challenging or novel environments (Treier *et al*., 2009; Pandit *et al*., 2011; te Beest *et al*., 2012; Wan *et al*., 2020). WGDs in higher plants have further been associated with increased cell and organ size, greater morphological complexity, enhanced biomass, and the diversification of secondary metabolites such as glucosinolates, which together improve competitive ability and grazing defense (Otto, 2007; Edger & Pires, 2009; Edger *et al*., 2015). Drawing on these parallels, our results suggest that WGDs in core *Caulerpa* lineages may also contribute to its ecological versatility, grazing resistance, and success in diverse marine environments.

## Acknowledgements

This work was supported by the Australian Biological Resources Study (Activity 4-G046WSD to HV), the Australian Research Council (DP200101613 to HV), and the Fundação para a Ciência e a Tecnologia (CEECIND:2023.06155 to HV). RH received support from the Hansjörg Eichler Scientific Research Grant through the Australian Systematic Botany Society and is funded by the Prime Minister’s Fellowship of the People’s Republic of Bangladesh. We utilized computational resources from the Spartan Supercomputer platform at the University of Melbourne, and we also extend our thanks to the Galaxy Australia platform for access to their high-performance computing network during data processing. Special thanks go to Amelia Hastings and Myles Courtney for their assistance with sampling, and to John A. West for his valuable suggestions on maintaining cultures and providing a few of the strains used here. We are grateful to Md. Abdullah Al Kamran Khan for his help with DNA extraction, to the members of the Verbruggen lab for insightful discussions, and to three anonymous reviewers for their constructive comments.

## Author contributions

RH and HV developed the conceptual framework. RH carried out the research, performed data analyses. RH drafted the manuscript, with contributions from SB and HV. All authors contributed to data analysis and visualization. HV obtained funding, carried out field work and provided supervision. All authors made critical revisions to the manuscript and approved the final version.

## Data availability

The raw sequencing reads are deposited in the NCBI SRA under BioProject accession number PRJNA1216519. Phylogenetic trees and sequence alignments for the 708 orthogroups are available on Zenodo (DOI: https://doi.org/10.5281/zenodo.17059203).

## References

Amat JN, Cardigos F, Santos RS. 2008. The recent northern introduction of the seaweed *Caulerpa webbiana* (Caulerpales, Chlorophyta) in Faial, Azores Islands (north-eastern Atlantic). Aquatic Invasions 3: 417–422.

Aramak T, Blanc-Mathieu R, Endo H, Ohkubo K, Kanehisa M, Goto S, Ogata H. 2020. KofamKOALA: KEGG Ortholog assignment based on profile HMM and adaptive score threshold. Bioinformatics 36: 2251–2252.

Arimoto A, Nishitsuji K, Higa Y, Arakaki N, Hisata, K, Shinzato C, Satoh N, Shoguchi E. 2019. A siphonous macroalgal genome suggests convergent functions of homeobox genes in algae and land plants. DNA Research 26: 183–192.

Bankevich A, Nurk S, Antipov D, Gurevich AA, Dvorkin M, et al. 2012. SPAdes: a new genome assembly algorithm and its applications to single-cell sequencing. Journal of Computational Biology 19: 455–77.

Belton GS, Draisma SG, Prud’homme van Reine WF, Huisman JM, Gurgel CFD. 2019. A taxonomic reassessment of *Caulerpa* (Chlorophyta, Caulerpaceae) in southern Australia, based on tuf A and rbcL sequence data. Phycologia 58: 234–253.

Benson G. 1999. Tandem repeats finder: a program to analyze DNA sequences. Nucleic acids research 27: 573–580.

Boutet E, Lieberherr D, Tognolli M, Schneider M, et al. 2016. UniProtKB/Swiss-Prot, the manually annotated section of the UniProt KnowledgeBase: how to use the entry view. Methods in Molecular Biology 1374: 23–54.

Bowers JE, Paterson AH. 2021. Chromosome number is key to longevity of polyploid lineages. New Phytologist 231: 19–28.

Brůna T, Hoff KJ, Lomsadze A, Stanke M, Borodovsky M. 2021.BRAKER2: automatic eukaryotic genome annotation with GeneMark-EP+ and AUGUSTUS supported by a protein database. NAR genomics and bioinformatics 3: lqaa108.

Bulleri F, Airoldi L, Branca G, Abbiati M. 2006. Positive effects of the introduced green alga, Codium fragile ssp. tomentosoides, on recruitment and survival of mussels. Marine Biology 148: 1213–1220.

Bulleri F, Balata D, Bertocci I, Tamburello L, Benedetti-Cecchi L. 2010. The seaweed *Caulerpa racemosa* on Mediterranean rocky reefs: from passenger to driver of ecological change. Ecology 91: 2205–2212.

Cai L, Xi Z, Amorim AM, Sugumaran M, Rest JS, Liu L, Davis CC. 2019. Widespread ancient whole-genome duplications in Malpighiales coincide with Eocene global climatic upheaval. New Phytologist 221: 565–576.

Carretero-Paulet L, Van de Peer Y. 2020. The evolutionary conundrum of whole-genome duplication. American journal of botany 107: 1101.

Castro-Sanguino C, Bozec Y-M, Mumby PJ. 2020. Dynamics of carbonate sediment production by *Halimeda*: implications for reef carbonate budgets. Marine Ecology Progress Series 639: 91–106.

Charron G, Marsit S, Hénault M, Martin H, Landry CR. 2019. Spontaneous whole-genome duplication restores fertility in interspecific hybrids. Nature Communications 10: 4126.

Challis R, Richards E, Rajan J, Cochrane G, Blaxter M. 2020. BlobToolKit–interactive quality assessment of genome assemblies. G3: Genes, Genomes, Genetics 10: 1361–1374.

Cho CH, Park SI, Huang T-Y, Lee Y, Ciniglia C, Yadavalli HC, et al. 2023. Genome-wide signatures of adaptation to extreme environments in red algae. Nature communications 14: 10.

Clark JW, Donoghue PC. 2017. Constraining the timing of whole genome duplication in plant evolutionary history. Proceedings of the Royal Society B: Biological Sciences 284: p.20170912.

Clo J, Kolář F. 2021. Short-and long-term consequences of genome doubling: a meta-analysis. American Journal of Botany 108: 2315–2322.

Colli-Silva M, Pérez-Escobar OA, Ferreira CD, Costa MT, et al. 2025. Taxonomy in the light of incongruence: An updated classification of Malvales and Malvaceae based on phylogenomic data. Taxon 74: 361–385.

Cremen MCM, Leliaert F, Marcelino VR, Verbruggen H. 2018. Large diversity of nonstandard genes and dynamic evolution of chloroplast genomes in siphonous green algae (Bryopsidales, Chlorophyta). Genome Biology and Evolution 10: 1048–1061.

Cremen MCM, Leliaert F, West J, Lam DW, Shimada S, Lopez-Bautista JM, Verbruggen H. 2019. Reassessment of the classification of Bryopsidales (Chlorophyta) based on chloroplast phylogenomic analyses. Molecular Phylogenetics and Evolution 130: 397–405.

Croft MT, Lawrence AD, Raux-Deery E, Warren MJ, Smith AG. 2005. Algae acquire vitamin B12 through a symbiotic relationship with bacteria. Nature 438: 90–93.

Curtis NE, Dawes CJ, Pierce SK. 2008. Phylogenetic analysis of the large subunit rubisco gene supports the exclusion of *Avrainvillea* and *Cladocephalus* from the Udoteaceae (Bryopsidales, Chlorophyta) 1. Journal of Phycology 44: 761–767.

Cuypers TD, Hogeweg P. 2014. A synergism between adaptive effects and evolvability drives whole genome duplication to fixation. PLoS computational biology 10: e1003547.

Del Cortona A, Jackson CJ, Bucchini F, Van Bel M, et al. 2020. Neoproterozoic origin and multiple transitions to macroscopic growth in green seaweeds. *Proceedings of the National Academy of Sciences*, USA 117: 2551–2559.

Dizdaroglu M, Jaruga P, Birincioglu M, Rodriguez H. 2002. Free radical-induced damage to DNA: mechanisms and measurement. Free Radical Biology and Medicine 32: 1102–15.

Draisma SG, van Reine WFPh, Sauvage T, Belton GS, Gurgel CFD, Lim PE, Phang SM. 2014. A re-assessment of the infra-generic classification of the genus *Caulerpa* (Caulerpaceae, Chlorophyta) inferred from a time-calibrated molecular phylogeny. Journal of Phycology 50: 1020–1034.

Edger PP, Heidel-Fischer HM, Bekaert M, Rota J, et al. 2015. The butterfly plant arms-race escalated by gene and genome duplications. *Proceedings of the National Academy of Sciences*, USA 112: 8362–8366.

Edger PP, Pires JC. 2009. Gene and genome duplications: the impact of dosage-sensitivity on the fate of nuclear genes. Chromosome Research 17: 699–717.

El Amerany F. 2024. The role of terpenoids in plant development and stress tolerance. In Molecular and physiological insights into plant stress tolerance and applications in agriculture-Part 2. Bentham Science Publishers, pp. 71–98.

Flynn JM, Hubley R, Goubert C, Rosen J, Clark AG, Feschotte C, Smit AF. 2020. RepeatModeler2 for automated genomic discovery of transposable element families. Proceedings of the National Academy of Sciences, USA 117: 9451–9457.

Foyer CH. 2018. Reactive oxygen species, oxidative signaling and the regulation of photosynthesis. Environmental and Experimental Botany 154: 134–142.

Gabriel L, Bruna T, Hoff KJ, Ebel M, Lomsadze A, Borodovsky M, Stanke M. 2024. BRAKER3: Fully automated genome annotation using RNA-seq and protein evidence with GeneMark-ETP, AUGUSTUS, and TSEBRA. Genome Research 34: 769–777.

Gabriel L, Hoff KJ, Bruna T, Borodovsky M, Stanke M. 2021. TSEBRA: transcript selector for BRAKER. BMC Bioinformatics 22: 566.

Galindo-Martínez CT, Weber M, Avila-Magaña V, Enríquez S, Kitano H, Medina M, Iglesias-Prieto R. 2022. The role of the endolithic alga *Ostreobium* spp. during coral bleaching recovery. Scientific reports 12: 2977.

Garcia-Pichel F, Ramirez-Reinat E, Gao Q. 2010. Microbial excavation of solid carbonates powered by P-type ATPase-mediated transcellular Ca2^+^ transport. *Proceedings of the National Academy of Sciences*, USA 107: 21749–54.

Grabherr MG, Haas BJ, Yassour M, Levin JZ, Thompson DA, et al. 2011. Full-length transcriptome assembly from RNA-Seq data without a reference genome. Nature biotechnology 29: 644–652.

Grass Phylogeny Working Group III, Arthan W, Baker WJ, et al. 2025. A nuclear phylogenomic tree of grasses (Poaceae) recovers current classification despite gene tree incongruence. New Phytologist 245: 818–834.

Guiry MD, Guiry G. 2024. AlgaeBase. World-wide electronic publication, National university of Ireland, Galway.

Guo J, Xu W, Hu Y, Huang J, Zhao Y, Zhang L, Huang C-H, Ma H. 2020. Phylotranscriptomics in Cucurbitaceae reveal multiple whole-genome duplications and key morphological and molecular innovations. Molecular Plant 13: 1117–1133.

Haas BJ, Papanicolaou A, Yassour M, Grabherr M, Blood PD, et al. 2013. De novo transcript sequence reconstruction from RNA-seq using the Trinity platform for reference generation and analysis. Nature Protocols 8: 1494–512.

Hahn MW. 2007. Bias in phylogenetic tree reconciliation methods: implications for vertebrate genome evolution. Genome biology 8: R141.

Hay ME. 1996. Marine chemical ecology: what’s known and what’s next? Journal of Experimental Marine Biology and Ecology 200: 103–134.

Hillis-Colinvaux L. 1984. Systematics of the Siphonales. In: Irvine DEG, John DM (Hrsg.): Systematics of the Green Algae. Academic Press, New York-London, pp. 271–296.

Hoff KJ, Lange S, Lomsadze A, Borodovsky M, Stanke M. 2016. BRAKER1: Unsupervised RNA-Seq-Based Genome Annotation with GeneMark-ET and AUGUSTUS. Bioinformatics 32: 767–9.

Hollants J, Leliaert F, Verbruggen H, Willems A, De Clerck O. 2013. Permanent residents or temporary lodgers: characterizing intracellular bacterial communities in the siphonous green alga *Bryopsis*. Proceedings of the Royal Society B 280: 28020122659.

Hossen R, Courtney M, Sim A, Khan M, Verbruggen H, Bringloe T. 2025. An optimized CTAB method for genomic DNA extraction from green seaweeds (Ulvophyceae). Applications in Plant Sciences 13: e11625.

Hou Z, Ma X, Shi X, Li X, Yang L, Xiao S, De Clerck O, Leliaert F, Zhong B. 2022. Phylotranscriptomic insights into a Mesoproterozoic–Neoproterozoic origin and early radiation of green seaweeds (Ulvophyceae). Nature Communications 13: 1610.

Huisman J. 2015. Algae of Australia: marine benthic algae of north-western Australia. 1. Green and brown algae. Melbourne, Australia: CSIRO Publishing.

Huisman J, Verbruggen H. 2015. Rhipiliaceae, in Huisman J. (ed), Algae of Australia: marine benthic algae of north-western Australia. 1. Green and brown algae. Melbourne, Vic., Australia: CSIRO Publishing, 139–143.

Huisman JM, Verbruggen H. 2023. A morphological and molecular study supports the recognition of *Rhipilia psammophila* sp. nov. and *Rhipilia baculifera* comb. nov. (Halimedaceae, Chlorophyta) from southern Australia. Australian Systematic Botany 36: 427–436.

Iha C, Dougan KE, Varela JA, Avila V, Jackson CJ, et al. 2021. Genomic adaptations to an endolithic lifestyle in the coral-associated alga *Ostreobium*. Current Biology 31: 1393–1402 e5.

Jackson C, Knoll AH, Chan CX, Verbruggen H. 2018. Plastid phylogenomics with broad taxon sampling further elucidates the distinct evolutionary origins and timing of secondary green plastids. Scientific Reports 8: 1523.

Johnson MG, Pokorny L, Dodsworth S, Botigué LR, et al. 2019. A universal probe set for targeted sequencing of 353 nuclear genes from any flowering plant designed using k-medoids clustering. Systematic biology 68: 594–606.

Kalyaanamoorthy S, Minh BQ, Wong TKF, von Haeseler A, Jermiin LS. 2017. ModelFinder: fast model selection for accurate phylogenetic estimates. Nature Methods 14: 587–589.

Kanehisa M, Sato Y, Morishima K. 2016. BlastKOALA and GhostKOALA: KEGG tools for functional characterization of genome and metagenome sequences. Journal of molecular biology 428: 726–731.

Kazi MA, Reddy C, Jha B. 2013. Molecular phylogeny and barcoding of *Caulerpa* (Bryopsidales) based on the tuf A, rbcL, 18S rDNA and ITS rDNA genes. PloS one 8: e82438.

Kellogg EA. 2016. Has the connection between polyploidy and diversification actually been tested? Current opinion in plant biology 30: 25–32.

Kerswell AP. 2006. Global biodiversity patterns of benthic marine algae. Ecology 87: 2479–2488.

Kim D, Paggi JM, Park C, Bennett C, Salzberg SL. 2019. Graph-based genome alignment and genotyping with HISAT2 and HISAT-genotype. Nature Biotechnology 37: 907–915.

Kim GH, Klotchkova TA, Kang Y-M. 2001. Life without a cell membrane: regeneration of protoplasts from disintegrated cells of the marine green alga *Bryopsis plumosa*. Journal of Cell Science 114: 2009–2014.

Kohli A, Narciso JO, Miro B, Raorane M. 2012. Root proteases: reinforced links between nitrogen uptake and mobilization and drought tolerance. Physiologia Plantarum 145: 165–179.

Kornmann P, Sahling PH. 1980. *Ostreobium quekettii* (Codiales, Chlorophyta). Helgoländer Meeresuntersuchungen 34: 115–22.

Kondrashov FA. 2012. Gene duplication as a mechanism of genomic adaptation to a changing environment. Proceedings of the Royal Society B: Biological Sciences 279: 5048–5057.

Kooistra WH. 2002. Molecular phylogenies of Udoteaceae (Bryopsidales, Chlorophyta) reveal nonmonophyly for *Udotea*, *Penicillus* and *Chlorodesmis*. Phycologia 41: 453–462.

Kuznetsov D, Tegenfeldt F, Manni M, Seppey M, Berkeley M, Kriventseva EV, Zdobnov EM. 2023. OrthoDB v11: annotation of orthologs in the widest sampling of organismal diversity. Nucleic Acids Research 51: 445–451.

Lagourgue L, Payri CE. 2022. Large-scale diversity reassessment, evolutionary history, and taxonomic revision of the green macroalgae family Udoteaceae (Bryopsidales, Chlorophyta). Journal of Systematics and Evolution 60: 101–127.

Lam DW, Zechman FW. 2006. Phylogenetic analysis of the Bryopsidales (Ulvophyceae, Chlorophyta) based on rubisco large subunit gene sequences1. Journal of Phycology 42: 669–678.

Leliaert F, Verbruggen H, D’hondt S, López-Bautista JM, De Clerck O. 2014. The forgotten genus *Pseudoderbesia* (Bryopsidales, Chlorophyta). Cryptogamie Algologie 35: 207–219.

Leslie AB, Mander L. 2025. Genomic correlates of vascular plant reproductive complexity and the uniqueness of angiosperms. New Phytologist 245: 1733–1745.

Levin DA. 2019. Why polyploid exceptionalism is not accompanied by reduced extinction rates. Plant Systematics and Evolution 305: 1–1.

Li H, Handsaker B, Wysoker A, Fennell T, Ruan J, et al. 2009. The sequence alignment/map format and SAMtools. bioinformatics 25: 2078–2079.

Li W, Godzik A. 2006. Cd-hit: a fast program for clustering and comparing large sets of protein or nucleotide sequences. Bioinformatics 22: 1658–1659.

Li X, Hou Z, Xu C, Shi X, Yang L, Lewis LA, Zhong B. 2021. Large phylogenomic data sets reveal deep relationships and trait evolution in chlorophyte green algae. Genome Biology and Evolution 13: p.evab101.

Li Z, Defoort J, Tasdighian S, Maere S, Van de Peer Y, De Smet R. 2016. Gene duplicability of core genes is highly consistent across all angiosperms. The Plant Cell 28: 326–44.

Lu J, Xu Y, Wang J, Singer SD, Chen G. 2020. The role of triacylglycerol in plant stress response. Plants (Basel) 9: p.472.

Ma X, Vanneste S, Chang J, Ambrosino L, Barry K, et al. 2024. Seagrass genomes reveal ancient polyploidy and adaptations to the marine environment. Nature plants 10: 240–255.

Magadum S, Banerjee U, Murugan P, Gangapur D, Ravikesavan R. 2013. Gene duplication as a major force in evolution. Journal of genetics 92: 155–161.

Marcelino VR, Verbruggen H. 2016. Multi-marker metabarcoding of coral skeletons reveals a rich microbiome and diverse evolutionary origins of endolithic algae. Scientific Report 6: 31508.

Mayrose I, Zhan SH, Rothfels CJ, Magnuson-Ford K, Barker MS, Rieseberg LH, Otto SP. 2011.. Recently formed polyploid plants diversify at lower rates. Science 333: 1257–1257.

Mayrose I, Zhan SH, Rothfels CJ, Arrigo N, Barker MS, Rieseberg LH, Otto SP. 2014. Methods for studying polyploid diversification and the dead end hypothesis: a reply to Soltis et al.(2014). New Phytologist 206:27–35.

Meinesz A, Benichou L, Blachier J, Komatsu T, Lemée R, Molenaar H, Mari X. 1995. Variations in the structure, morphology and biomass of *Caulerpa taxifolia* in the Mediterranean Sea. Botanica Marina 38: 499–508.

Meinita MDN, Harwanto D, Choi JS. 2022. A concise review of the bioactivity and pharmacological properties of the genus *Codium* (Bryopsidales, Chlorophyta). Journal of Applied Phycology 34: 2827–2845.

Mandel JR, Barker MS, Bayer RJ, Dikow RB, et al. 2017. The compositae tree of life in the age of phylogenomics. Journal of Systematics and Evolution 55: 405–410.

Minh BQ, Nguyen MA, von Haeseler A. 2013. Ultrafast approximation for phylogenetic bootstrap. Molecular Biology and Evolution 30: 1188–95.

Morgulis A, Gertz EM, Schaffer AA, Agarwala R. 2006. WindowMasker: window-based masker for sequenced genomes. Bioinformatics 22: 134–41.

Ochiai KK, Hanawa D, Ogawa HA, Tanaka H, et al. 2024. Genome sequence and cell biological toolbox of the highly regenerative, coenocytic green feather alga *Bryopsis*. The Plant Journal 119: 1091–1111.

Ohno S. 1970. Ohno S. Evolution by gene duplication. Springer-Verlag.

One thousand plant transcriptomes and the phylogenomics of green plants. Nature 574, no. 7780 (2019): 679–685.

Otto SP. 2007. The evolutionary consequences of polyploidy. Cell 131: 452–462.

Panchy N, Lehti-Shiu M, Shiu SH.2016. Evolution of gene duplication in plants. Plant physiology 171: 2294–316.

Pandit MK, Pocock MJ, Kunin WE. 2011. Ploidy influences rarity and invasiveness in plants. Journal of Ecology 99: 1108–1115.

Pan-Utai W, Satmalee P, Saah S, Paopun Y, Tamtin M. 2023. Brine-processed *Caulerpa lentillifera* macroalgal stability: physicochemical, nutritional and microbiological properties. Life 13: 2112.

Paul VJ, Fenical W. 1983. Isolation of halimedatrial: chemical defense adaptation in the calcareous reef-building alga *Halimeda*. Science 221: 747–749.

Piazzi L, Balata D, Bulleri F, Gennaro P, Ceccherelli G. 2016. The invasion of *Caulerpa cylindracea* in the Mediterranean: the known, the unknown and the knowable. Marine biology 163: 1–14.

Pospíšil P. 2016. Production of reactive oxygen species by photosystem II as a response to light and temperature stress. Frontiers in plant science 7: 1950.

Pryszcz LP, Gabaldon T. 2016. Redundans: an assembly pipeline for highly heterozygous genomes. Nucleic Acids Research 44: e113.

Pushpakumara BLDU, Tandon K, Willis A, Verbruggen H. 2023. The bacterial microbiome of the coral skeleton algal symbiont *Ostreobium* shows preferential associations and signatures of phylosymbiosis. Microbial Ecology 86: 2032–2046.

Ranjan A, Townsley BT, Ichihashi Y, Sinha NR, Chitwood DH. 2015. An intracellular transcriptomic atlas of the giant coenocyte *Caulerpa taxifolia*. PLoS genetics 11: e1004900.

Ren R, Wang H, Guo C, Zhang N, Zeng L, Chen Y, Ma H, Qi J. 2018. Widespread whole genome duplications contribute to genome complexity and species diversity in angiosperms. Molecular Plant 11: 414–428.

Rensing SA. 2014. Gene duplication as a driver of plant morphogenetic evolution. Current opinion in plant biology 17: 43–48.

Roberts IN, Caputo C, Criado MV, Funk C. 2012. Senescence-associated proteases in plants. Physiol Plant 145: 130–9.

Sayyari E, Mirarab S. 2016. Fast coalescent-based computation of local branch support from quartet frequencies. Molecular Biology and Evolution 33: 1654–68.

Schaller A. 2004. A cut above the rest: the regulatory function of plant proteases. Planta 220: 183–97.

Schranz ME, Mohammadin S, Edger PP. 2012. Ancient whole genome duplications, novelty and diversification: the WGD Radiation Lag-Time Model. Current Opinion in Plant Biology 15: 147–53.

Seppey M, Manni M, Zdobnov EM. 2019. BUSCO: assessing genome assembly and annotation completeness. In Gene prediction: methods and protocols. New York, NY: Springer New York, pp. 227–245.

Simmons MP. 2000. A fundamental problem with amino-acid-sequence characters for phylogenetic analyses. Cladistics 16: 274–282.

Smit A, Hubley R, Green P. 2013. RepeatMasker Open-4.0., 2013. Available online: http://www.repeatmasker.org [Accessed: June 2024]

Soltis DE, Segovia-Salcedo MC, Jordon-Thaden I, Majure L, et al. 2014. Are polyploids really evolutionary dead-ends (again)? A critical reappraisal of Mayrose et al.(2011). New Phytologist 202: 1105–1117.

Steenwyk JL, Buida III TJ, Labella AL, Li Y, Shen X-X, Rokas A. 2021. PhyKIT: a broadly applicable UNIX shell toolkit for processing and analyzing phylogenomic data. Bioinformatics 37: 2325–2331.

Sun P, Jiao B, Yang Y, Shan L, Li T, Li X, Xi Z, Wang X, Liu J. 2022. WGDI: a user-friendly toolkit for evolutionary analyses of whole-genome duplications and ancestral karyotypes. Molecular plant 15:1841–51.

Szöllősi GJ, Tannier E, Daubin V, Boussau B. 2015. The inference of gene trees with species trees. Systematic biology 64: e42–62.

Tandon K, Pasella MM, Iha C, Ricci F, Hu J, O’Kelly CJ, Medina M, Kühl M, Verbruggen H. Every refuge has its price: *Ostreobium* as a model for understanding how algae can live in rock and stay in business. 2023. Seminars in cell & developmental biology 134: 27–36.

Te Beest M, Le Roux JJ, Richardson DM, Brysting AK, Suda J, Kubešová M, Pyšek P. 2012. The more the better? The role of polyploidy in facilitating plant invasions. Annals of botany 109: 19–45.

Terada R, Tanaka T, Uchimura M. 2012. Morphology and distribution of *Caulerpa lentillifera* J. Agardh (Chrolophyceae) in Japanese waters, including the first record from southern Kyushu and northern Ryukyu Islands. The journal of Japanese botany 87: 260–267

Tiley GP, Barker MS, Burleigh JG. 2018. Assessing the performance of Ks plots for detecting ancient whole genome duplications. Genome biology and evolution 10: 2882–98.

Timilsena PR, Wafula EK, Barrett CF, Ayyampalayam S, et al. 2022. Phylogenomic resolution of order-and family-level monocot relationships using 602 single-copy nuclear genes and 1375 BUSCO genes. Frontiers in Plant Science 13: 876779.

Treier UA, Broennimann O, Normand S, Guisan A, Schaffner U, Steinger T, Müller-Schärer H. 2009. Shift in cytotype frequency and niche space in the invasive plant *Centaurea maculosa*. Ecology 90: 1366–1377.

Tuteja N, Ahmad P, Panda BB, Tuteja R. 2009. Genotoxic stress in plants: shedding light on DNA damage, repair and DNA repair helicases. Mutation Research/Reviews in Mutation Research 681: 134–149.

Ukabi S, Dubinsky Z, Steinberger Y, Israel A. 2013. Temperature and irradiance effects on growth and photosynthesis of *Caulerpa* (Chlorophyta) species from the eastern Mediterranean. Aquatic botany 104: 106–110.

van de Peer Y, Maere S, Meyer A. 2009. The evolutionary significance of ancient genome duplications. Nature Reviews Genetics 10: 725–32.

van Hoek MJ, Hogeweg P. 2009. Metabolic adaptation after whole genome duplication. Molecular Biology and Evolution 26: 2441–53.

Varela-Álvarez E, Balau AC, Marbà N, Afonso-Carrillo J, Duarte CM, Serrão EA. 2015. Genetic diversity and biogeographical patterns of *Caulerpa prolifera* across the Mediterranean and Mediterranean/Atlantic transition zone. Marine Biology 162: 557–569.

Verbruggen H, Ashworth M, LoDuca ST, Vlaeminck C, et al. 2009a. A multi-locus time-calibrated phylogeny of the siphonous green algae. Molecular phylogenetics and evolution 50: 642–653.

Verbruggen H, Clerck OD, Schils T, Kooistra WH, Coppejans E. 2005. Evolution and phylogeography of *Halimeda* section *Halimeda* (Bryopsidales, Chlorophyta). Molecular Phylogenetics and Evolution 37: 789–803.

Verbruggen H, Leliaert F, Maggs CA, Shimada S, Schils T, et al. 2007. Species boundaries and phylogenetic relationships within the green algal genus *Codium* (Bryopsidales) based on plastid DNA sequences. Molecular phylogenetics and evolution 44: 240–254.

Verbruggen H, Marcelino VR, Guiry MD, Cremen MCM, Jackson CJ. 2017. Phylogenetic position of the coral symbiont *Ostreobium* (Ulvophyceae) inferred from chloroplast genome data. Journal of Phycology 53: 790–803.

Verbruggen H, Tyberghein L, Pauly K, Vlaeminck C, Nieuwenhuyze KV, Kooistra WH, Leliaert F, Clerck OD. 2009b. Macroecology meets mamulti-locus time-calibrated phylogeny of the siphonous green algcroevolution: evolutionary niche dynamics in the seaweed *Halimeda*. Global Ecology and Biogeography 18: 393–405.

Verbruggen H, Vlaeminck C, Sauvage T, Sherwood AR, Leliaert F, De Clerck O. 2009c. Phylogenetic analysis of *Pseudochlorodesmis* strains reveals cryptic diversity above the family level in the siphonous green algae (Bryopsidales, Chlorophyta)(1). Journal of Phycology 45: 726–31.

Vernot B, Stolzer M, Goldman A, Durand D. 2008. Reconciliation with non-binary species trees. Journal of computational biology 15: 981–1006.

Verlaque M, Durand C, Huisman JM, Boudouresque C-F, Le Parco Y. 2003. On the identity and origin of the Mediterranean invasive *Caulerpa racemosa* (Caulerpales, Chlorophyta). European Journal of Phycology 38: 325–339.

Vogel K. 1993. Bioeroders in fossil reefs. Facies 28: 109–113.

Vogel K, Brett CE. 2009. Record of microendoliths in different facies of the Upper Ordovician in the Cincinnati Arch region USA: the early history of light-related microendolithic zonation. Palaeogeography, Palaeoclimatology, Palaeoecology 281: 1–24.

Vroom PS, Smith CM. 2003. Life without cells. Biologist 50: 222–226.

Vroom PS, Smith CM, Keeley SC. 1998. Cladistics of the Bryopsidales: a preliminary analysis. Journal of phycology 34: 351–360.

Walters LJ, Brown KR, Stam WT, Olsen JL. 2006. E-commerce and *Caulerpa*: unregulated dispersal of invasive species. Frontiers in Ecology and the Environment 4: 75–79.

Wan J, Oduor AM, Pouteau R, Wang B, Chen L, Yang B, Yu F, Li J. 2020. Can polyploidy confer invasive plants with a wider climatic tolerance? A test using Solidago canadensis. Ecology and Evolution 10: 5617–5630.

Wickham H. 2016. ggplot2 : Elegant Graphics for Data Analysis, 2nd edition [eBook]. Cham: Springer International Publishing : Imprint: Springer.

Williams SL, Grosholz ED. 2002. Preliminary reports from the *Caulerpa taxifolia* invasion in southern California. Marine Ecology Progress Series 233: 307–310.

Woolcott GW, Knöller K, King RJ. 2000. Phylogeny of the Bryopsidaceae (Bryopsidales, Chlorophyta): cladistic analyses of morphological and molecular data. Phycologia 39: 471–481.

Wu S, Han B, Jiao Y. 2020. Genetic contribution of paleopolyploidy to adaptive evolution in angiosperms. Molecular plant 13: 59–71.

Xu P, Liu X, Ke L, Li K, Wang W, Jiao Y. 2025. The genomic insights of intertidal adaptation in *Bryopsis corticulans*. New Phytologist 246:1691–709.

Xu S, Guo Z, Feng X, Shao S, Yang Y, Li J, Zhong C, He Z, Shi S. 2023. Where whole-genome duplication is most beneficial: Adaptation of mangroves to a wide salinity range between land and sea. Molecular Ecology 32: 460–475.

Yant L, Bomblies K. 2015. Genome management and mismanagement—cell-level opportunities and challenges of whole-genome duplication. Genes and development 29: 2405–2419.

Yang Y, Moore MJ, Brockington SF, Mikenas J, Olivieri J, Walker JF, Smith SA. 2018. Improved transcriptome sampling pinpoints 26 ancient and more recent polyploidy events in Caryophyllales, including two allopolyploidy events. New Phytologist 217: 855–870.

Yang Y, Moore MJ, Brockington SF, Soltis DE, et al. 2015. Dissecting molecular evolution in the highly diverse plant clade caryophyllales using transcriptome sequencing. Molecular Biology and Evolution 32: 2001–14.

Yang X, Hu R, Yin H, Jenkins J, Shu S, et al. 2017. The Kalanchoë genome provides insights into convergent evolution and building blocks of crassulacean acid metabolism. Nature communications 8: 1899.

Yildiz G, Dere Ş. 2015. The effects of elevated-CO2 on physiological performance of *Bryopsis plumosa*. Acta Oceanologica Sinica 34: 125–129.

Zhang F, Ding Y, Zhu CD, Zhou X, Orr MC, Scheu S, Luan YX. 2019. Phylogenomics from low-coverage whole-genome sequencing. Methods in Ecology and Evolution 10: 507–517.

Zhang H, Wang X, Qu M, Yu H, Yin J, Liu X, et al. 2024. Genome of *Halimeda opuntia* reveals differentiation of subgenomes and molecular bases of multinucleation and calcification in algae. Proceedings of the National Academy of Sciences, USA 121: e2403222121.

Zhang X, Han W, Fan X, Wang Y, Xu D, Sun K, Wang W, Zhang Y, Ma J, Ye N. 2023. Gene duplication and functional divergence of new genes contributed to the polar acclimation of Antarctic green algae. Marine Life Science and Technology 5: 511–524.

