## supplementary materials for "Extensive nuclear datasets resolve the phylogeny of siphonous green algae and identify genome duplications as a contributing factor to evolutionary adaptations"

### Supporting Information

#### Coalescence (ASTRAL) Trees

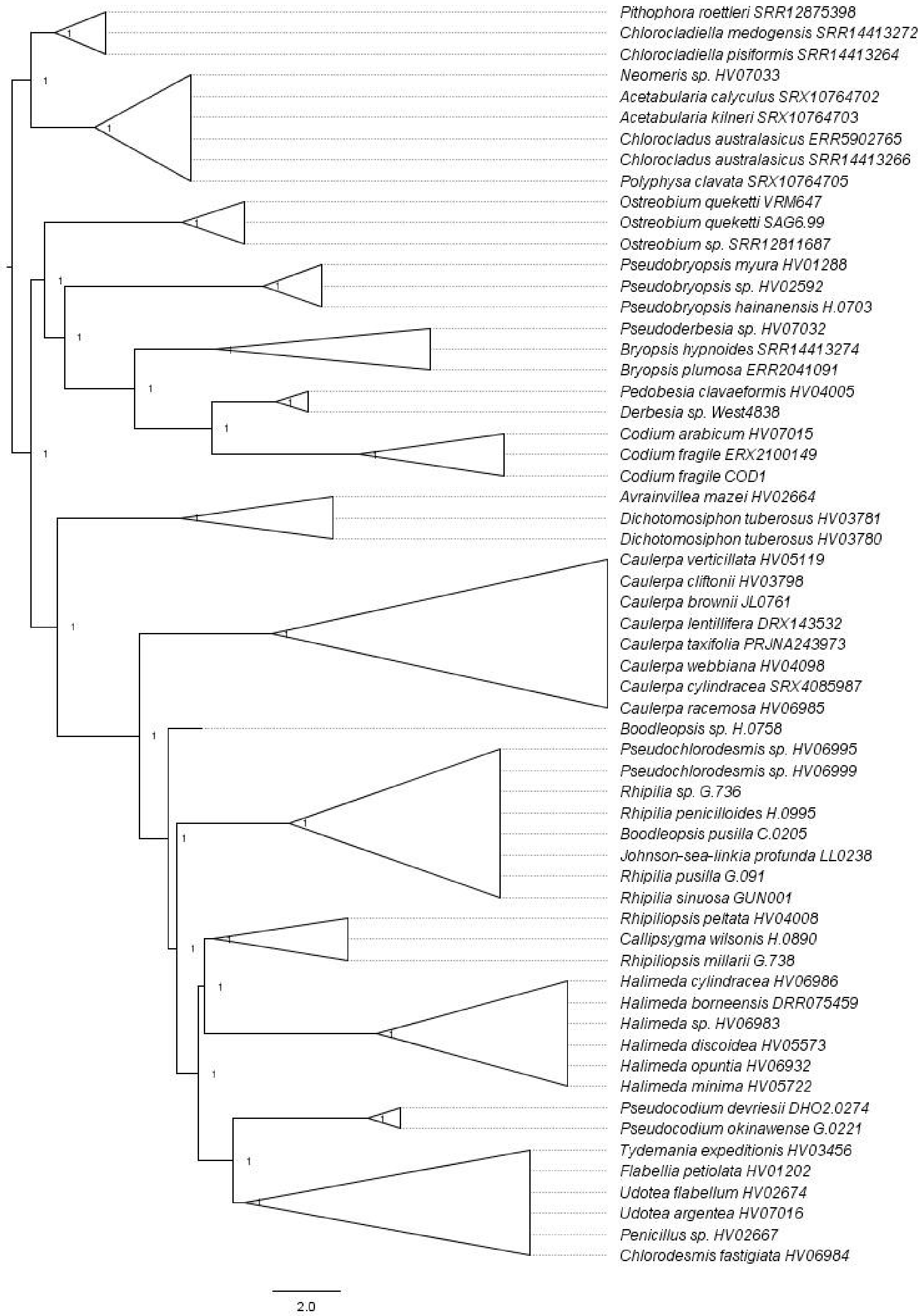


**Fig. S1** ASTRAL species tree inferred from nucleotide-based gene trees, with Dasycladales and Cladophorales as outgroups.

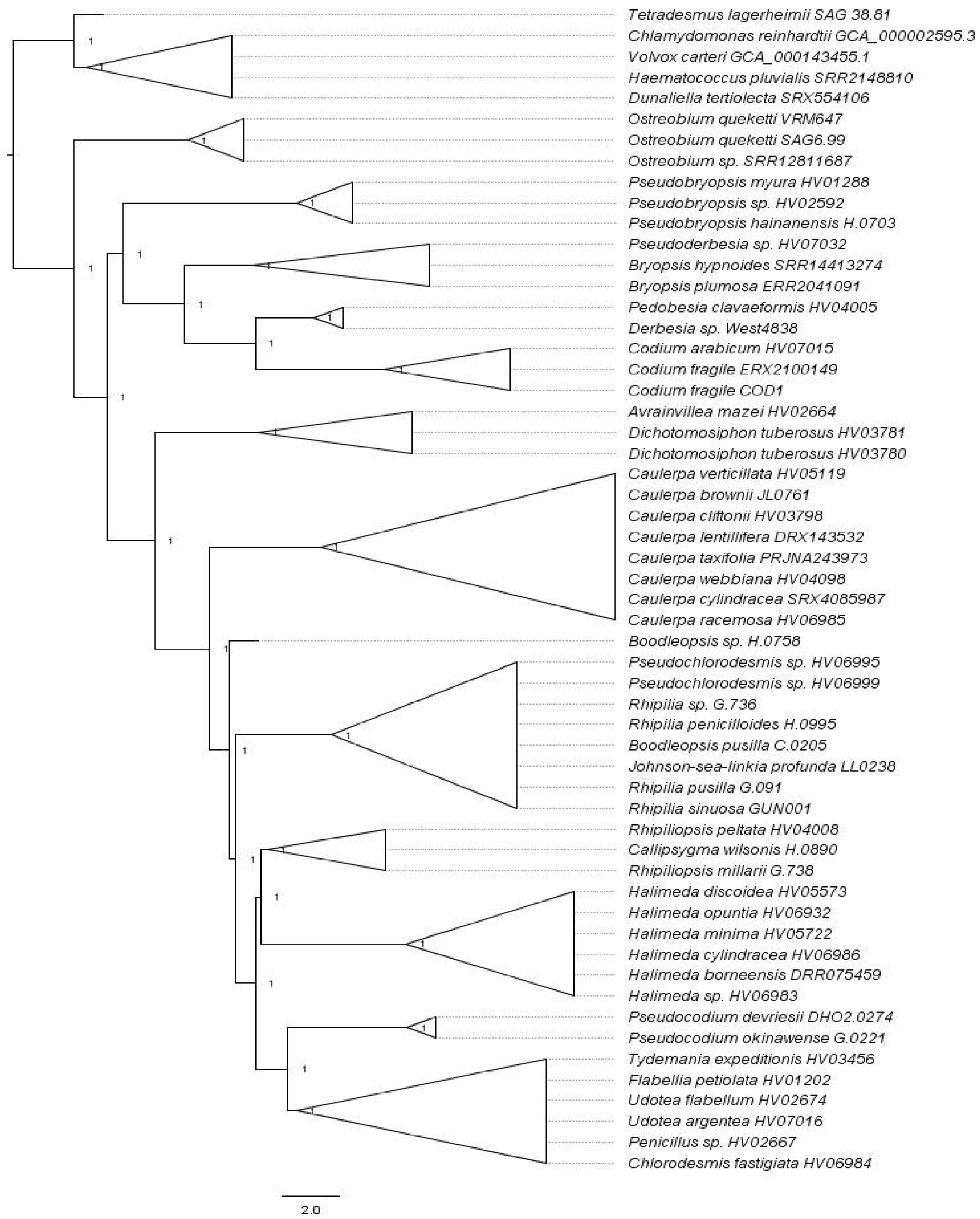


**Fig. S2** ASTRAL species tree inferred from nucleotide-based gene trees, with Chlorophyceae as the outgroup.

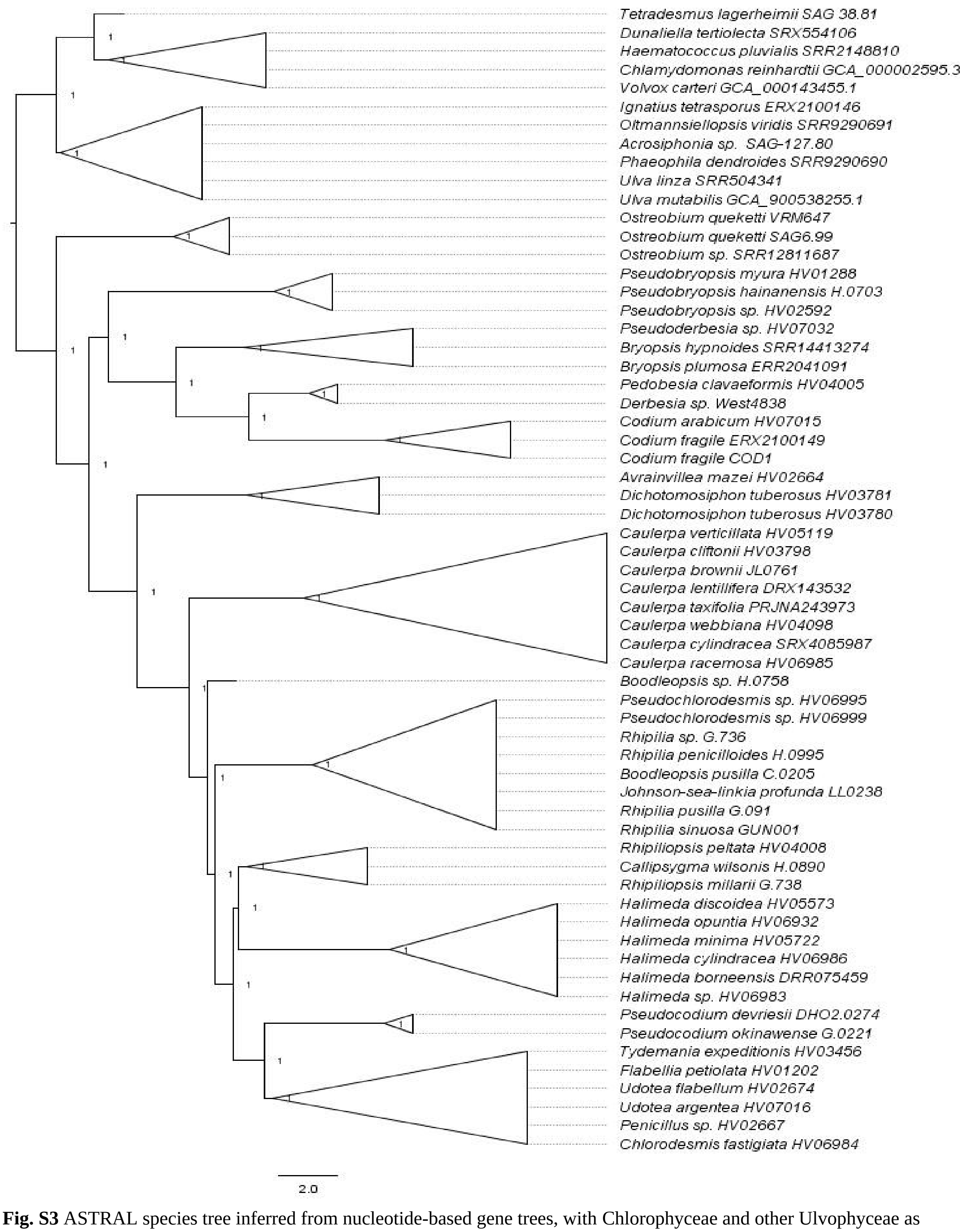


**Fig. S3** ASTRAL species tree inferred from nucleotide-based gene trees, with Chlorophyceae and other Ulvophyceae as outgroups.

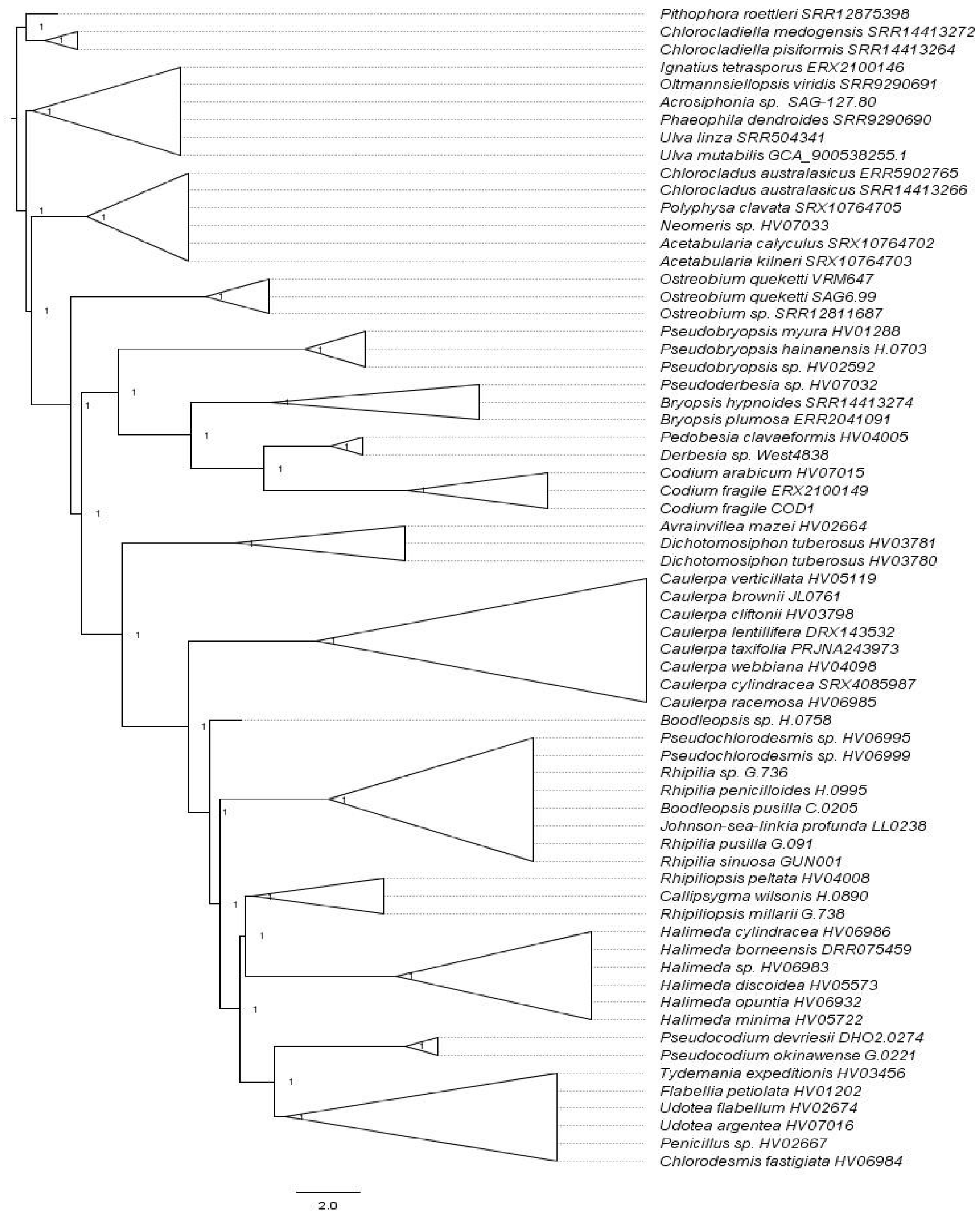


**Fig. S4** ASTRAL species tree inferred from nucleotide-based gene trees, with Dasycladales, Cladophorales and other Ulvophyceae as outgroups.

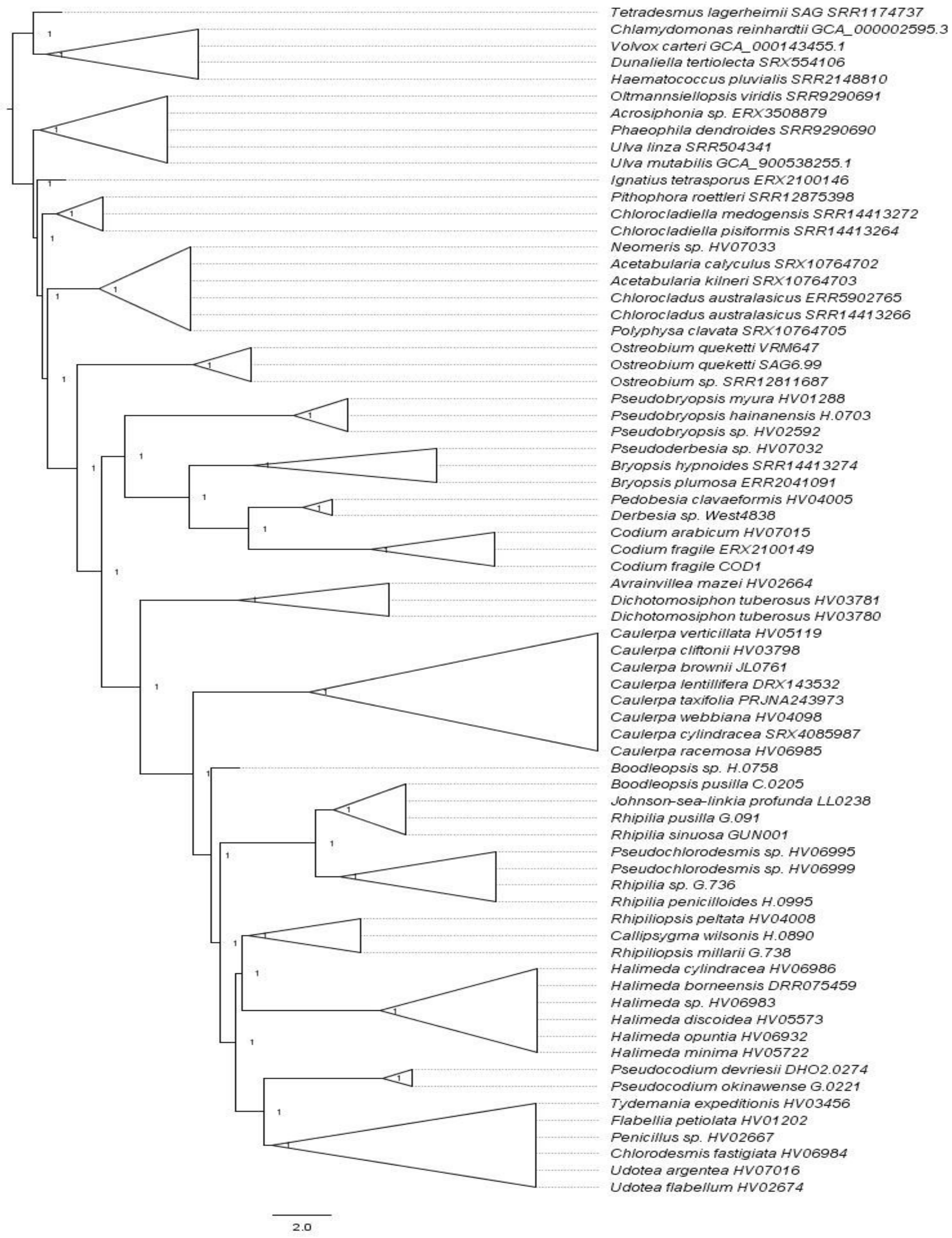


**Fig. S5** ASTRAL species tree inferred from nucleotide-based gene trees, with Dasycladales, Cladophorales, Chlorophyceae and other Ulvophyceae as outgroups.

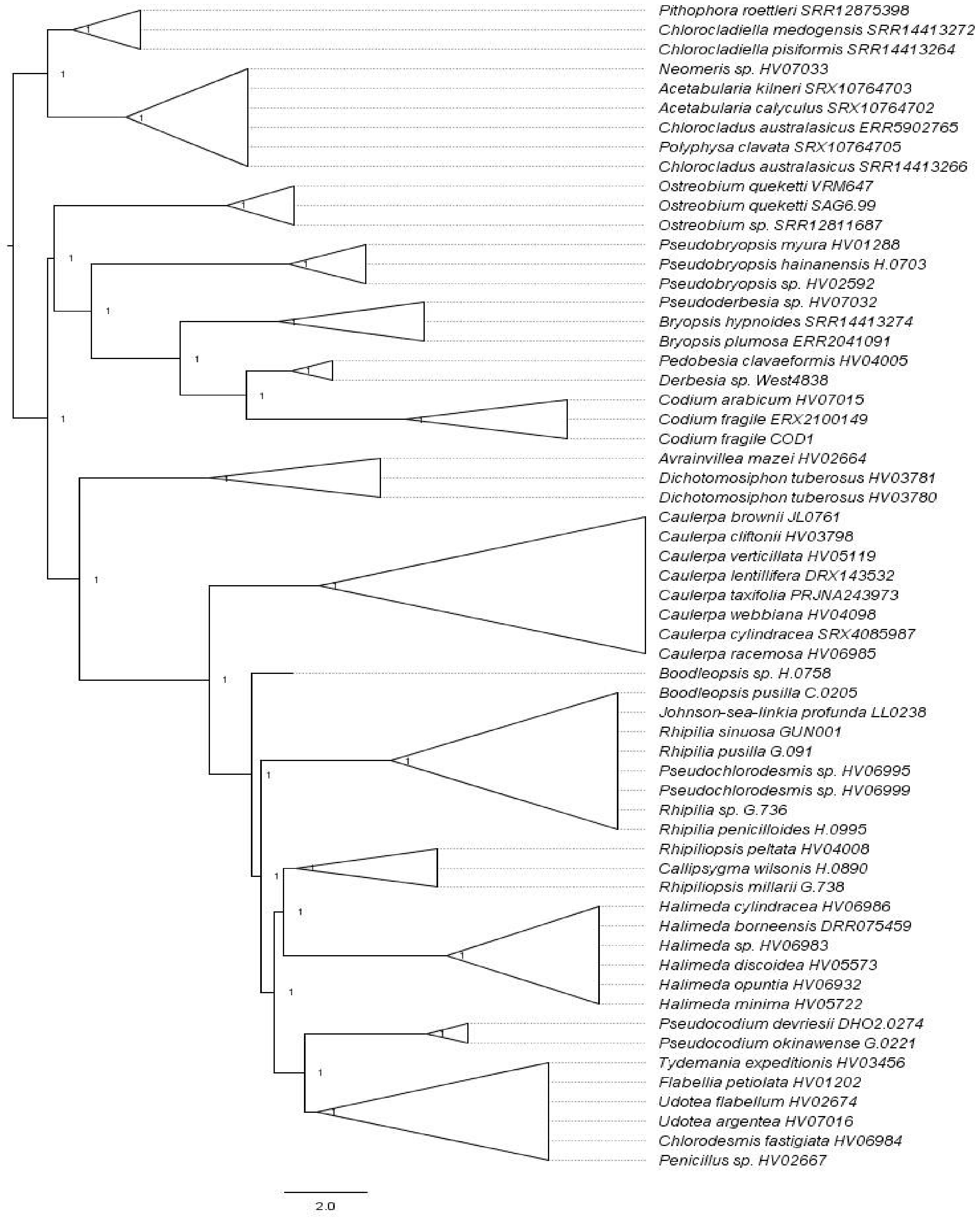


**Fig. S6** ASTRAL species tree inferred from amino acid-based gene trees, with Dasycladales and Cladophorales as outgroups.

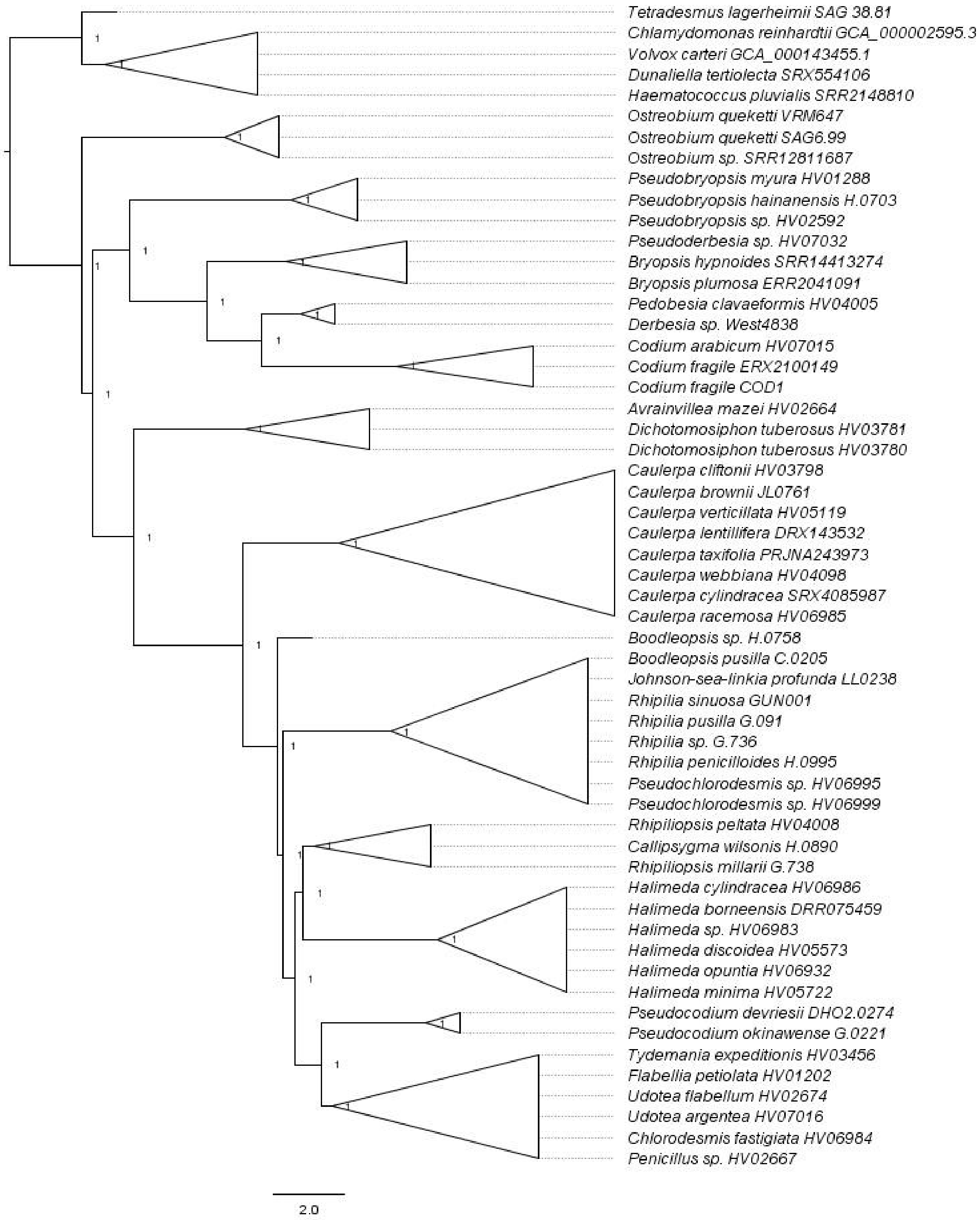


**Fig. S7** ASTRAL species tree inferred from amino acid-based gene trees, with Chlorophyceae as the outgroup.

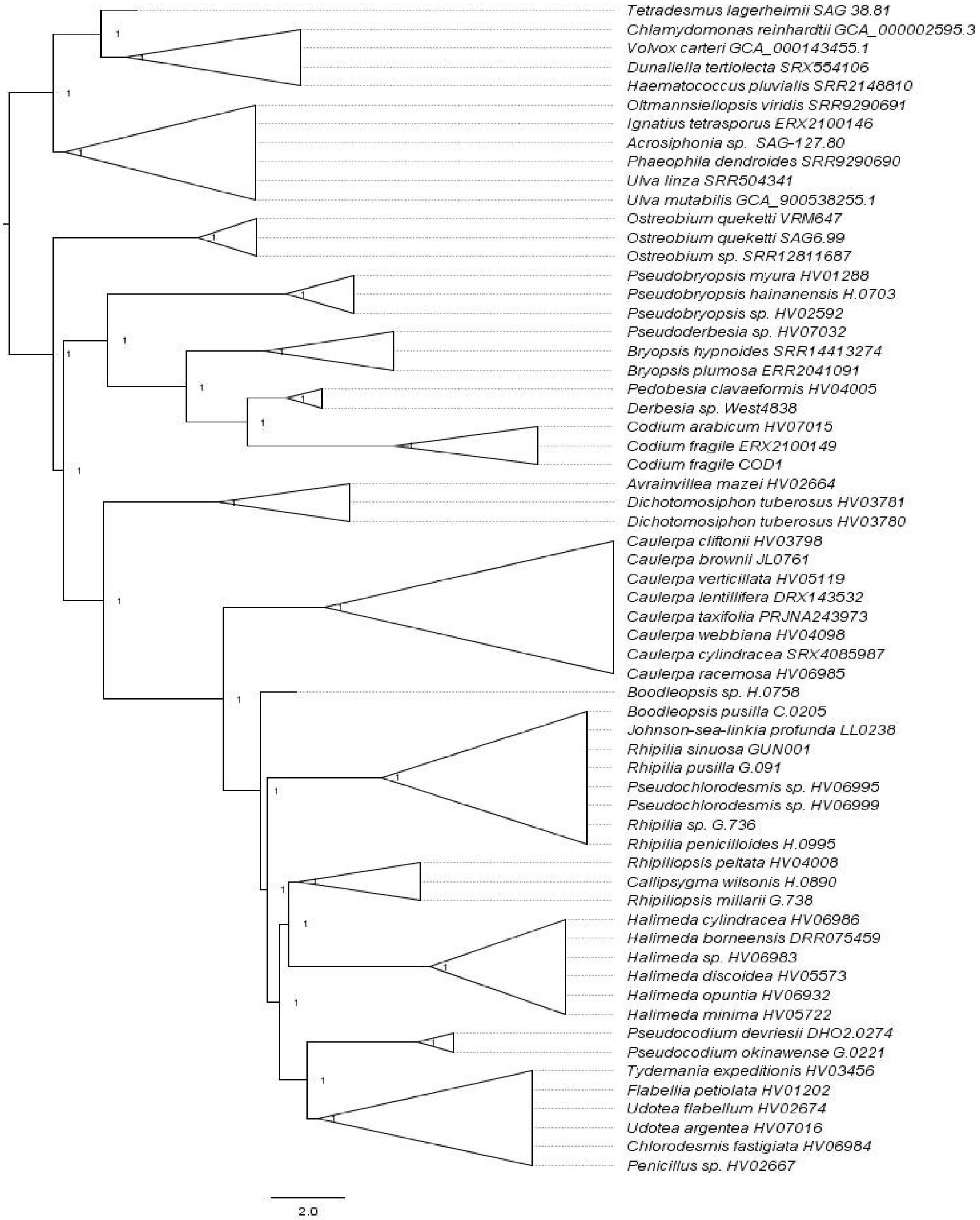


**Fig. S8** ASTRAL species tree inferred from amino acid-based gene trees, with Chlorophyceae and other Ulvophyceae as outgroups.

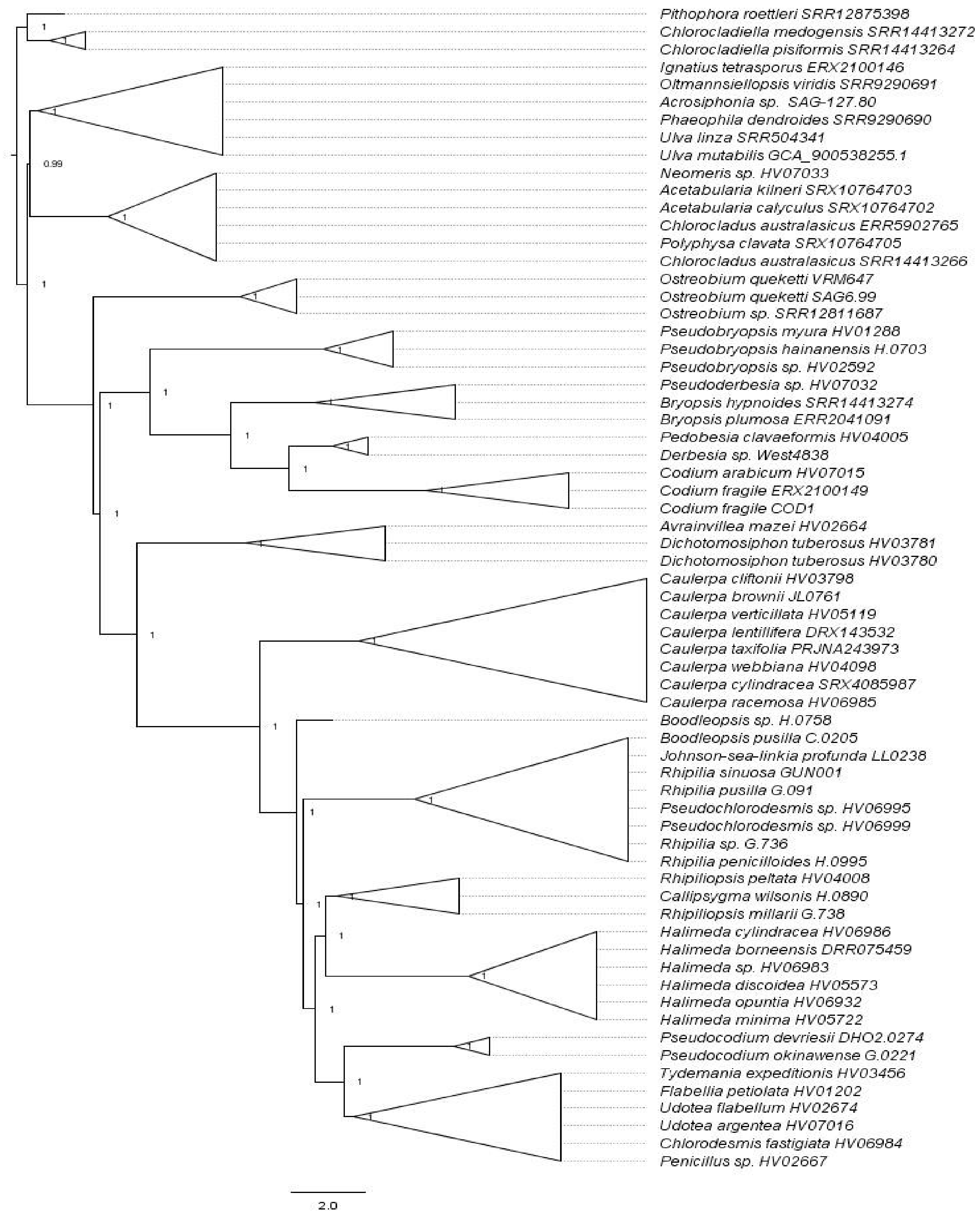


**Fig. S9** ASTRAL species tree inferred from amino acid-based gene trees, with Dasycladales, Cladophorales and other Ulvophyceae as outgroups.

.

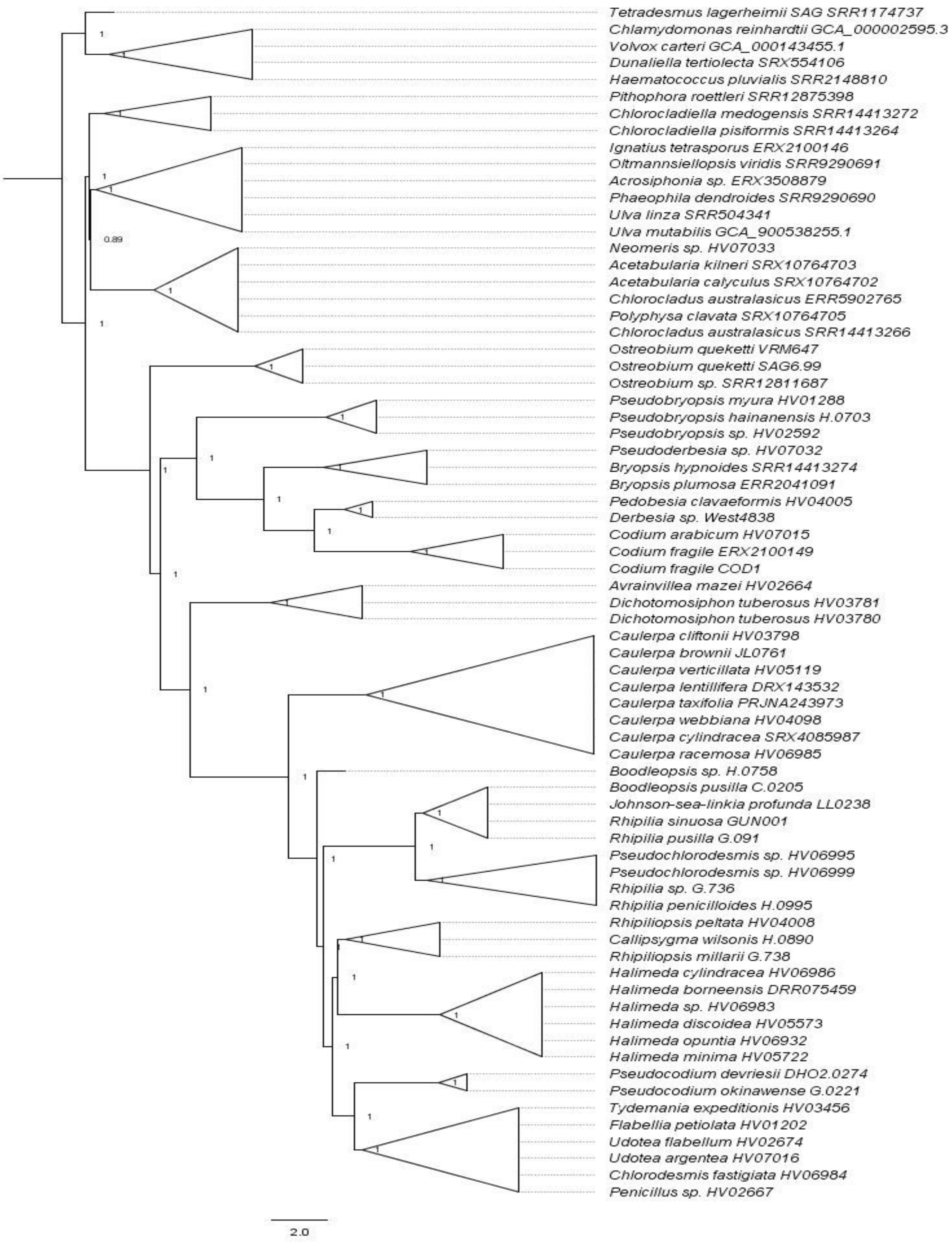


**Fig. S10** ASTRAL species tree inferred from amino acid-based gene trees, with Dasycladales, Cladophorales, Chlorophyceae and other Ulvophyceae as outgroups.

**Concatenation trees**

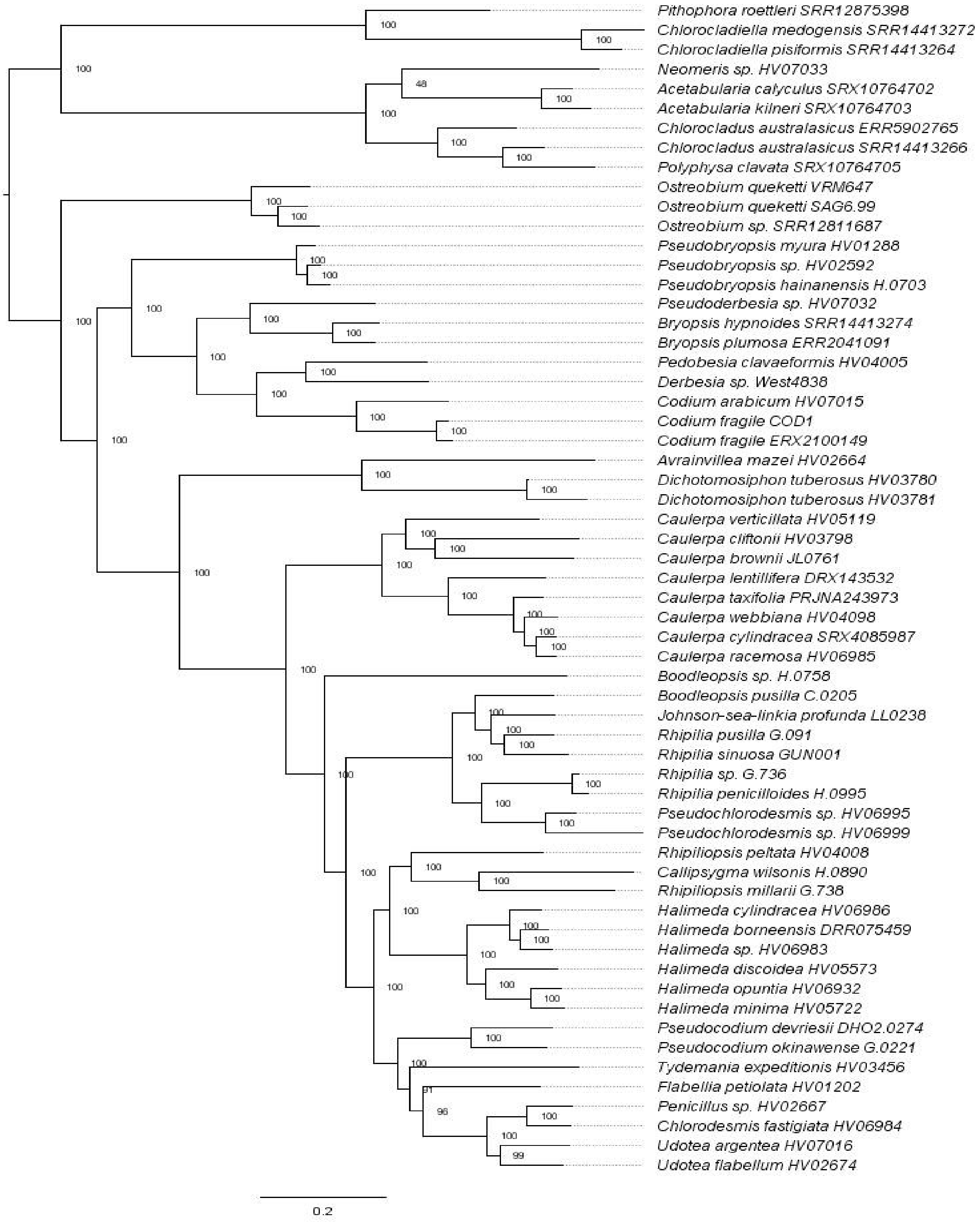


**Fig. S11** Maximum likelihood tree of the concatenated nucleotide alignment, using the model suggested by MFP, and with Dasycladales and Cladophorales as outgroups.

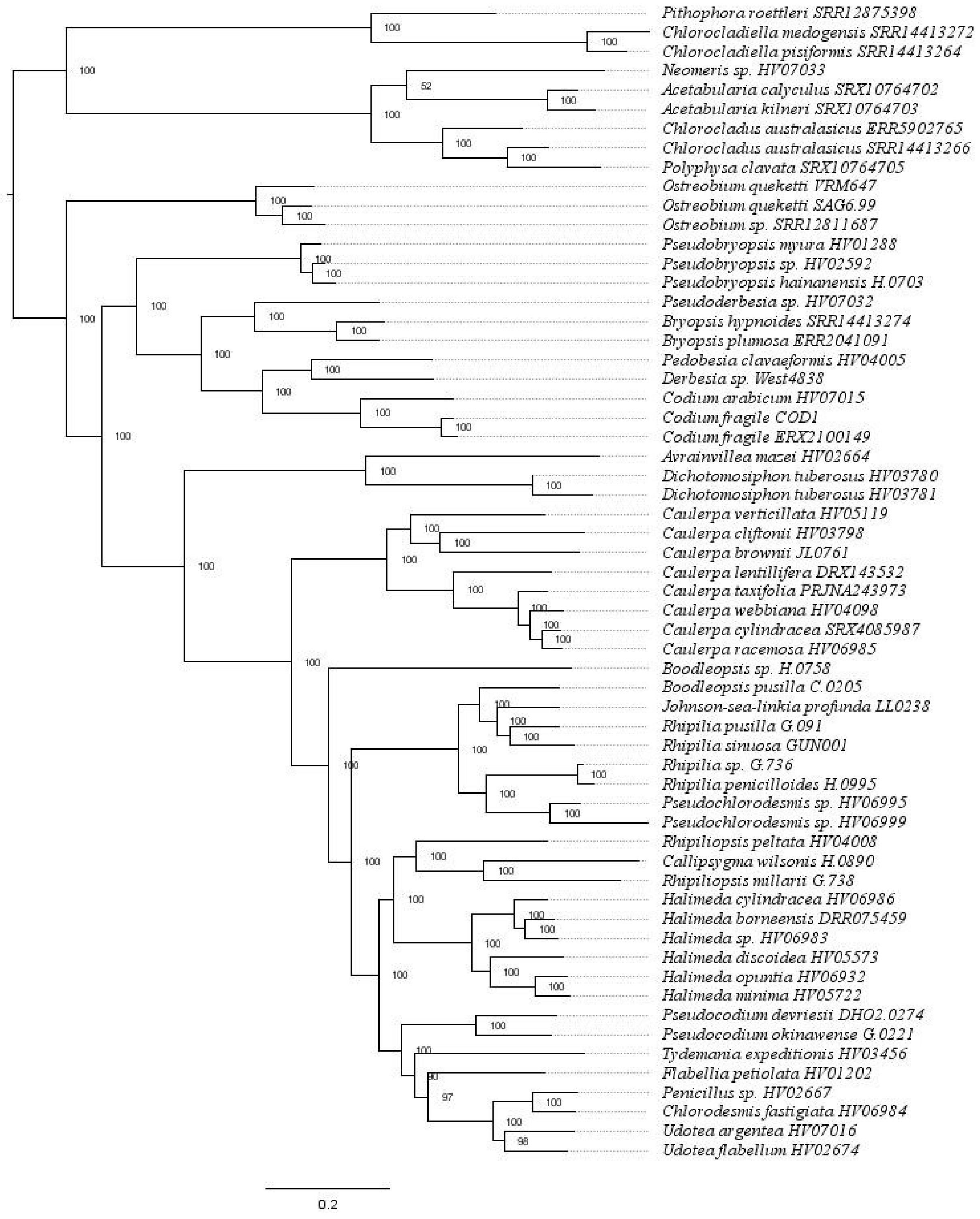


**Fig. S12** Maximum likelihood tree of the concatenated nucleotide alignment, using the GTR+F+I+R7 model, and with Dasycladales and Cladophorales as outgroups.

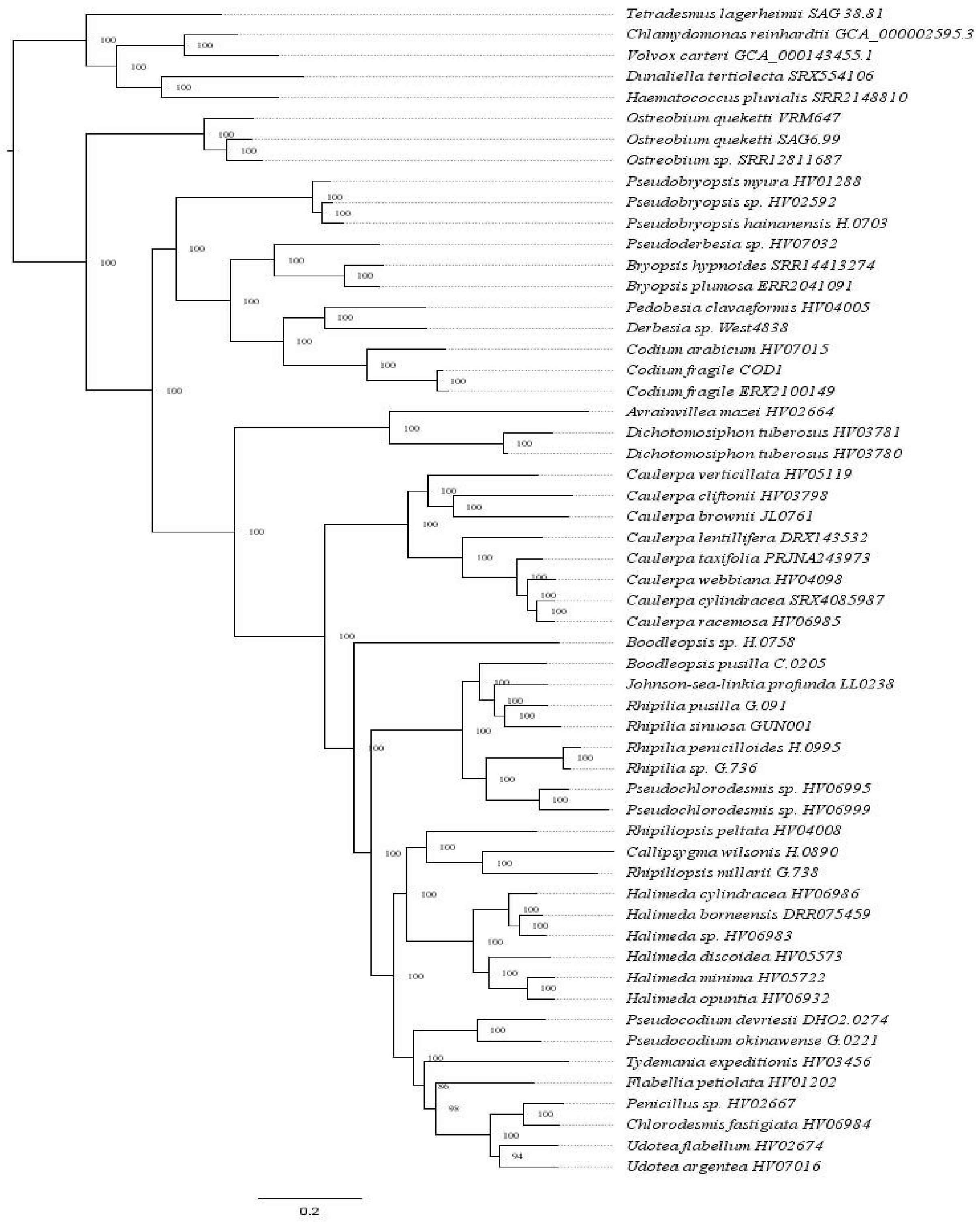


**Fig. S13** Maximum likelihood tree of the concatenated nucleotide alignment, using the GTR+F+I+R6 model, and with Chlorophyceae as the outgroup.

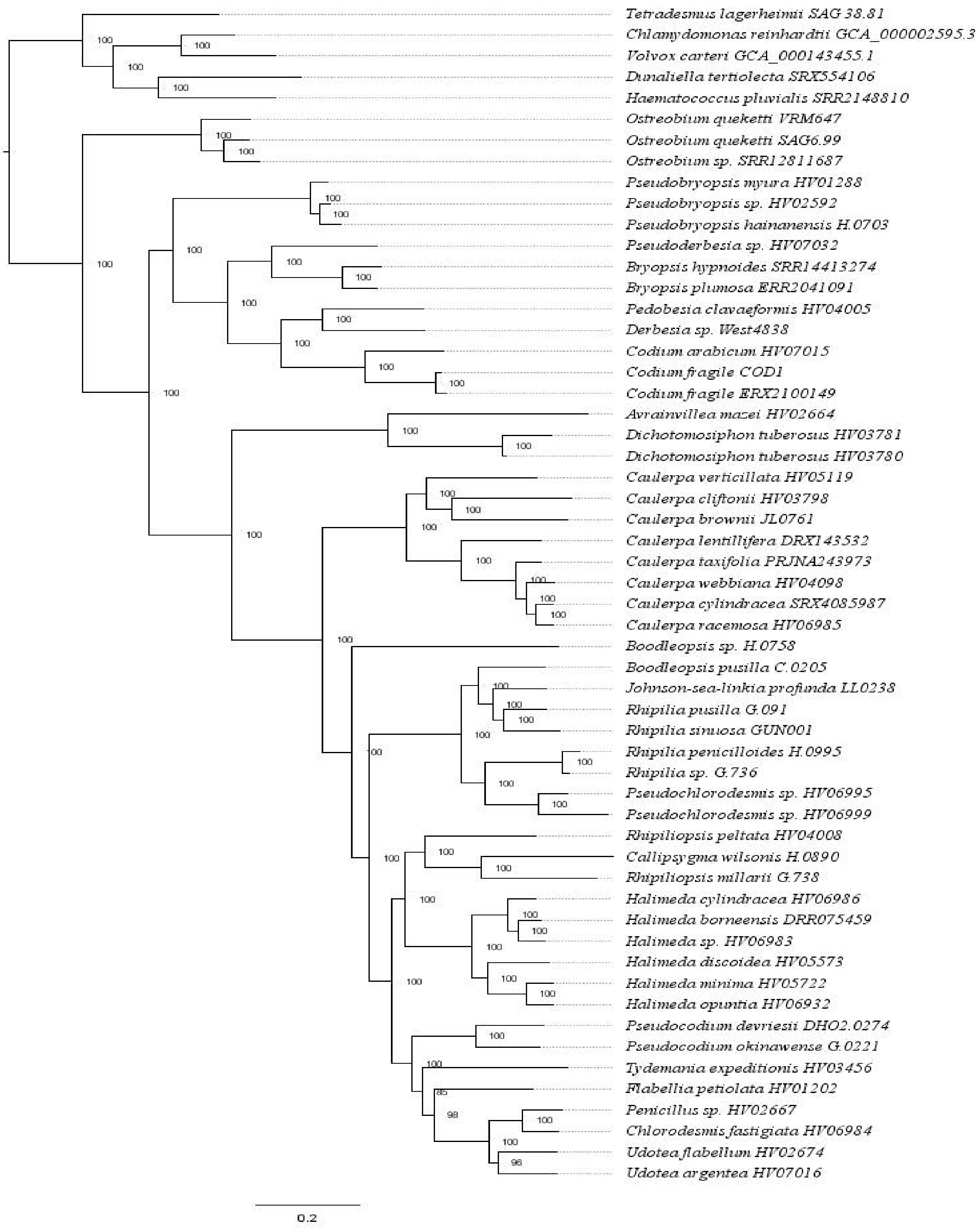


**Fig. S14** Maximum likelihood tree of the concatenated nucleotide alignment, using the GTR+F+I+R7 model, and with Chlorophyceae as outgroup.

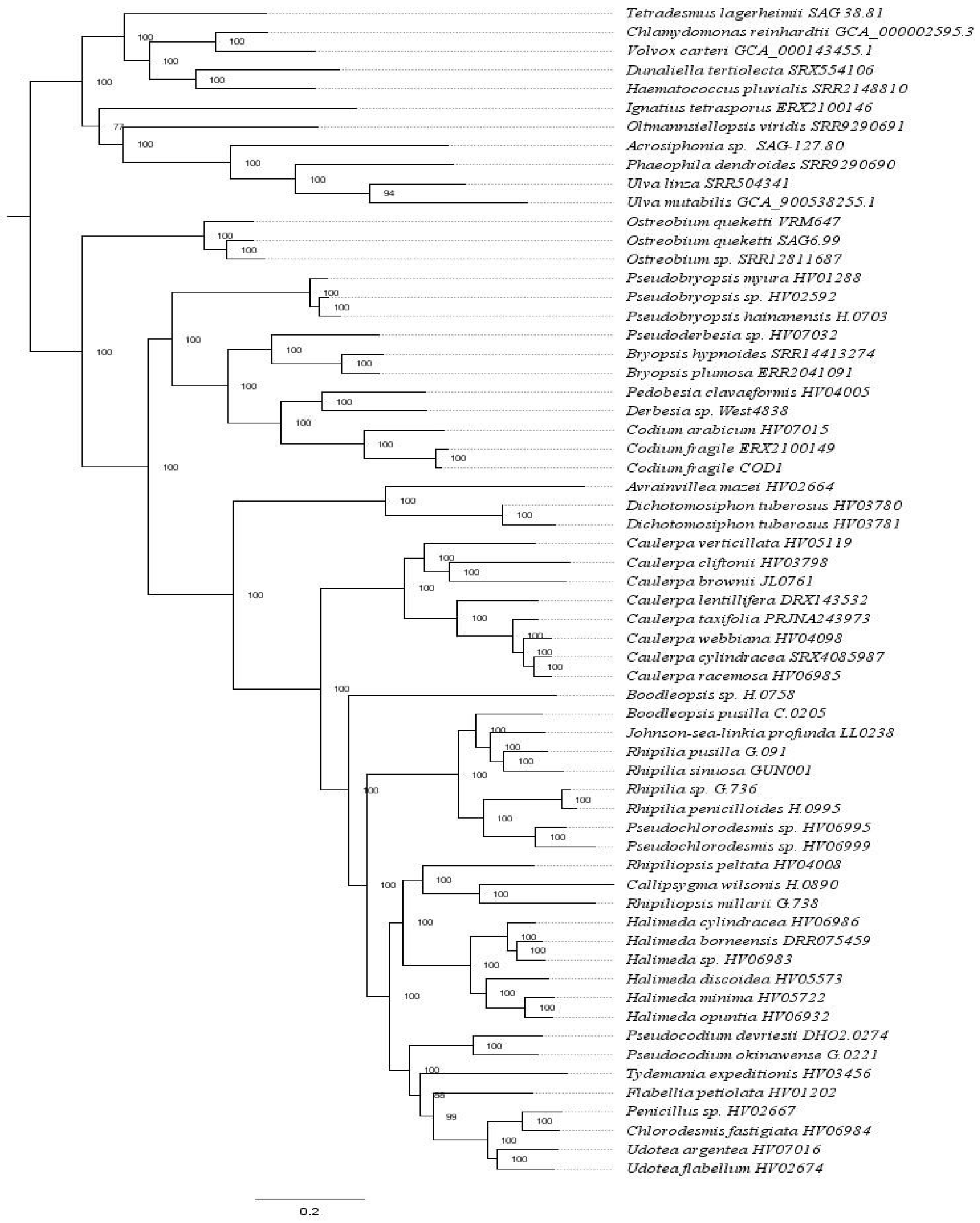


**Fig. S15** Maximum likelihood tree of the concatenated nucleotide alignment, using the GTR+F+I+R7 model, and with Chlorophyceae and other Ulvophyceae as outgroups.

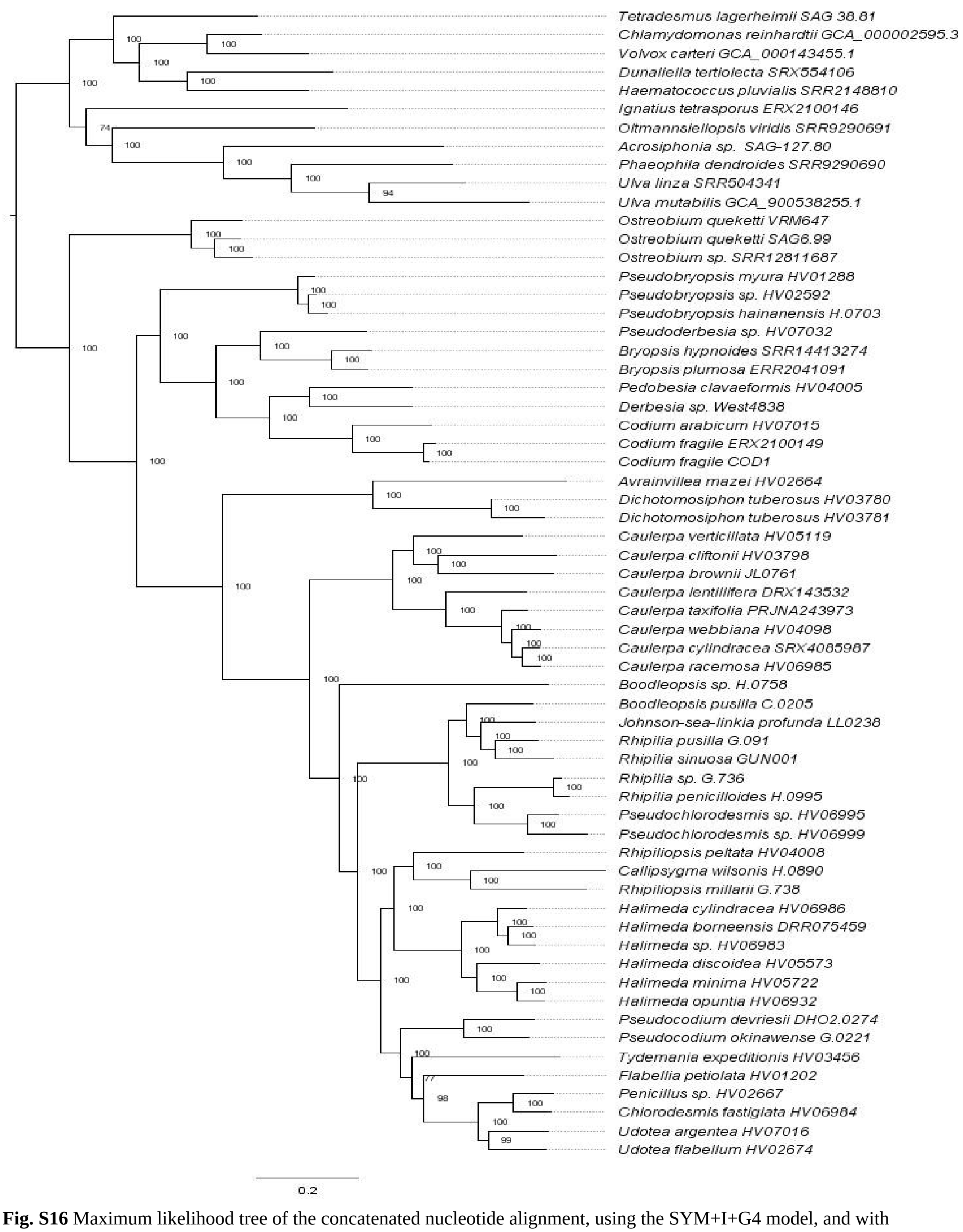


**Fig. S16** Maximum likelihood tree of the concatenated nucleotide alignment, using the SYM+I+G4 model, and with Chlorophyceae and other Ulvophyceae as outgroups.

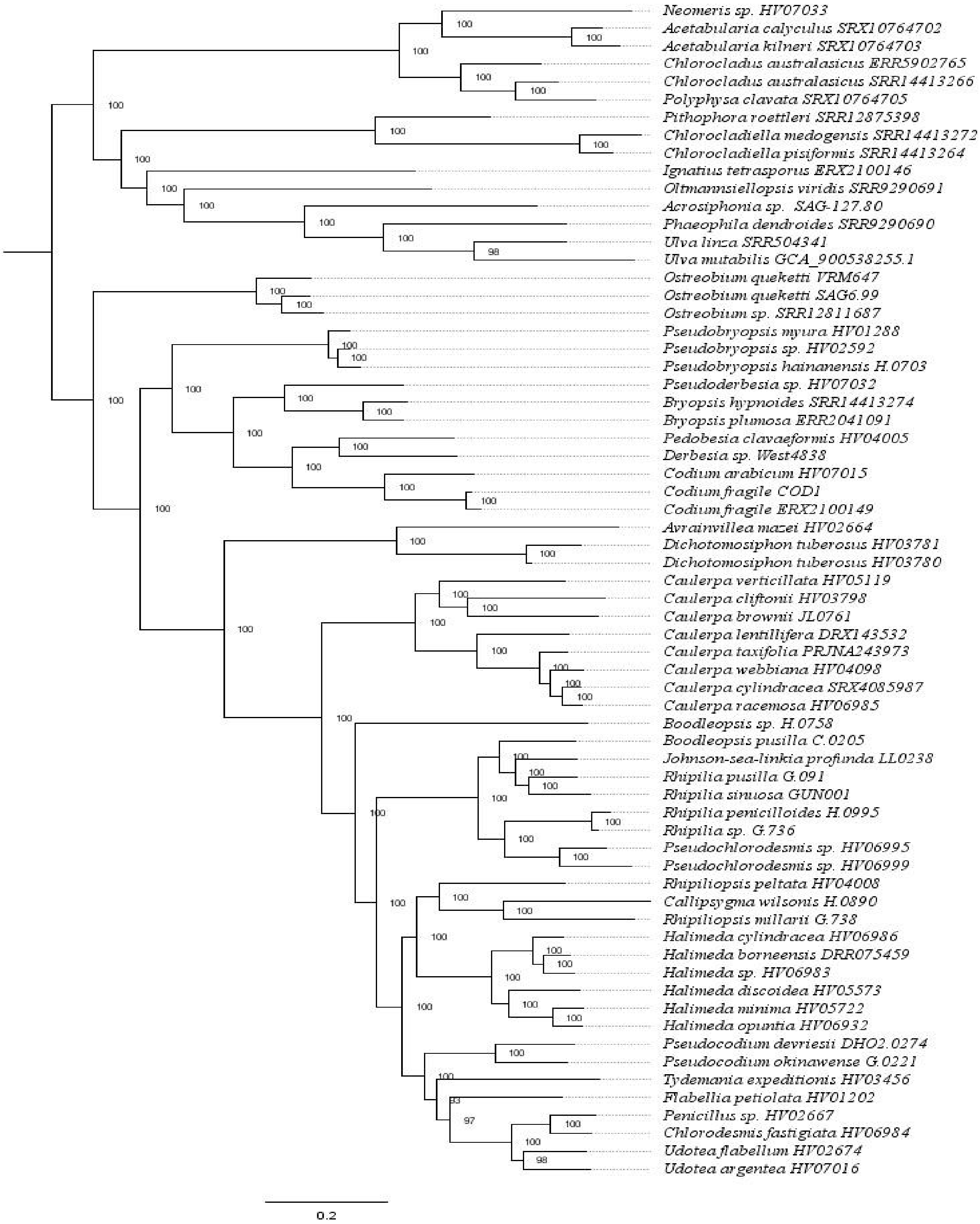


**Fig. S17** Maximum likelihood tree of the concatenated nucleotide alignment, using the GTR+F+I+R7 model, and with Dasycladales, Cladophorales and other Ulvophyceae as outgroups.

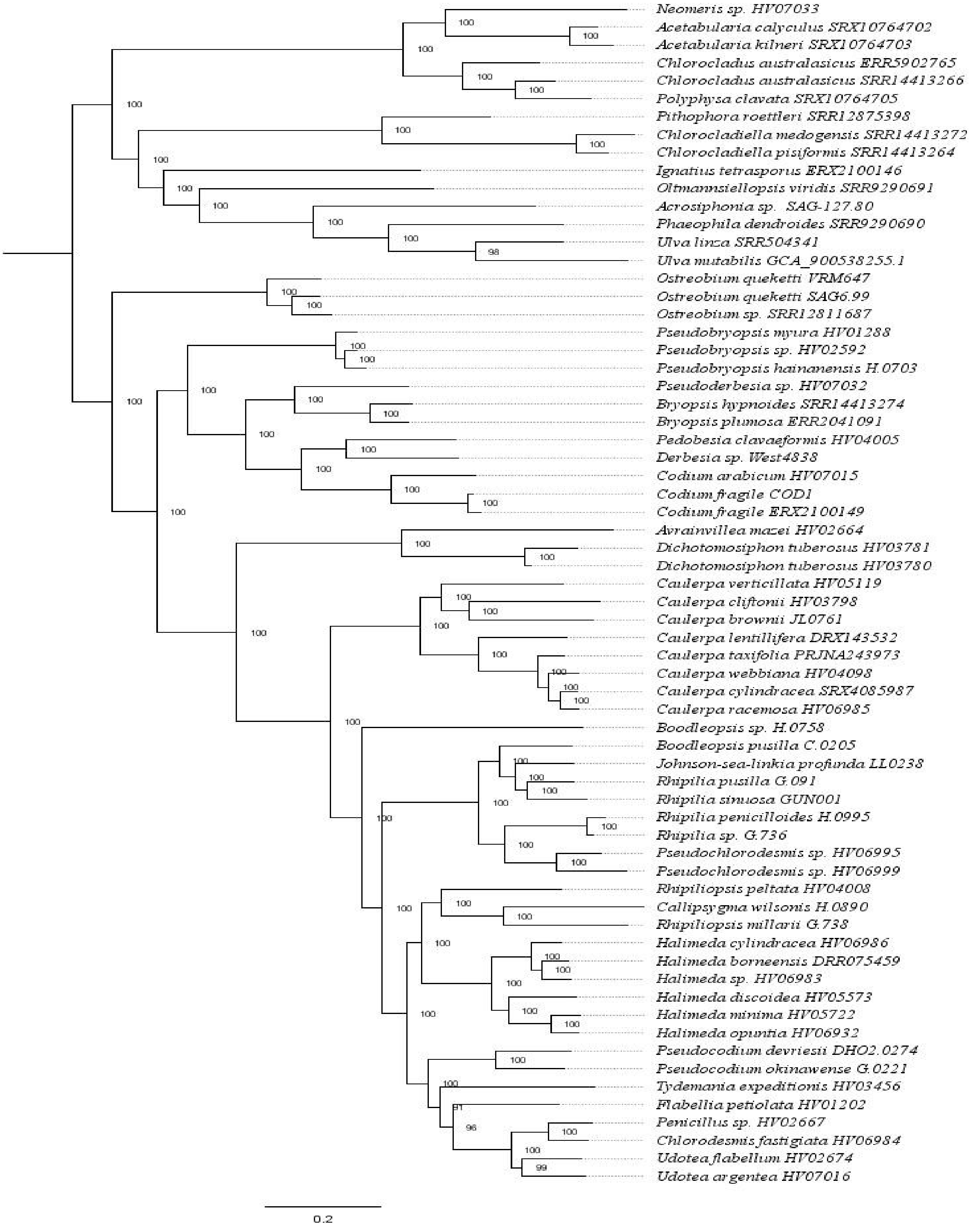


**Fig. S18** Maximum likelihood tree of the concatenated nucleotide alignment, using the GTR+F+I+R6 model, and with Dasycladales, Cladophorales and other Ulvophyceae as outgroups.

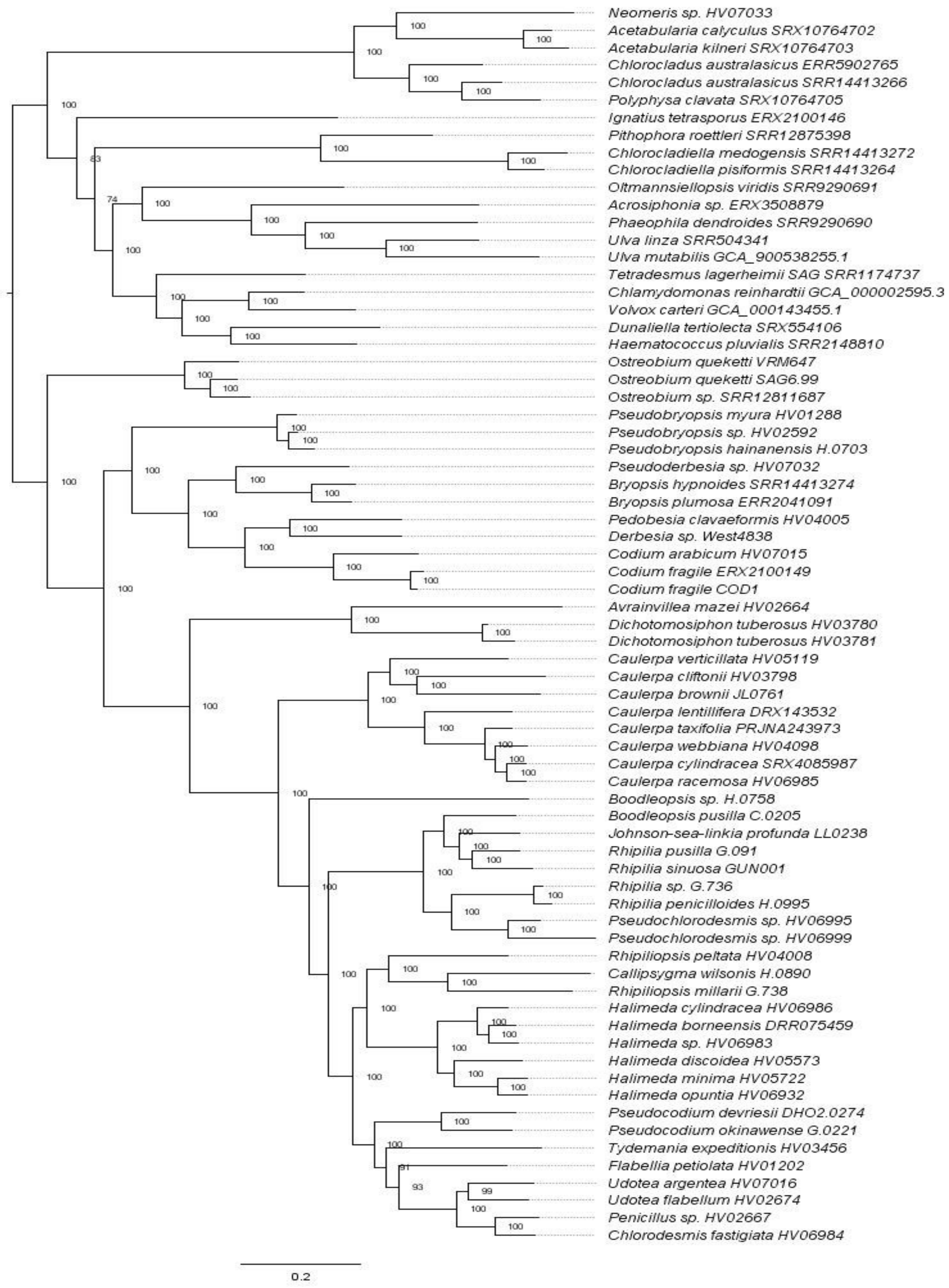


**Fig. S19** Maximum likelihood tree of the concatenated nucleotide alignment, using the GTR+F+I+R7 model, and with Dasycladales, Cladophorales, Chlorophyceae and other Ulvophyceae as outgroups.

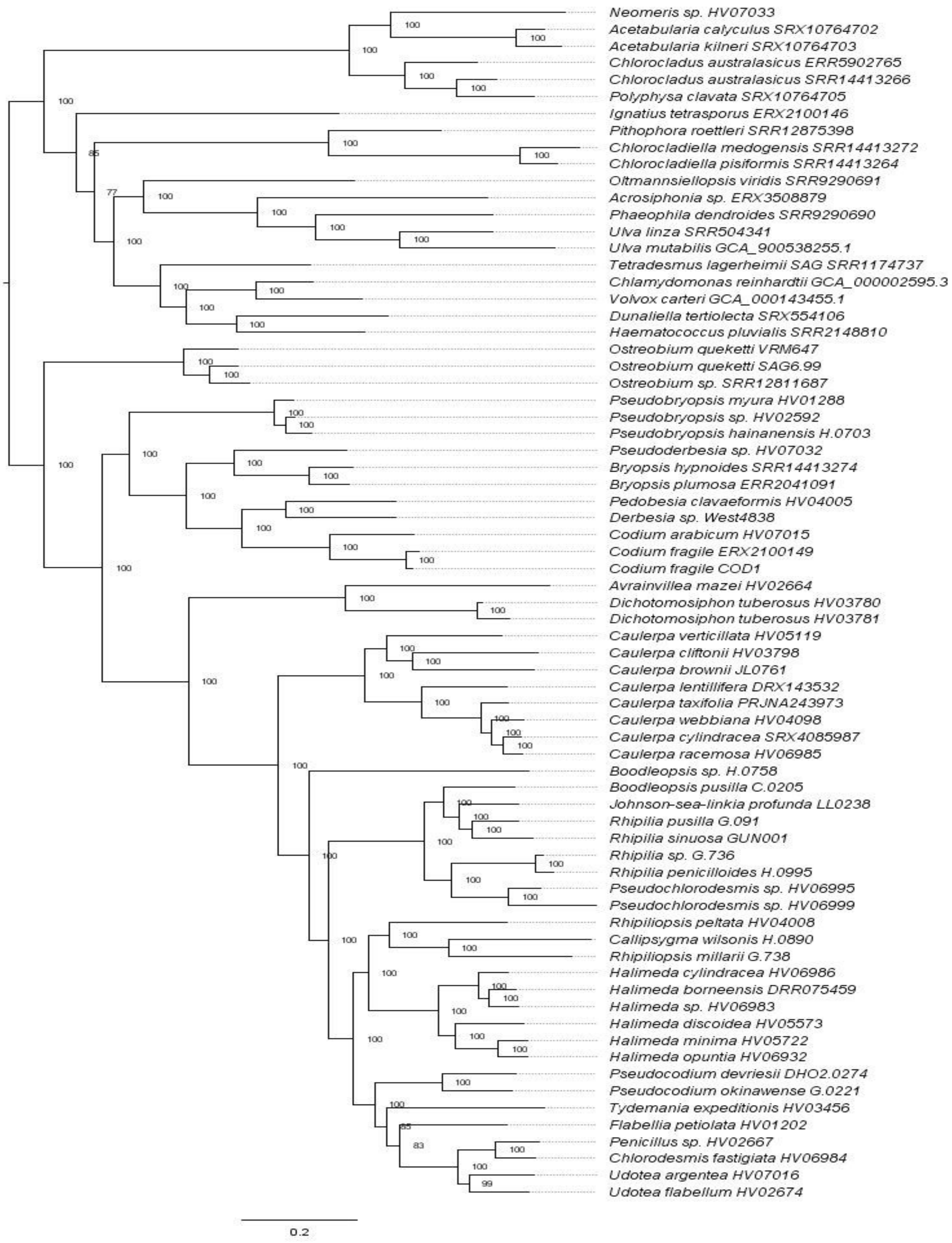


**Fig. S20** Maximum likelihood tree of the concatenated nucleotide alignment, using the model suggested by MFP, and with Dasycladales, Cladophorales, Chlorophyceae and other Ulvophyceae as outgroups.

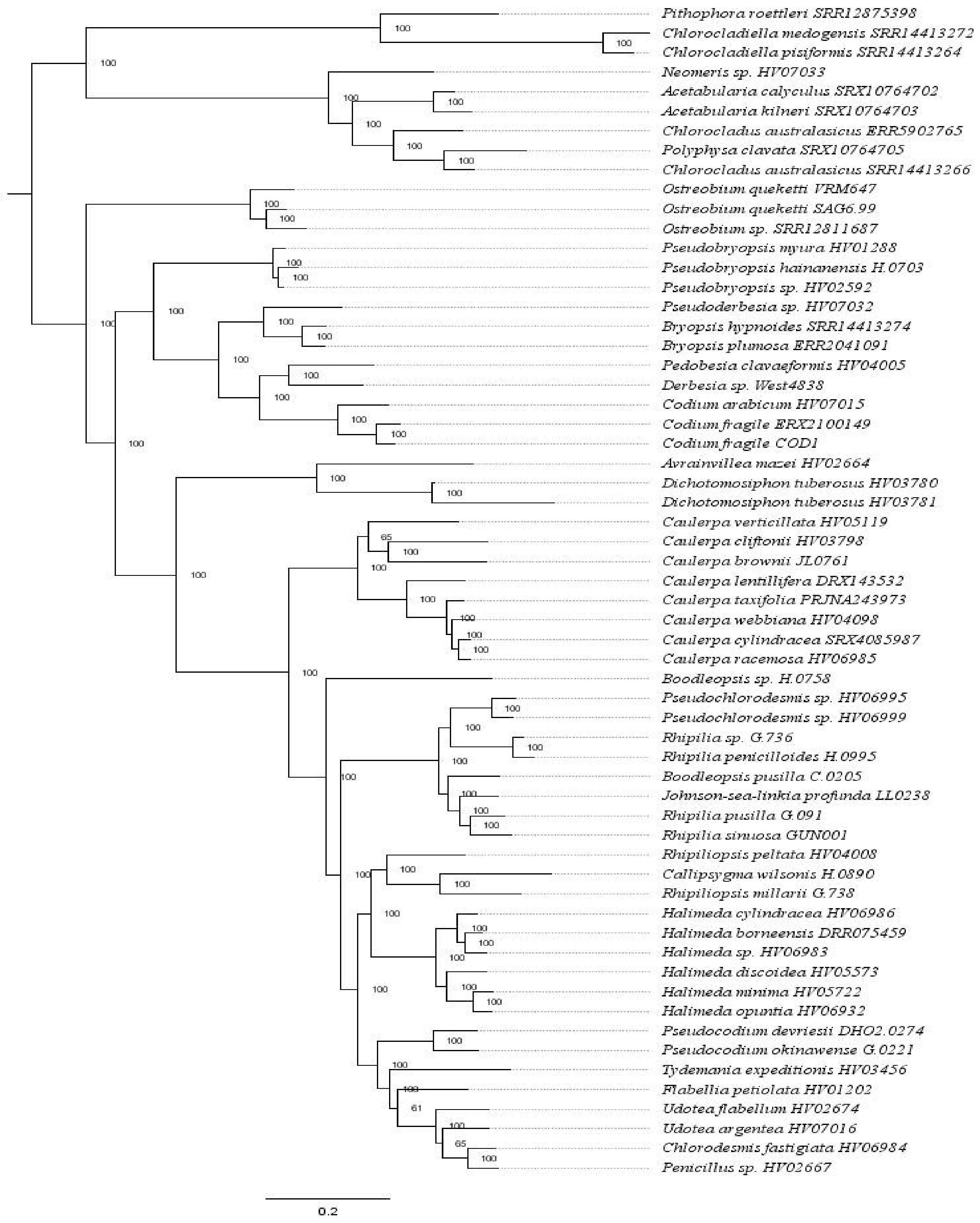


**Fig. S21** Maximum likelihood tree of the concatenated amino acid alignment, using the LG+F+I+G4 model, and with Dasycladales and Cladophorales as outgroups.

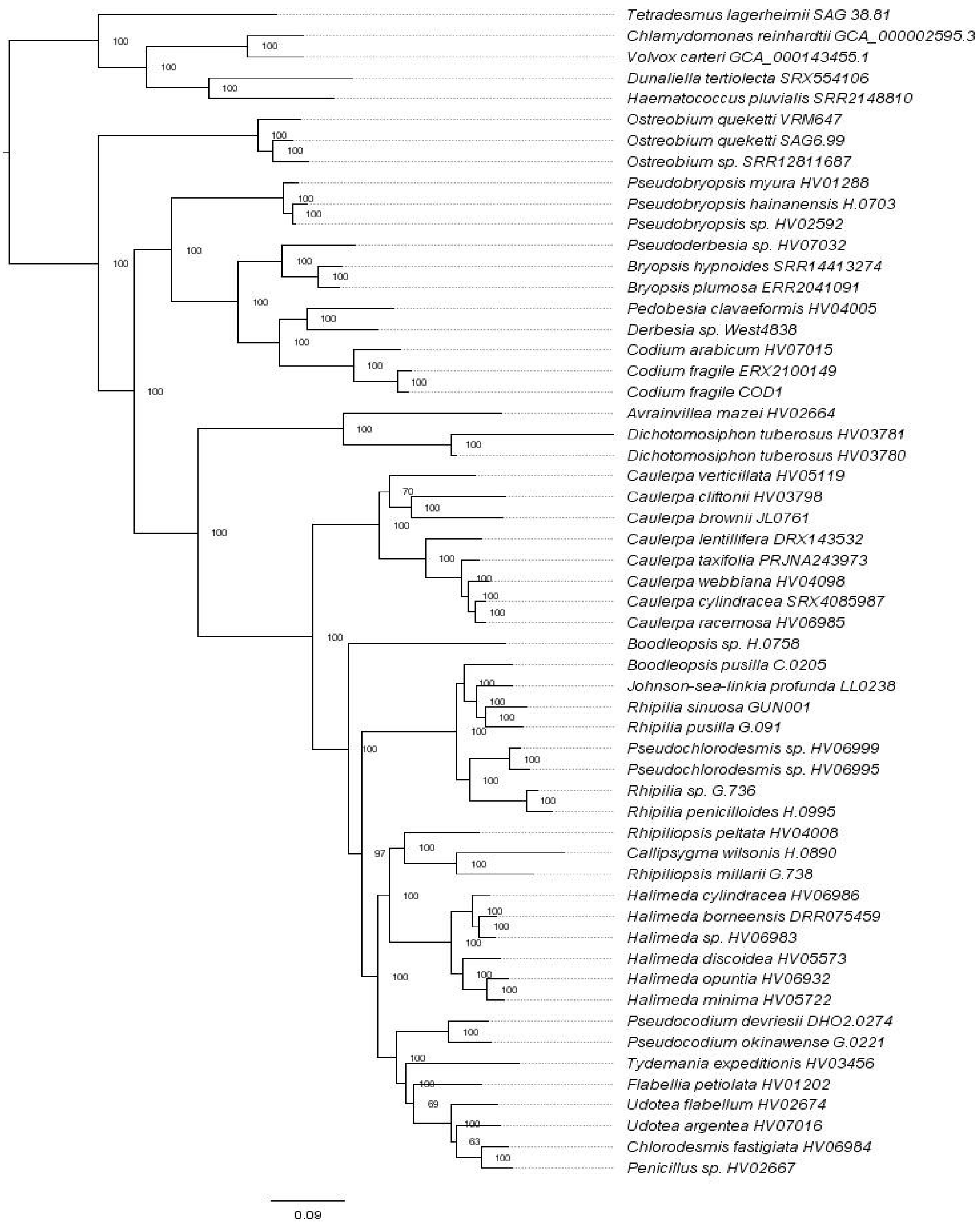


**Fig. S22** Maximum likelihood tree of the concatenated amino acid alignment, using the LG+I+R6 model, and with Chlorophyceae as the outgroup.

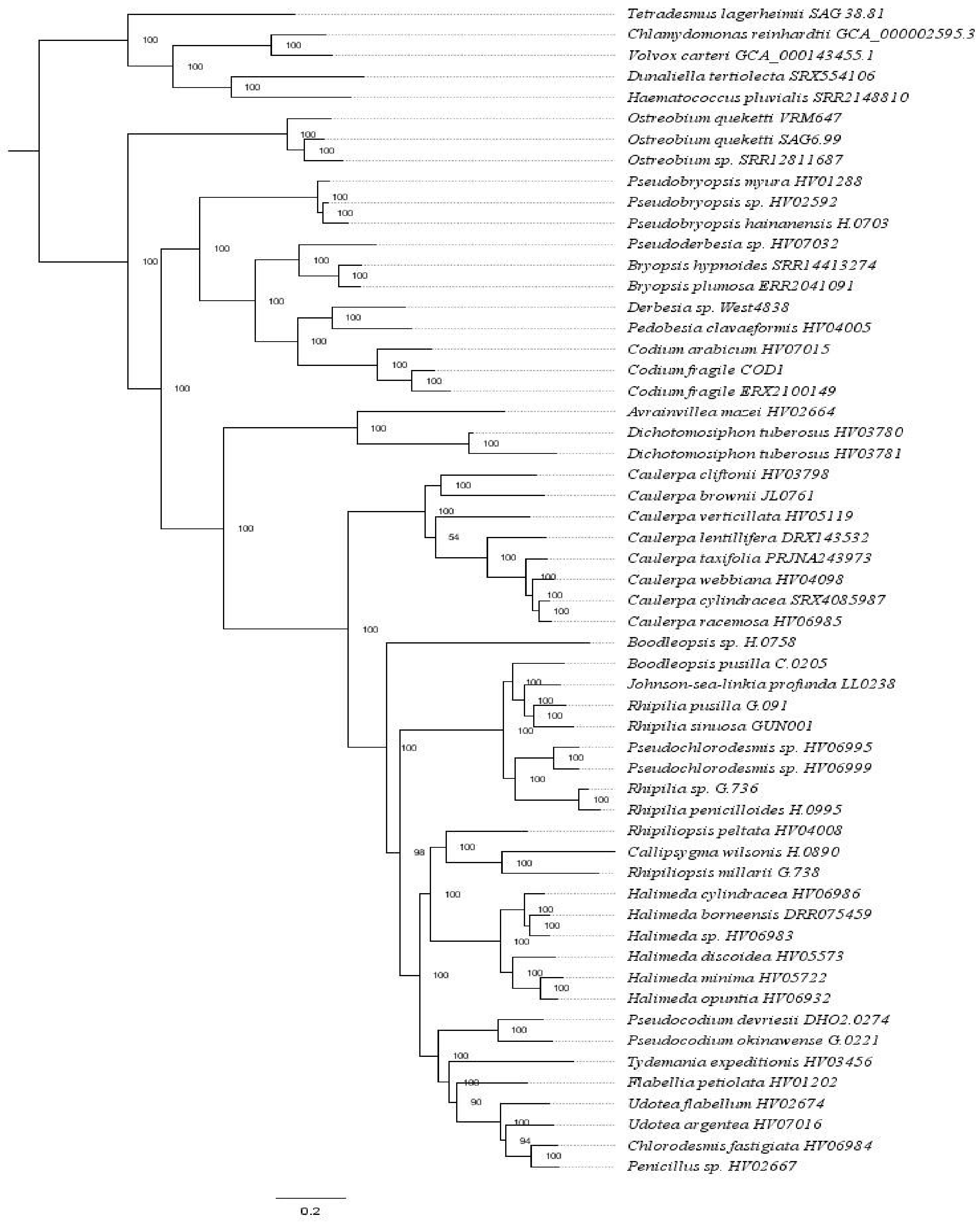


**Fig. S23** Maximum likelihood tree of the concatenated amino acid alignment, using the model suggested by MFP, and with Chlorophyceae as the outgroup.

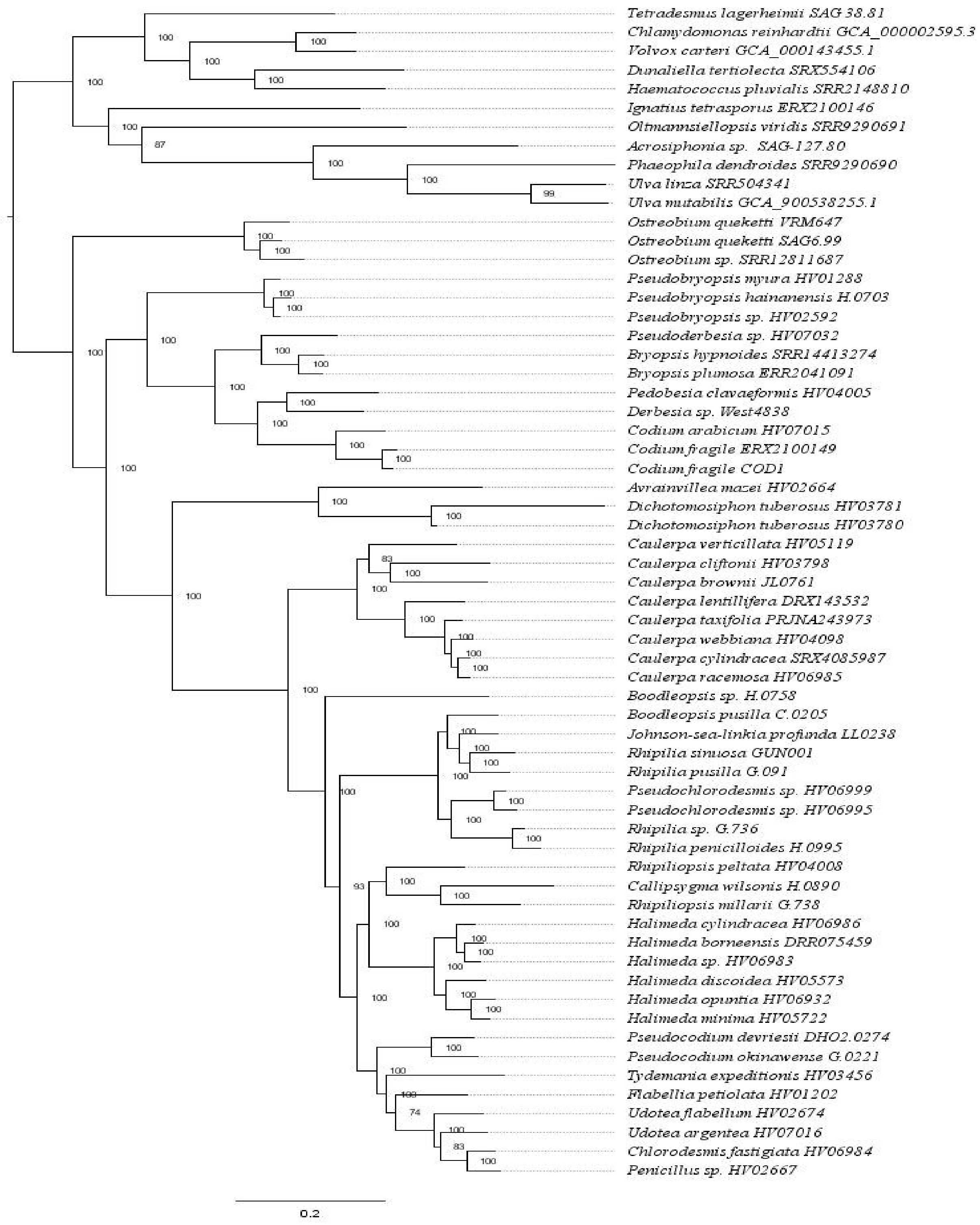


**Fig. S24** Maximum likelihood tree of the concatenated amino acid alignment, using the LG+F+I+G4 model, and with Chlorophyceae and other Ulvophyceae as outgroups.

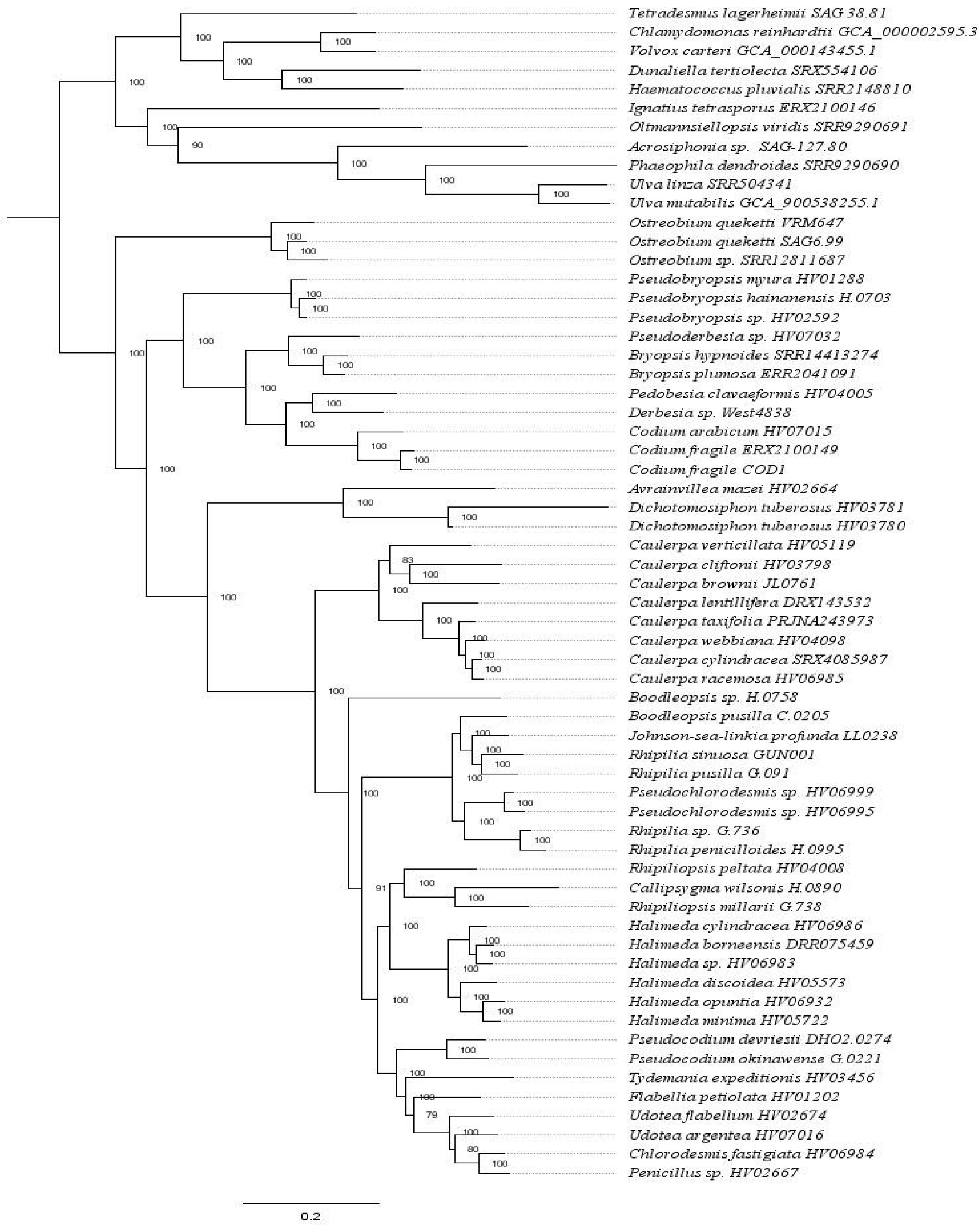


**Fig. S25** Maximum likelihood tree of the concatenated amino acid alignment, using the LG+I+G4 model, and with Chlorophyceae and other Ulvophyceae as outgroups.

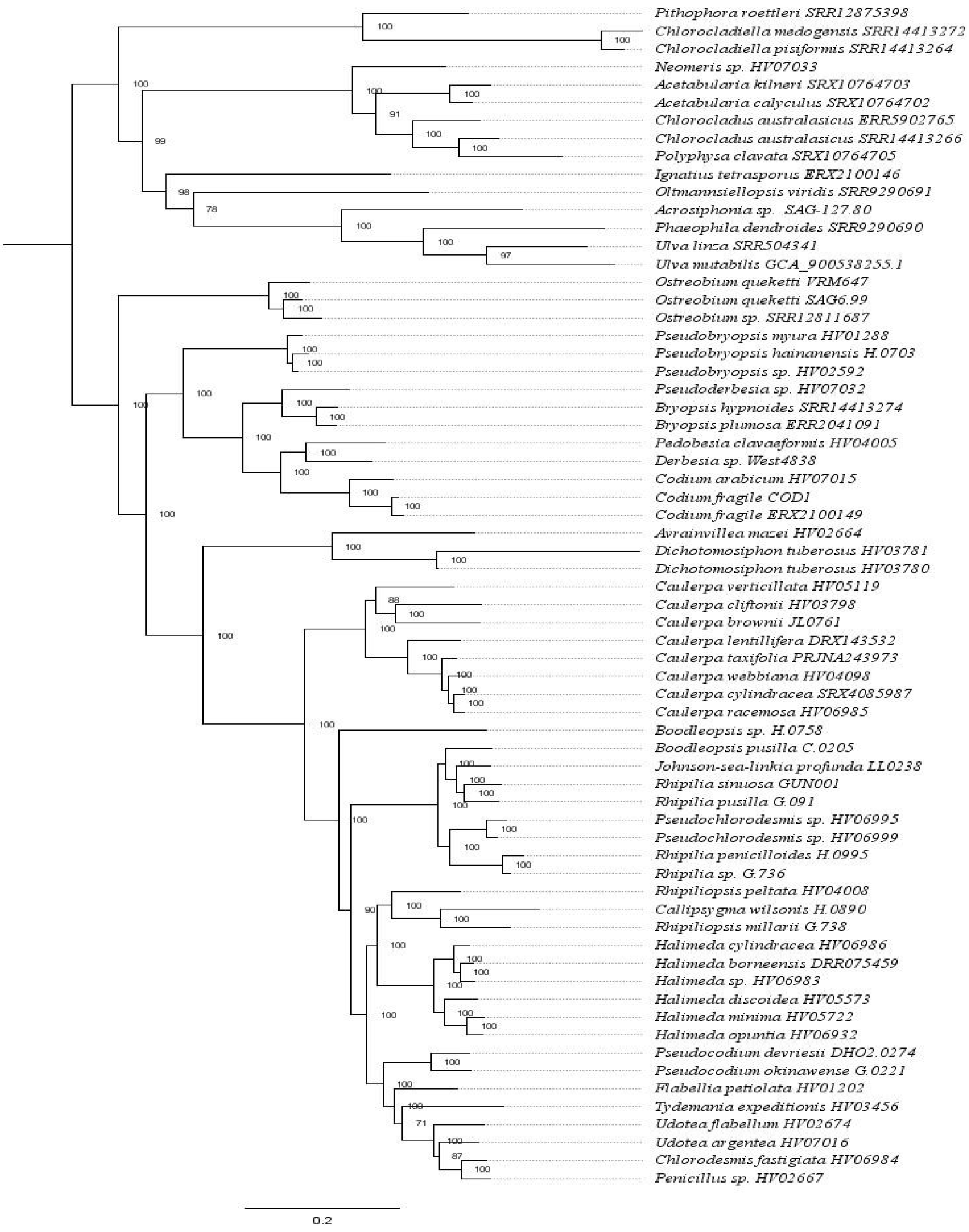


**Fig. S26** Maximum likelihood tree of the concatenated amino acid alignment, using the model suggested by MFP, and with Dasycladales, Cladophorales and other Ulvophyceae as outgroups.

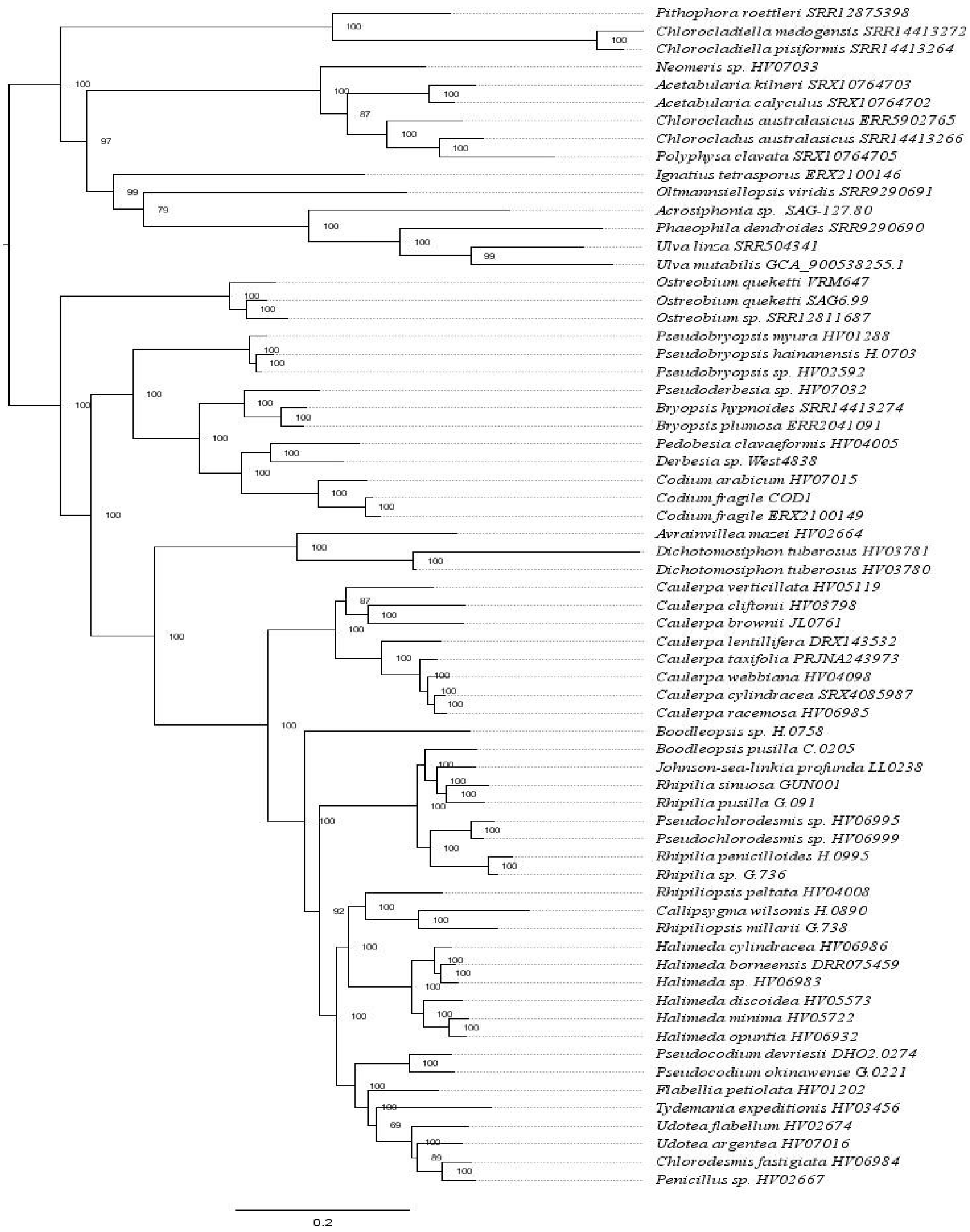


**Fig. S27** Maximum likelihood tree of the concatenated amino acid alignment, using the LG+F+I+R6 model, and with Dasycladales, Cladophorales and other Ulvophyceae as outgroups.

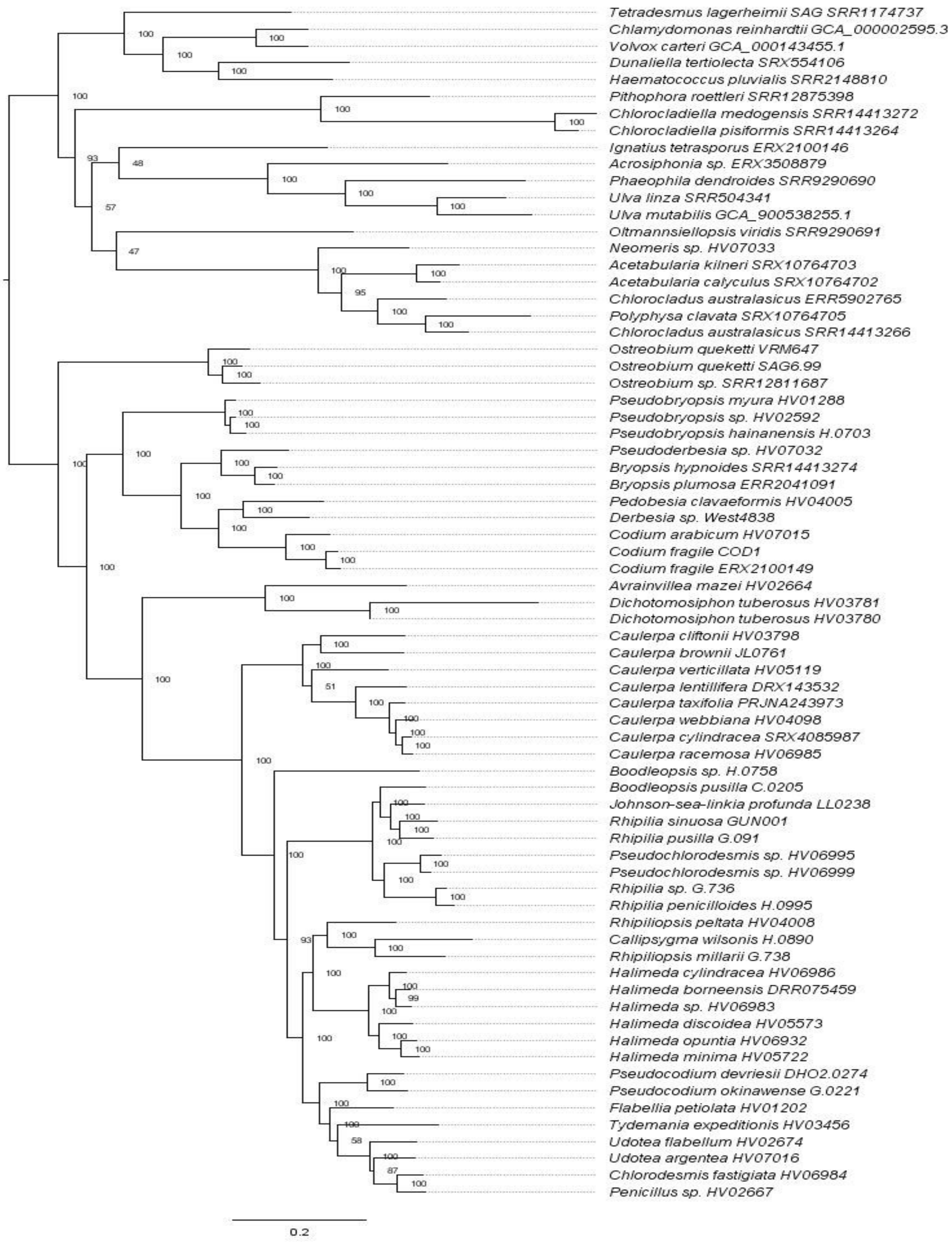


**Fig. S28** Maximum likelihood tree of the concatenated amino acid sequence alignment, using the LG+F+I+G4 model, and with Dasycladales, Cladophorales, Chlorophyceae and other Ulvophyceae as outgroups.

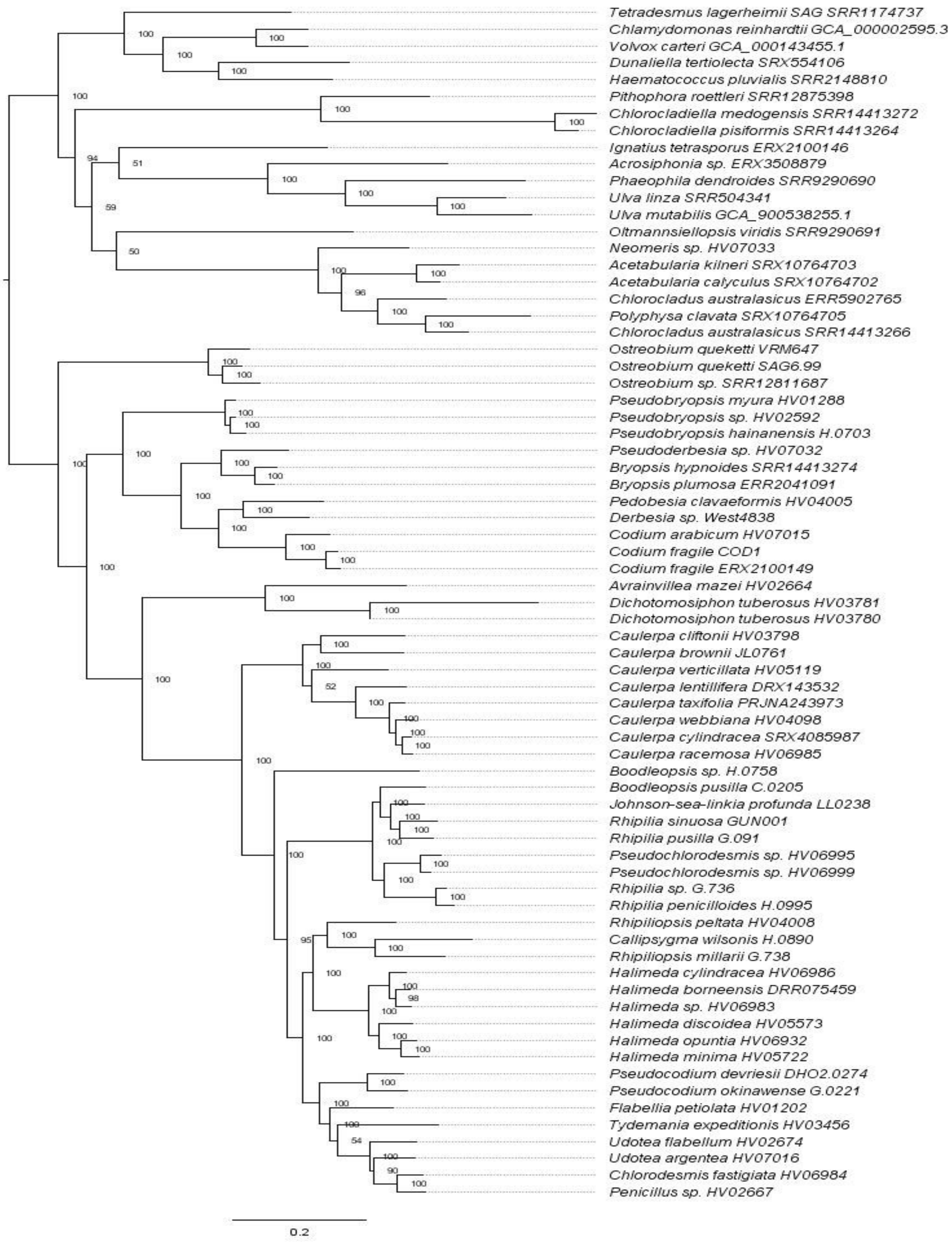


**Fig. S29** Maximum likelihood tree of the concatenated amino acid alignment, using the model suggested by MFP, and with Dasycladales, Cladophorales, Chlorophyceae and other Ulvophyceae as outgroups.

***rbcL* Tree (CDS sequences)**

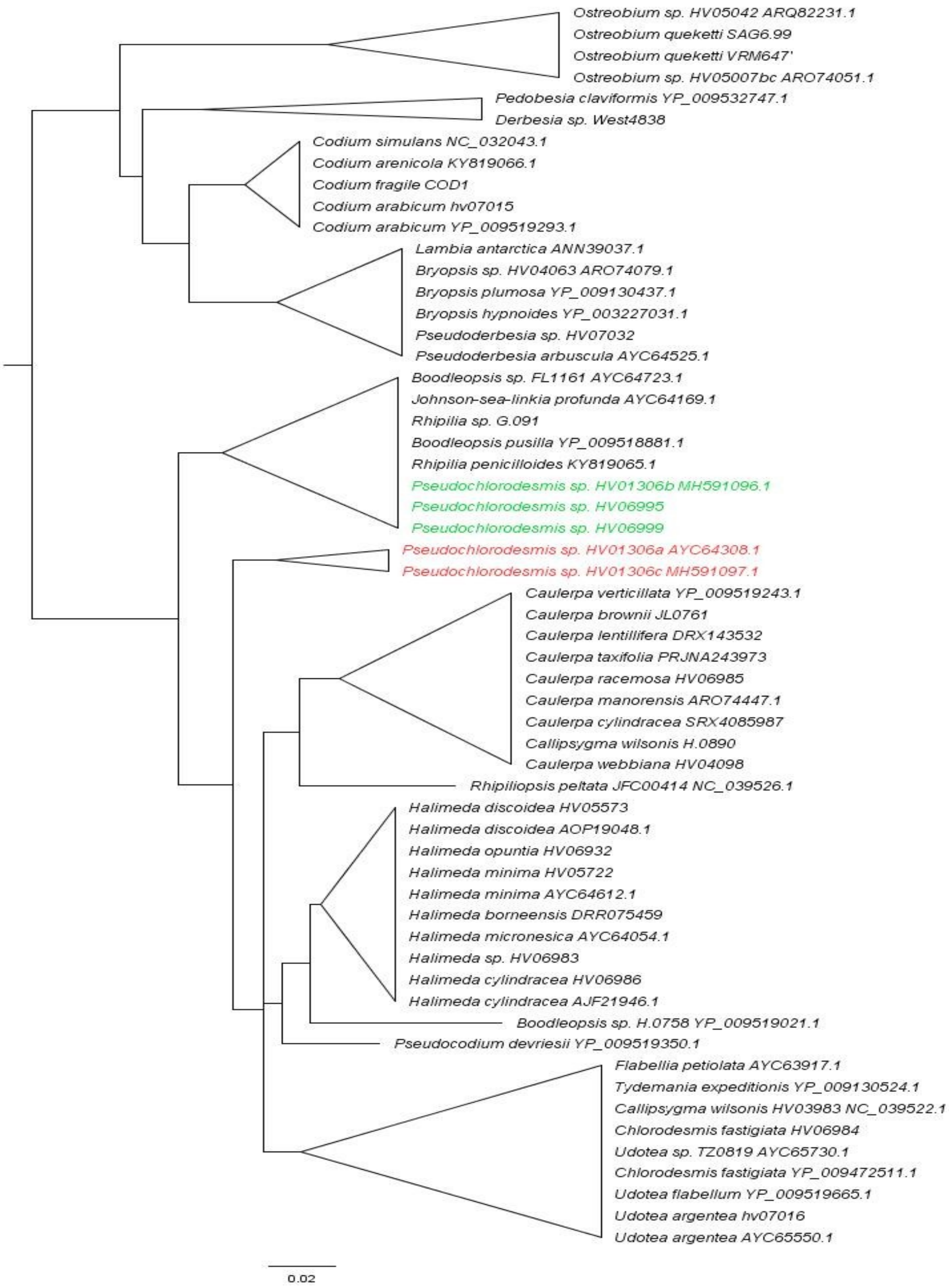


**Fig. S30** Phylogenetic tree inferred from nucleotide sequences of the chloroplast-localised *rbcL* gene, highlighting the placement of newly incorporated *Pseudochlorodesmis* taxa alongside those from Cremen et al. (2019).

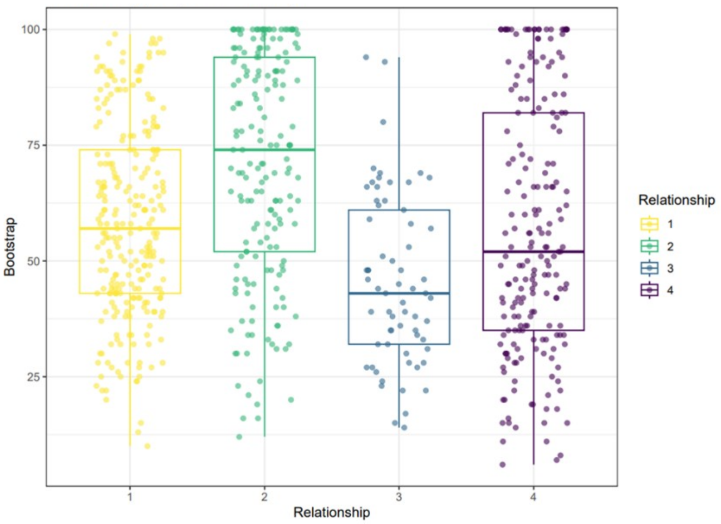


**Fig. S31** Jitter plot of bootstrap values showing the distribution of phylogenetic support for alternative relationships for Ostreobineae. The observed topologies include: (1) Ostreobineae as sister to Bryopsidineae; (2) sister to Bryopsidineae + Halimedineae; (3) nested within Bryopsidineae; and (4) placed elsewhere.

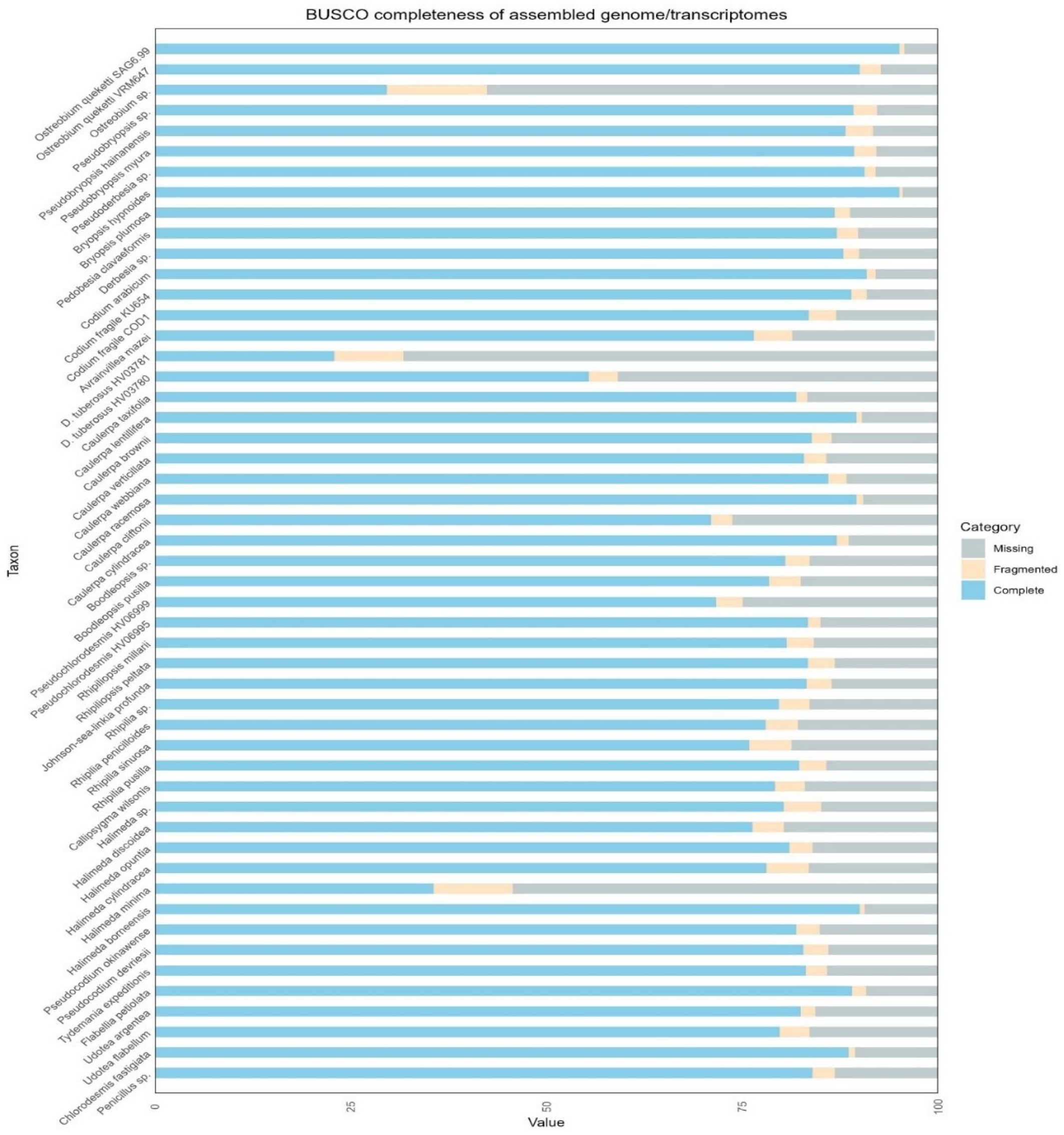


**Fig. S32** Distribution of BUSCO scores based on predicted coding sequences across all the Bryopsidalean taxa.

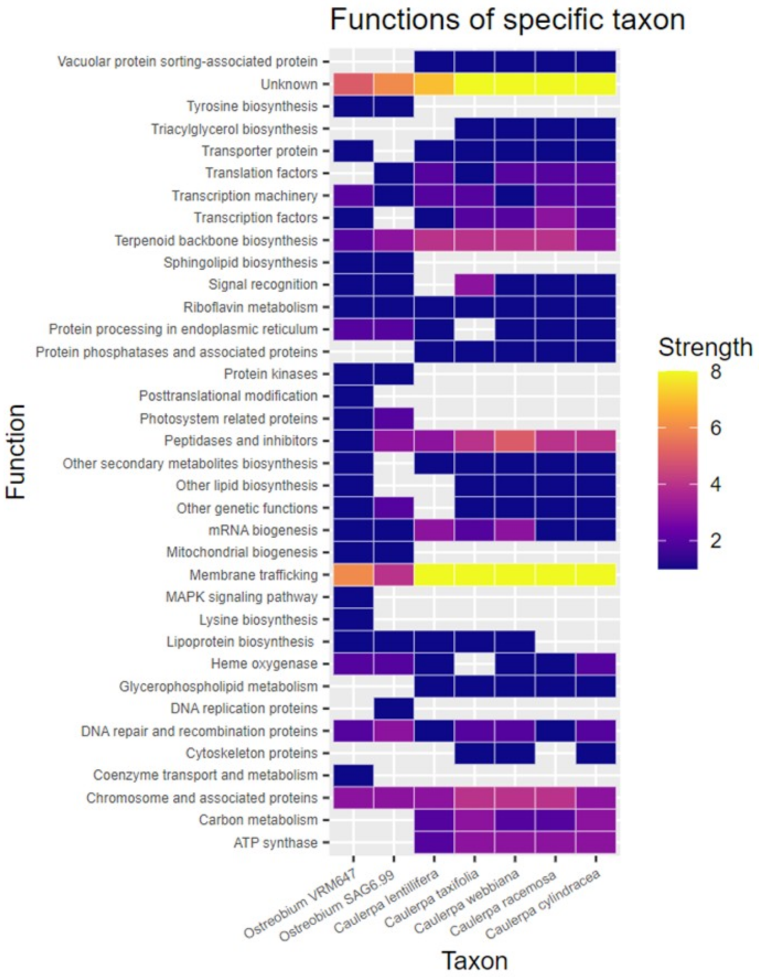


**Fig. S33** Heatmap showing functions of the retained outparalogs in two *Ostreobium* species and five *Caulerpa* species.

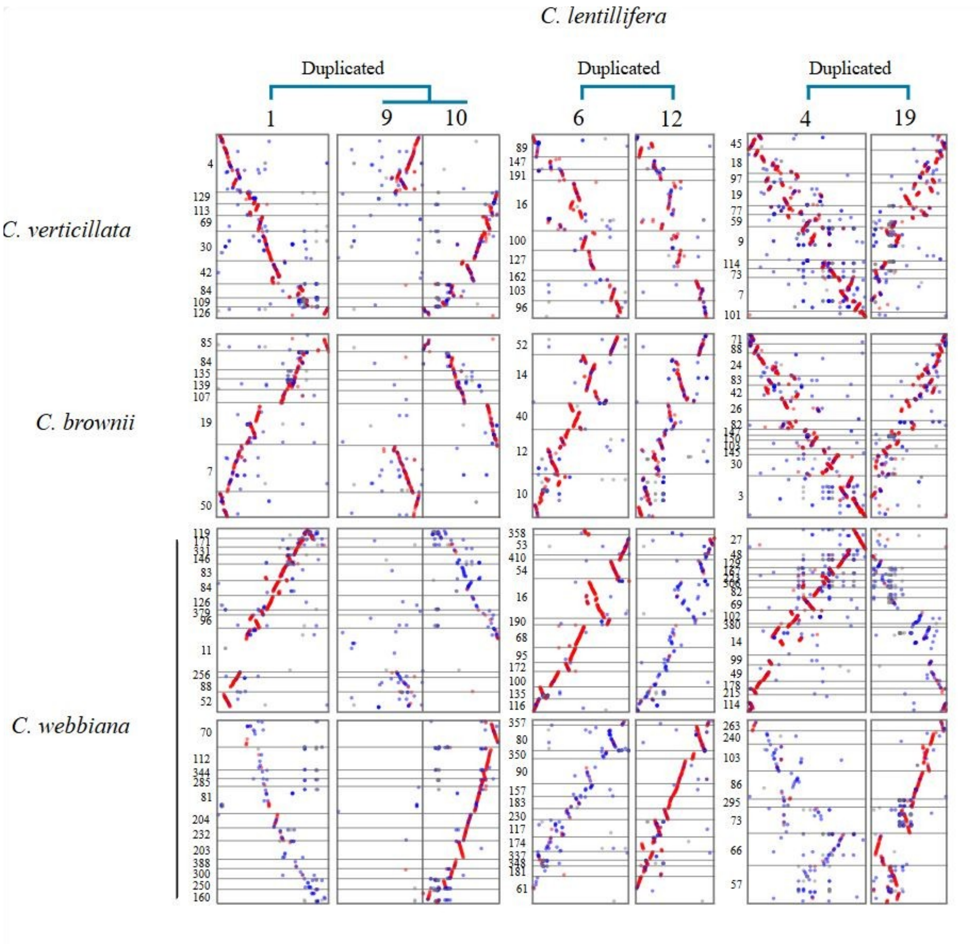


**Fig. S34.** The synteny blocks presenting the blast hits including contigs with their matches per chromosome in Caulerpaceae lineage.

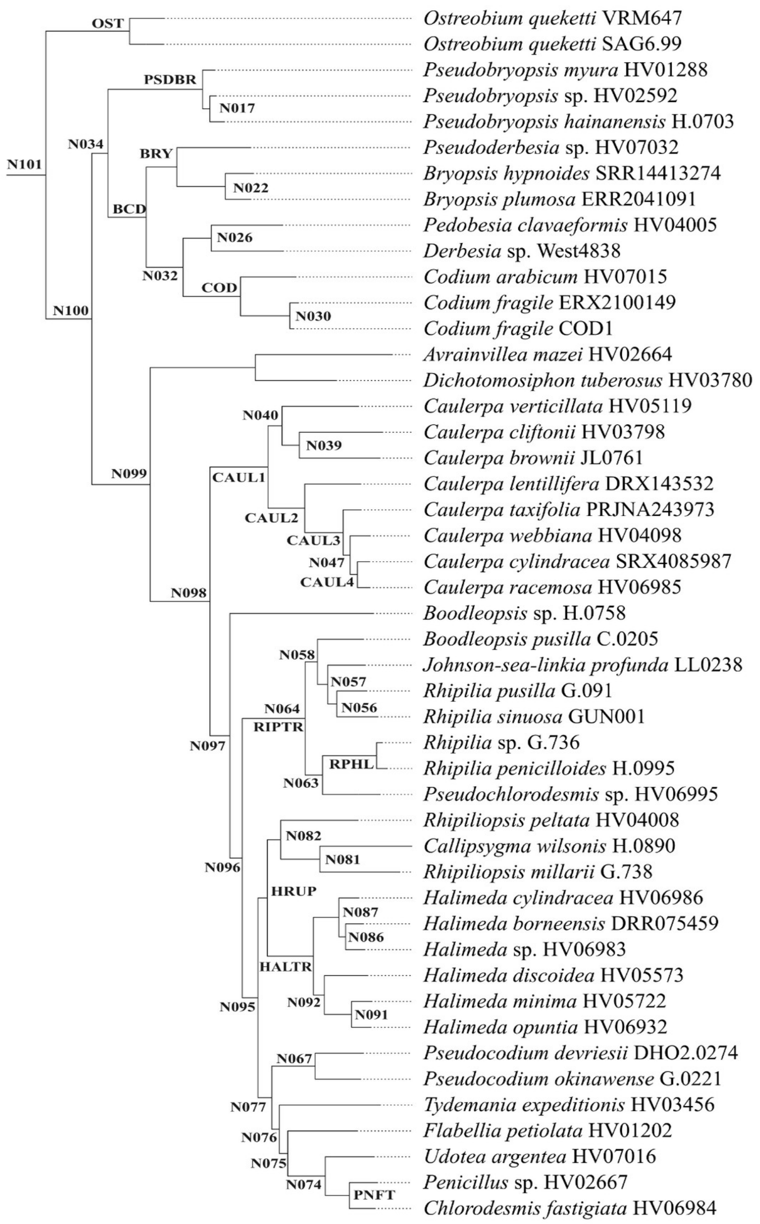


**Fig. S35.** Phylogeny showing the labeled nodes used for gene tree simulations, reconciliation analyses, and statistical testing.

**Table S1** Sampling details including higher-level classification, NCBI accession, origin (with new denoting newly sequenced here and public denoting previously published and publicly available) and sequence type (DNA or RNA library).

| SL | Taxon | Strain/Accession | Classification | Origin | Type |
|---|---|---|---|---|---|
| 1 | *Ostreobium* sp. | HV05007 | Ostreobiaceae | Public | RNA |
| 2 | *Ostreobium quekettii* Bornet & Flahault | VRM647 | | New | RNA |
| 3 | *Ostreobium quekettii* Bornet & Flahault | SAG6.99 | | New | RNA |
| 4 | *Pseudobryopsis* sp. | HV02592 | Pseudobryopsidaceae | New | DNA |
| 5 | *Pseudobryopsis myura* (J.Agardh) Berthold | HV01288 | | New | DNA |
| 6 | *Pseudobryopsis hainanensis* C.K.Tseng | H.0703 | | New | DNA |
| 7 | *Derbesia* sp. | West4838 | Derbesiaceae | New | RNA |
| 8 | *Pedobesia clavaeformis* (J.Agardh) MacRaild & Womersley | HV04005 | | New | DNA |
| 9 | *Codium arabicum* Kützing | HV07015 | Codiaceae | New | RNA |
| 10 | *Codium fragile* (Suringar) Hariot | KU654 | | Public | RNA |
| 11 | *Codium fragile* (Suringar) Hariot | COD1 | | New | RNA |
| 12 | *Bryopsis hypnoides* J.V.Lamouroux | SRR14413274 | Bryopsidaceae | Public | RNA |
| 13 | *Bryopsis plumosa* (Hudson) C.Agardh | ERR2041091 | | Public | RNA |
| 14 | *Pseudoderbesia* sp. | HV07032 | | New | RNA |
| 15 | *Dichotomosiphon tuberosus* (A.Braun ex Kützing) A.Ernst | HV03781 | Dichotomosiphonaceae | New | RNA |
| 16 | *Dichotomosiphon tuberosus* (A.Braun ex Kützing) A.Ernst | HV03780 | | New | DNA |
| 17 | *Avrainvillea mazei* G.Murray & Boodle | HV02664 | | New | DNA |
| 18 | *Caulerpa lentillifera* J.Agardh | DRX143532 | Caulerpaceae | Public | RNA |
| 19 | *Caulerpa taxifolia* (M.Vahl) C.Agardh | PRJNA243973 | | Public | RNA |
| 20 | *Caulerpa cylindracea* Sonder | SRX4085987 | | Public | RNA |
| 21 | *Caulerpa racemosa* (Forsskål) J.Agardh | HV06985 | | New | RNA |
| 22 | *Caulerpa verticillata J.Agardh* | HV05119 | | New | DNA |
| 23 | *Caulerpa brownii* (C.Agardh) Endlicher | JL0761 | | New | DNA |
| 24 | *Caulerpa cliftonii* Harvey | HV03798 | | New | DNA |
| 25 | *Caulerpa webbiana* Montagne | HV04098 | | New | DNA |
| 26 | *Halimeda* sp. | HV06983 | Halimedaceae | New | RNA |
| 27 | *Halimeda borneensis* W.R.Taylor | DRR075459 | | Public | RNA |
| 28 | *Halimeda discoidea* Decaisne | HV05573 | | New | RNA |
| 29 | *Halimeda opuntia* (Linnaeus) J.V.Lamouroux | HV06932 | | New | RNA |
| 30 | *Halimeda cylindracea* Decaisne | HV06986 | | New | RNA |
| 31 | *Halimeda minima* (W.R.Taylor) Hillis-Colinvaux | HV05722 | | New | RNA |
| 32 | *Rhipilia penicilloides* A.D.R.N'Yeurt & D.W.Keats | H.0995 | | New | DNA |
| 33 | *Rhipilia sinuosa* Gilbert | GUN001 | | New | DNA |
| 34 | *Rhipilia* sp. | G.736 | | New | DNA |
| 35 | *Rhipilia pusilla* (Womersley) Ducker | G.091 | | New | DNA |
| 36 | *Callipsygma wilsonis* J.Agardh | H.0890 | | New | DNA |
| 37 | *Johnson-sea-linkia profunda* Eiseman & S.A.Earle | LL0238 | | New | DNA |
| 38 | *Rhipiliopsis peltata* (J.Agardh) A.Gepp & E.Gepp | HV04008 | | New | DNA |
| 39 | *Rhipiliopsis millarii* Kraft | G.738 | | New | DNA |
| 40 | *Pseudochlorodesmis* sp. | HV06995 | | New | RNA |
| 41 | *Pseudochlorodesmis* sp. | HV06999 | | New | RNA |
| 42 | *Boodleopsis* sp. | H.0758 | | New | DNA |
| 43 | *Boodleopsis pusilla* (Collins) W.R.Taylor, A.B.Joly & Bernatowicz | C.0205 | | New | DNA |
| 44 | *Pseudocodium devriesii* Weber Bosse | DHO2.0274 | | New | DNA |
| 45 | *Pseudocodium okinawense* E.J.Faye, M.Uchimura & S.Smimada | G.0221 | | New | DNA |
| 46 | *Udotea argentea* Zanardini | HV07016 | | New | RNA |
| 47 | *Udotea flabellum* (J.Ellis & Solander) M.Howe | HV02674 | | New | RNA |

| | | | | | |
|---|---|---|---|---|---|
| 48 | *Penicillus* sp. | HV02667 | | New | DNA |
| 49 | *Chlorodesmis fastigiata* (C.Agardh) S.C.Ducker | HV06984 | | New | RNA |
| 50 | *Flabellia petiolata* (Turra) Nizamuddin | HV01202 | | New | DNA |
| 51 | *Tydemania expeditionis* Weber Bosse | HV03456 | | New | DNA |
| 52 | *Acetabularia calyculus* J.V.Lamouroux | SRX10764702 | Dasycladales | Public | RNA |
| 53 | *Acetabularia kilneri* J.Agardh | SRX10764703 | | Public | RNA |
| 54 | *Polyphysa clavata* (Yamada) Schnetter & Bula-Mayer | SRX10764705 | | Public | RNA |
| 55 | *Chloroclados australasicus* Sonder | SRR14413266 | | Public | RNA |
| 56 | *Neomeris* sp. | HV07033 | | New | RNA |
| 57 | *Chlorocladiella pisiformis* Huan Zhu, Guoxiang Liu & Zhengyu Hu | SRR14413264 | Cladophorales | Public | RNA |
| 58 | *Chlorocladiella medogensis* Huan Zhu, Guoxiang Liu & Zhengyu Hu | SRR14413272 | | Public | RNA |
| 59 | *Pithophora roettleri* (Roth) Wittrock | SRR12875398 | | Public | RNA |
| 60 | *Tetradesmus lagerheimii* M.J.Wynne & Guiry | SRR9290691 | Chlorophyceae | Public | RNA |
| 61 | *Chlamydomonas reinhardtii* P.A.Dangeard | GCA_000002595.3 | | Public | DNA |
| 62 | *Dunaliella tertiolecta* Butcher | SRX554106 | | Public | RNA |
| 63 | *Volvox carteri* F.Stein | GCA_000143455.1 | | Public | DNA |
| 64 | *Haematococcus pluvialis* Flotow | SRR2148810 | | Public | RNA |
| 65 | *Oltmannsiellopsis viridis* (P.E.Hargraves & R.L.Steele) Chihara & I.Inouye | SRR9290691 | Oltmansiellopsidales; Ignatiales; Ulotrichales; Ulvales | Public | RNA |
| 66 | *Ignatius tetrasporus* H.C.Bold & F.J.MacEntee | ERX2100146 | | Public | RNA |
| 67 | *Acrosiphonia* sp. | ERX3508879 | | Public | RNA |
| 68 | *Ulva mutabilis* Föyn | GCA_900538255.1 | | Public | DNA |
| 69 | *Phaeophila dendroides* (P.Crouan & H.Crouan) Batters | SRR9290690 | | Public | RNA |
| 70 | *Ulva linza* Linnaeus | SRR504341 | | Public | RNA |

**Table S2** List of taxa with the number of counted paralogs and their individual sequence representation across the selected 708 OGs. Taxa with low OG coverage are highlighted in red and were excluded from the paralog counting due to insufficient representation.

| SL | Name | No. of out-paralog | OGs coverage |
|---|---|---|---|
| 1 | *Ostreobium quekettii* SAG6.99 | 41 | 706 |
| 2 | *Ostreobium quekettii* VRM647 | 43 | 689 |
| 3 | *Ostreobium* sp. | -- | 325 |
| 4 | *Pseudobryopsis hainanensis* | 1 | 452 |
| 5 | *Pseudobryopsis myura* HV01288 | 1 | 658 |
| 6 | *Pseudobryopsis* sp. HV02592 | 1 | 666 |
| 7 | *Derbesia* sp. | 0 | 693 |
| 8 | *Pedobesia clavaeformis* | 1 | 681 |
| 9 | *Codium fragile* COD1 | 1 | 535 |
| 10 | *C. fragile* KU654 | 0 | 340 |
| 11 | *Codium arabicum* | 4 | 703 |
| 12 | *Bryopsis hypnoides* | 3 | 706 |
| 13 | *Bryopsis plumosa* | 2 | 700 |
| 14 | *Pseudoderbesia* sp. | 3 | 699 |
| 15 | *Dichotomosiphon tuberosus* HV03781 | -- | 117 |
| 16 | *Dichotomosiphon tuberosus* HV03780 | 0 | 549 |
| 17 | *Avrainvillea mazei* | 1 | 571 |
| 18 | *Caulerpa lentillifera* | 47 | 701 |
| 19 | *Caulerpa verticillata* | 1 | 687 |
| 20 | *Caulerpa taxifolia* | 60 | 674 |
| 21 | *Caulerpa cylindracea* | 58 | 699 |
| 22 | *Caulerpa racemosa* | 57 | 703 |
| 23 | *Caulerpa brownii* | 0 | 674 |
| 24 | *Caulerpa cliftonii* | 0 | 653 |
| 25 | *Caulerpa webbiana* | 63 | 700 |
| 26 | *Halimeda opuntia* | 4 | 658 |
| 27 | *Halimeda discoidea* | 3 | 675 |
| 28 | *Halimeda minima* | 2 | 418 |
| 29 | *Halimeda cylindracea* | 4 | 602 |
| 30 | *Halimeda* sp. | 3 | 588 |
| 31 | *Halimeda borneensis* | 5 | 697 |
| 32 | *Pseudochlorodesmis* sp. HV06995 | 2 | 621 |
| 33 | *Pseudochlorodesmis* sp. HV06999 | -- | 282 |
| 34 | *Udotea argentea* | 1 | 685 |
| 35 | *Udotea flabellum* | -- | 222 |
| 36 | *Penicillus* sp. | 2 | 668 |
| 37 | *Chlorodesmis fastigiata* | 2 | 701 |
| 38 | *Flabellia petiolata* | 1 | 688 |
| 39 | *Tydemania expeditionis* | 1 | 679 |
| 40 | *Rhipiliopsis peltata* | 3 | 678 |
| 41 | *Rhipiliopsis millarii* | 1 | 640 |
| 42 | *Johnson-sea-linkia profunda* | 1 | 665 |
| 43 | *Callipsygma wilsonis* | 1 | 678 |
| 44 | *Boodleopsis* sp. | 0 | 484 |
| 45 | *Boodleopsis pusilla* | 2 | 639 |
| 46 | *Rhipilia penicilloides* | 2 | 429 |
| 47 | *Rhipilia sinuosa* | 1 | 639 |
| 48 | *Rhipilia* sp. | 2 | 664 |
| 49 | *Rhipilia pusilla* | 3 | 667 |

| | | | |
|---|---|---|---|
| 50 | *Pseudocodium devriesii* | 1 | 676 |
| 51 | *Pseudocodium okinawense* | 0 | 654 |

**Table S3** Functional annotations of the 708 orthogroups used in this study, listing their BUSCO IDs and KEGG term inferred with BLASTKOALA.

| BUSCO_ID | KO_term | BUSCO_ID | KO_term | BUSCO_ID | KO_term |
|---|---|---|---|---|---|
| 15at3041 | K02925 | 2941at3041 | K03037 | 6072at3041 | K02734 |
| 42at3041 | K03265 | 2945at3041 | K15104 | 6074at3041 | K23095 |
| 45at3041 | K15104 | 2954at3041 | K02941 | 6097at3041 | K02148 |
| 91at3041 | K11594 | 2969at3041 | K13789 | 6111at3041 | K12737 |
| 155at3041 | No_hit | 2971at3041 | K01899 | 6144at3041 | K02868 |
| 171at3041 | K26752 | 2976at3041 | K12825 | 6145at3041 | K02134 |
| 187_2at3041 | K10807 | 2983at3041 | K14498 | 6146at3041 | K12272 |
| 207at3041 | K02150 | 2984at3041 | K12657 | 6156at3041 | K03648 |
| 207_1at3041 | K17267 | 2990at3041 | K01609 | 6192at3041 | K12873 |
| 211at3041 | K12614 | 3005at3041 | K03038 | 6198at3041 | K10580 |
| 228at3041 | K01714 | 3009at3041 | K08245 | 6207at3041 | K02150 |
| 230at3041 | K12581 | 3012at3041 | K09613 | 6225at3041 | K02149 |
| 231at3041 | K01662 | 3019at3041 | K06063 | 6290at3041 | K12849 |
| 239_1at3041 | K14416 | 3029at3041 | No_hit | 6330at3041 | K12188 |
| 239_2at3041 | K14416 | 3050at3041 | K14399 | 6334at3041 | K20280 |
| 251at3041 | K14293 | 3062at3041 | K14842 | 6345at3041 | K02303 |
| 290_1at3041 | K12849 | 3080at3041 | K13506 | 6347at3041 | No_hit |
| 290at3041 | K07904 | 3112at3041 | K03553 | 6348at3041 | K02964 |
| 294_1at3041 | K03094 | 3120at3041 | K05928 | 6355at3041 | No_hit |
| 294at3041 | K17086 | 3121at3041 | No_hit | 6375at3041 | K01802 |
| 295_2at3041 | K01885 | 3133at3041 | K00025 | 6397at3041 | K02639 |
| 322_1at3041 | K10601 | 3144at3041 | K03843 | 6403at3041 | No_hit |
| 336at3041 | K02355 | 3151at3041 | K20181 | 6407at3041 | K09595 |
| 345_1at3041 | K02303 | 3152at3041 | K00133 | 6423at3041 | No_hit |
| 381at3041 | K25058 | 3173at3041 | K00227 | 6445at3041 | No_hit |
| 388at3041 | K12623 | 3178at3041 | K10258 | 6455at3041 | K03263 |
| 391_1at3041 | K01262 | 3181at3041 | K06215 | 6479at3041 | K06196 |
| 406_1at3041 | K12883 | 3198at3041 | K03036 | 6506at3041 | No_hit |
| 406at3041 | K00615 | 3232at3041 | K03251 | 6516at3041 | K01519 |
| 412at3041 | K00234 | 3261at3041 | K00981 | 6517at3041 | K14831 |
| 415at3041 | K02871 | 3291at3041 | K14820 | 6577at3041 | K00761 |
| 425at3041 | K14950 | 3304at3041 | K24940 | 6621at3041 | K07575 |
| 427_1at3041 | K03526 | 3312at3041 | K12606 | 6622at3041 | K13093 |
| 427at3041 | K15918 | 3315at3041 | K03372 | 6628at3041 | K14560 |
| 439at3041 | K02726 | 3333at3041 | K11996 | 6653at3041 | K08681 |
| 477_1at3041 | K10843 | 3337at3041 | K03035 | 6673at3041 | K01462 |
| 477_2at3041 | K18655 | 3350at3041 | K07513 | 6681at3041 | K03124 |
| 505at3041 | K02145 | 3355at3041 | K20184 | 6682at3041 | No_hit |
| 510at3041 | K15042 | 3370at3041 | K03639 | 6689at3041 | K08246 |
| 550_1at3041 | K06685 | 3371at3041 | No_hit | 6717at3041 | K02372 |
| 550at3041 | K01652 | 3391at3041 | K00856 | 6718at3041 | K20302 |
| 561at3041 | K02727 | 3396at3041 | No_hit | 6721at3041 | K12877 |
| 570_1at3041 | K20791 | 3398at3041 | K15029 | 6733at3041 | No_hit |
| 570_3at3041 | K03032 | 3418at3041 | No_hit | 6734at3041 | K08272 |
| 581at3041 | K01858 | 3428at3041 | K09013 | 6738at3041 | No_hit |
| 599at3041 | K13679 | 3446at3041 | K02906 | 6836at3041 | K10578 |
| 619at3041 | K03456 | 3454at3041 | K03262 | 6837at3041 | No_hit |
| 629at3041 | K03031 | 3456at3041 | K15103 | 6847at3041 | K08343 |
| 635at3041 | K10908 | 3483at3041 | K14964 | 6850at3041 | No_hit |
| 639_2at3041 | K12735 | 3509at3041 | K17605 | 6851at3041 | No_hit |
| 676at3041 | K25639 | 3537at3041 | K12795 | 6861at3041 | No_hit |
| 681_1at3041 | K19306 | 3540at3041 | K03665 | 6880at3041 | No_hit |
| 681_2at3041 | K10956 | 3554at3041 | K03941 | 6885at3041 | K00894 |
| 681at3041 | K03124 | 3556at3041 | K05542 | 6891at3041 | K07023 |

| | | | | | |
|---|---|---|---|---|---|
| 714at3041 | K10590 | 3559at3041 | K03264 | 6896at3041 | K01855 |
| 721_1at3041 | K03943 | 3583at3041 | K00648 | 6905at3041 | K10839 |
| 721_2at3041 | K03237 | 3588at3041 | K14191 | 6909at3041 | K02876 |
| 721_3at3041 | K12877 | 3613at3041 | K06961 | 6930at3041 | K10047 |
| 721_4at3041 | K03942 | 3614at3041 | K00609 | 6952at3041 | K07442 |
| 721at3041 | K03943 | 3618at3041 | K00413 | 6962at3041 | K10575 |
| 731at3041 | K01880 | 3621at3041 | K01807 | 6972at3041 | No_hit |
| 733_1at3041 | K08054 | 3678at3041 | K01764 | 6989at3041 | K05309 |
| 761at3041 | K01887 | 3697at3041 | K02737 | 7007at3041 | No_hit |
| 763_2at3041 | K06965 | 3709at3041 | K01695 | 7011at3041 | K03217 |
| 766at3041 | K09834 | 3721at3041 | K03943 | 7013at3041 | K22074 |
| 779at3041 | K01687 | 3733at3041 | K08054 | 7040at3041 | K04505 |
| 781at3041 | K00392 | 3774at3041 | K03349 | 7053at3041 | K00670 |
| 782_1at3041 | K03979 | 3802at3041 | K04799 | 7062at3041 | K01087 |
| 796at3041 | K07942 | 3806at3041 | K13102 | 7064at3041 | K11885 |
| 803at3041 | K01889 | 3820at3041 | K14799 | 7067at3041 | K15456 |
| 815_1_2at3041 | K00611 | 3835at3041 | K14563 | 7070at3041 | K07297 |
| 818_2at3041 | K15451 | 3854at3041 | K03844 | 7072at3041 | K00943 |
| 832at3041 | K02147 | 3864at3041 | K13617 | 7101at3041 | K02872 |
| 863at3041 | K03246 | 3869at3041 | K01412 | 7128at3041 | K07955 |
| 872_1at3041 | K09498 | 3870at3041 | K14561 | 7132at3041 | K12403 |
| 872at3041 | No_hit | 3895at3041 | K03679 | 7167at3041 | K14397 |
| 878_2at3041 | K01951 | 3911at3041 | K12872 | 7205at3041 | No_hit |
| 878at3041 | No_hit | 3914at3041 | K02728 | 7224at3041 | K21198 |
| 879_2at3041 | K22763 | 3954at3041 | K14810 | 7248at3041 | K21480 |
| 884_2at3041 | K13111 | 3955at3041 | No_hit | 7268at3041 | K18999 |
| 890at3041 | K11968 | 3988at3041 | K01814 | 7269at3041 | No_hit |
| 891_2at3041 | K11594 | 3994at3041 | K06176 | 7276at3041 | K10704 |
| 900at3041 | K01649 | 4012at3041 | K07877 | 7288at3041 | K20303 |
| 911at3041 | K12872 | 4047at3041 | K03428 | 7294at3041 | K03094 |
| 919at3041 | K01251 | 4050at3041 | K10758 | 7308at3041 | K14769 |
| 928at3041 | K06943 | 4055at3041 | K13119 | 7319at3041 | K02953 |
| 930_1_2at3041 | K10047 | 4063at3041 | K01074 | 7331at3041 | K02879 |
| 943_1at3041 | No_hit | 4072at3041 | K12831 | 7334at3041 | K14850 |
| 943at3041 | K12501 | 4074at3041 | K02906 | 7357at3041 | No_hit |
| 963at3041 | K02144 | 4081at3041 | K14568 | 7388at3041 | K11153 |
| 965at3041 | K15113 | 4083at3041 | K02731 | 7392at3041 | K13115 |
| 969_1at3041 | K13789 | 4084at3041 | No_hit | 7394at3041 | K23562 |
| 988at3041 | K02933 | 4086at3041 | K10436 | 7428at3041 | K17972 |
| 992at3041 | K03061 | 4088at3041 | No_hit | 7448at3041 | No_hit |
| 1005at3041 | K01883 | 4127at3041 | K00215 | 7461at3041 | K02896 |
| 1027at3041 | K00914 | 4146at3041 | K17268 | 7462at3041 | K00759 |
| 1029at3041 | K01940 | 4152at3041 | No_hit | 7512at3041 | K09140 |
| 1032at3041 | K12860 | 4160at3041 | No_hit | 7545at3041 | K18532 |
| 1071at3041 | K01845 | 4188at3041 | K02932 | 7548at3041 | K02955 |
| 1072at3041 | K03250 | 4200at3041 | K18328 | 7564at3041 | No_hit |
| 1084at3041 | K27413 | 4202at3041 | K09503 | 7568at3041 | K03152 |
| 1089at3041 | K01885 | 4203at3041 | K00797 | 7570at3041 | K20791 |
| 1090at3041 | K12176 | 4204at3041 | K22912 | 7579at3041 | No_hit |
| 1093at3041 | No_hit | 4217at3041 | No_hit | 7588at3041 | K03975 |
| 1103at3041 | K03062 | 4230at3041 | K12581 | 7633at3041 | K27318 |
| 1128at3041 | K12736 | 4231at3041 | K14864 | 7642at3041 | K12863 |
| 1143at3041 | K01881 | 4259at3041 | K06062 | 7661at3041 | K02880 |
| 1149at3041 | K01881 | 4262at3041 | K14684 | 7698at3041 | K14012 |
| 1170at3041 | K10389 | 4288at3041 | K00930 | 7716at3041 | K20301 |
| 1176at3041 | K14537 | 4290at3041 | K07904 | 7763at3041 | K01520 |
| 1194at3041 | K01733 | 4292at3041 | K13127 | 7785at3041 | K11827 |
| 1233at3041 | K11808 | 4294at3041 | K14066 | 7823at3041 | K08489 |
| 1248at3041 | K13421 | 4295at3041 | K07199 | 7848at3041 | K25817 |
| 1257at3041 | K03253 | 4299at3041 | K02729 | 7853at3041 | K02694 |
| 1267at3041 | K11718 | 4300at3041 | K00254 | 7856at3041 | K01489 |
| 1271at3041 | K24887 | 4302at3041 | K13137 | 7859at3041 | No_hit |

| | | | | | |
|---|---|---|---|---|---|
| 1278at3041 | K00818 | 4312at3041 | K03768 | 7879at3041 | K22763 |
| 1280at3041 | K13100 | 4317at3041 | K01507 | 7914at3041 | No_hit |
| 1282at3041 | No_hit | 4322at3041 | K10601 | 7922at3041 | K20304 |
| 1301at3041 | K12858 | 4327at3041 | K24104 | 7955at3041 | K25866 |
| 1303at3041 | K00705 | 4331at3041 | K05610 | 7956at3041 | K22943 |
| 1320at3041 | K01919 | 4348at3041 | K14190 | 7965at3041 | K15113 |
| 1348at3041 | K01755 | 4381at3041 | K25058 | 7977at3041 | K18156 |
| 1350at3041 | K00899 | 4405at3041 | K15296 | 7993at3041 | K03015 |
| 1379at3041 | K12196 | 4417at3041 | No_hit | 8001at3041 | K20793 |
| 1385at3041 | K12819 | 4419at3041 | K02863 | 8004at3041 | K15731 |
| 1391at3041 | K01262 | 4427at3041 | K15918 | 8021at3041 | K01922 |
| 1428at3041 | No_hit | 4439at3041 | K02726 | 8052at3041 | No_hit |
| 1459at3041 | K04567 | 4446at3041 | K05925 | 8061at3041 | K02966 |
| 1476at3041 | K14564 | 4458at3041 | K14846 | 8069at3041 | K18588 |
| 1477at3041 | K18655 | 4511at3041 | K08495 | 8106at3041 | No_hit |
| 1479at3041 | K10393 | 4519at3041 | K03439 | 8151at3041 | K10666 |
| 1503at3041 | K13832 | 4530at3041 | K02725 | 8157at3041 | K03680 |
| 1511at3041 | K05917 | 4533_1at3041 | K03259 | 8173at3041 | K20359 |
| 1514at3041 | K02146 | 4544at3041 | No_hit | 8191at3041 | K13345 |
| 1518_1at3041 | K03066 | 4550at3041 | K15227 | 8206at3041 | K12197 |
| 1529at3041 | K01657 | 4553at3041 | K08911 | 8229at3041 | No_hit |
| 1532at3041 | K07198 | 4563at3041 | No_hit | 8231at3041 | K11092 |
| 1535at3041 | K14298 | 4573at3041 | K01918 | 8248at3041 | K06268 |
| 1537at3041 | K01939 | 4606at3041 | K12870 | 8289at3041 | K02939 |
| 1553at3041 | K12402 | 4608at3041 | K01079 | 8310at3041 | K10579 |
| 1574at3041 | K00099 | 4627at3041 | K17497 | 8348at3041 | K15280 |
| 1578at3041 | K03163 | 4630at3041 | No_hit | 8372at3041 | K17892 |
| 1590at3041 | K20298 | 4633at3041 | No_hit | 8373at3041 | No_hit |
| 1593at3041 | K01696 | 4664at3041 | K18551 | 8377at3041 | No_hit |
| 1594at3041 | K09458 | 4665at3041 | K00939 | 8384at3041 | K02838 |
| 1596at3041 | K11843 | 4670at3041 | K02866 | 8406at3041 | K12883 |
| 1597at3041 | K01001 | 4703at3041 | K06997 | 8408at3041 | K12876 |
| 1600at3041 | K17491 | 4708at3041 | K00919 | 8415at3041 | K02871 |
| 1622at3041 | K00855 | 4754at3041 | K08916 | 8438at3041 | K26405 |
| 1627at3041 | K00088 | 4759at3041 | K01358 | 8465at3041 | K03016 |
| 1666at3041 | K14376 | 4763at3041 | No_hit | 8494at3041 | K01057 |
| 1670at3041 | K00121 | 4797at3041 | K02730 | 8500at3041 | K12833 |
| 1683at3041 | K00831 | 4800at3041 | K08516 | 8542at3041 | K26403 |
| 1684at3041 | K24939 | 4808at3041 | K20884 | 8595at3041 | No_hit |
| 1720at3041 | K01900 | 4813at3041 | K02836 | 8618at3041 | K08234 |
| 1722at3041 | K11717 | 4822at3041 | K14574 | 8623at3041 | K23564 |
| 1738at3041 | K03065 | 4825at3041 | K07953 | 8629at3041 | K03031 |
| 1815at3041 | K00611 | 4840at3041 | K02865 | 8641at3041 | No_hit |
| 1825at3041 | K20032 | 4855at3041 | K07874 | 8645at3041 | K07766 |
| 1842at3041 | K03265 | 4864at3041 | K17804 | 8662at3041 | K01309 |
| 1848at3041 | K12847 | 4872at3041 | No_hit | 8718at3041 | K12394 |
| 1863at3041 | K00231 | 4873at3041 | K01658 | 8754at3041 | No_hit |
| 1872at3041 | K11583 | 4884at3041 | K03033 | 8778at3041 | K02910 |
| 1917at3041 | K11584 | 4899at3041 | K26736 | 8793at3041 | K00591 |
| 1922at3041 | K14835 | 4904at3041 | No_hit | 8826at3041 | K05609 |
| 1966at3041 | K00817 | 4933at3041 | K15026 | 8836at3041 | K20362 |
| 1974at3041 | K05356 | 4943at3041 | K12501 | 8859at3041 | K02357 |
| 1990at3041 | K01698 | 4956at3041 | K17618 | 8863at3041 | K05969 |
| 1998at3041 | K08193 | 4962at3041 | No_hit | 8868at3041 | K11096 |
| 2014at3041 | K00382 | 4984at3041 | K00586 | 8869at3041 | K02151 |
| 2020at3041 | K10686 | 4993at3041 | K14859 | 8895at3041 | No_hit |
| 2042at3041 | K17255 | 4997at3041 | No_hit | 8920at3041 | No_hit |
| 2043at3041 | K04040 | 5027at3041 | K24621 | 8925at3041 | K03627 |
| 2044at3041 | K00801 | 5035at3041 | K03027 | 8989at3041 | No_hit |
| 2051at3041 | K01866 | 5047at3041 | K07897 | 8991at3041 | K03145 |
| 2057at3041 | K01749 | 5053at3041 | K08517 | 9034at3041 | K12845 |
| 2073at3041 | K03527 | 5057at3041 | K02993 | 9042at3041 | K12817 |

| | | | | | |
|---|---|---|---|---|---|
| 2088at3041 | K00555 | 5060at3041 | K12836 | 9051at3041 | K04460 |
| 2101at3041 | No_hit | 5073at3041 | K03120 | 9101at3041 | K12861 |
| 2104at3041 | K12827 | 5080at3041 | K01850 | 9152at3041 | K03952 |
| 2112at3041 | K15102 | 5108at3041 | K15430 | 9180at3041 | K02947 |
| 2124at3041 | K03953 | 5123at3041 | K02136 | 9197at3041 | K09647 |
| 2130at3041 | K00052 | 5130at3041 | K01244 | 9261at3041 | K15306 |
| 2134at3041 | K02115 | 5142at3041 | K06276 | 9283at3041 | K01522 |
| 2139at3041 | K01760 | 5213at3041 | K03025 | 9401at3041 | K00685 |
| 2152at3041 | K13412 | 5230at3041 | K02989 | 9423at3041 | K20368 |
| 2159at3041 | K01923 | 5231at3041 | K03754 | 9467at3041 | K02717 |
| 2182at3041 | K07562 | 5262at3041 | K10949 | 9470at3041 | No_hit |
| 2202at3041 | K01012 | 5279at3041 | K07565 | 9478at3041 | No_hit |
| 2228at3041 | K01714 | 5292at3041 | K02736 | 9580at3041 | K12611 |
| 2231at3041 | K12864 | 5305at3041 | K01069 | 9600at3041 | K12948 |
| 2233at3041 | K14004 | 5319at3041 | K13519 | 9631at3041 | K22071 |
| 2281at3041 | K00787 | 5335at3041 | K02995 | 9632at3041 | K03236 |
| 2298at3041 | K00145 | 5338at3041 | No_hit | 9637at3041 | K02898 |
| 2318at3041 | K14684 | 5398at3041 | K00991 | 9651at3041 | No_hit |
| 2358at3041 | K01889 | 5409at3041 | K02735 | 9668at3041 | K17795 |
| 2366at3041 | K07178 | 5429at3041 | K02739 | 9687at3041 | K20352 |
| 2370at3041 | K07561 | 5503at3041 | No_hit | 9693at3041 | No_hit |
| 2390at3041 | K03977 | 5430at3041 | K11884 | 9788at3041 | K02881 |
| 2397at3041 | K04354 | 5531at3041 | K03115 | 9812at3041 | K03626 |
| 2423at3041 | K00671 | 5550at3041 | K06685 | 9901at3041 | K14050 |
| 2455at3041 | K01792 | 5551at3041 | K02699 | 9927at3041 | K13171 |
| 2461at3041 | K01867 | 5554at3041 | K06127 | 9999at3041 | K25079 |
| 2490at3041 | K07748 | 5561at3041 | K02727 | 10034at3041 | K00616 |
| 2496at3041 | K18466 | 5563at3041 | K24750 | 10111at3041 | K08066 |
| 2528at3041 | K02259 | 5568at3041 | K12191 | 10167at3041 | No_hit |
| 2531at3041 | K13917 | 5588at3041 | K07943 | 10175at3041 | K20372 |
| 2547at3041 | K11108 | 5589at3041 | K11600 | 10198at3041 | K14396 |
| 2581at3041 | K15283 | 5590at3041 | K03110 | 10205at3041 | K19995 |
| 2583at3041 | K07399 | 5594at3041 | K00999 | 10253at3041 | No_hit |
| 2618at3041 | K14327 | 5644at3041 | K15340 | 10273at3041 | No_hit |
| 2639at3041 | K12735 | 5645at3041 | No_hit | 10293at3041 | K00417 |
| 2664at3041 | K08287 | 5659at3041 | No_hit | 10388at3041 | K12623 |
| 2687at3041 | K13124 | 5662at3041 | K14566 | 10412at3041 | K23490 |
| 2688at3041 | K01735 | 5677at3041 | K03248 | 10491at3041 | K00621 |
| 2699at3041 | K19269 | 5681at3041 | K19306 | 10545at3041 | K02871 |
| 2720at3041 | K07179 | 5701at3041 | K00655 | 10744at3041 | No_hit |
| 2721at3041 | K03237 | 5744at3041 | K18477 | 10873at3041 | K15448 |
| 2728at3041 | K25877 | 5766at3041 | K09834 | 11022at3041 | K18749 |
| 2729at3041 | K14809 | 5785at3041 | K14015 | 11023at3041 | K12871 |
| 2743at3041 | K00606 | 5786at3041 | K01094 | 11078at3041 | K05759 |
| 2749at3041 | K11518 | 5788at3041 | No_hit | 11227at3041 | No_hit |
| 2782at3041 | K03979 | 5794at3041 | K08486 | 11322at3041 | K09549 |
| 2802at3041 | K19073 | 5818at3041 | K15451 | 11392at3041 | K01527 |
| 2820at3041 | K15849 | 5835at3041 | K19828 | 11502at3041 | No_hit |
| 2831at3041 | K01634 | 5839at3041 | K03661 | 11515at3041 | No_hit |
| 2843at3041 | K24752 | 5845at3041 | K12843 | 11518at3041 | K23953 |
| 2851at3041 | K03030 | 5886at3041 | K14016 | 11632at3041 | K02899 |
| 2863at3041 | K03246 | 5905at3041 | K01823 | 11693at3041 | K22746 |
| 2871at3041 | K12479 | 5929at3041 | No_hit | 11703at3041 | No_hit |
| 2878at3041 | No_hit | 5934at3041 | K11755 | 11750at3041 | K03105 |
| 2892at3041 | K17907 | 5988at3041 | K02933 | 11776at3041 | K11086 |
| 2896at3041 | K18649 | 6003at3041 | K02926 | 11870at3041 | K02109 |
| 2932at3041 | K00208 | 6013at3041 | K14696 | 12103at3041 | K03123 |
| 2940at3041 | K05863 | 6065at3041 | K12826 | 12131at3041 | K02942 |
